# Modeling quantifies in vivo neutralization, Fc-mediated killing, and resistance in human clinical trials of five anti-HIV broadly neutralizing antibodies

**DOI:** 10.64898/2026.06.13.732071

**Authors:** Lucero Rodriguez Rodriguez, Jonah Hall, E Fabian Cardozo-Ojeda, Rebecca M Lynch, Alan S Perelson, Joshua T Schiffer, Daniel B Reeves

## Abstract

Broadly neutralizing antibodies (bnAbs) are a promising intervention for HIV prevention, therapy, and cure. bnAb optimization requires precise quantification of *in vivo* functions, many of which cannot be directly measured in humans. We therefore performed a mathematical modeling meta-analysis which integrated four clinical trials and reproduced serial bnAb concentrations, viral loads, and bnAb sensitivities (IC50) in 43 viremic trial participants who received an infusion of VRC01, VRC01LS, VRC07-523LS, 3BNC117 or 10-1074. We compared >300 mathematical models for their ability to recapitulate multi-strain HIV dynamics following bnAb infusion. For each bnAb, our best model identified a scaling factor of 36-462 to pro*j*ect *in vivo* activity from *in vitro* IC50, quantified Fc-mediated infected cell killing in humans over time, and pro*j*ected the timing of bnAb-resistant strain emergence. Using this holistic profile, VRC07-523-LS was generally optimal.

## Introduction

Monoclonal antibodies (mAbs) are approved to prevent or treat respiratory syncytial virus (RSV)^1^, Ebola^2^, rabies^3^, and hepatitis B^4^. Multiple mAbs received emergency use authorizations during the COVID-19 pandemic, though most are no longer effective due to variant escape^5^. Ibalizumab is approved for HIV-1 treatment in some antiretroviral therapy (ART)-resistant cases^6^ and mAb development for HIV-1 remains a vigorous research space.

Broadly neutralizing antibodies (bnAbs) are special mAbs isolated from people with chronic HIV^7,8^, which neutralize a vast range of globally diverse HIV viruses^9^. bnAbs are well tolerated and have been engineered to have long plasma half-lives^10,11^, facilitating weekly or monthly dosing^12–14^. Though long-acting ART limits the need for therapeutic bnAbs at present^15^, vectorized delivery of bnAbs could enable HIV treatment and prevention after a single dose^16,17^.

Prophylactic bnAb trials support the search for an HIV vaccine by providing proof of concept that bnAbs prevent HIV acquisition^18,19^. The antibody mediated prevention (AMP) trials showed modest protective efficacy of the bnAb VRC01 to prevent acquisition of non-resistant HIV^20^ and future trials will include bnAb combinations (or multi-specific bnAbs^21^) to broaden coverage^22–25^.

Relative to ART, bnAbs have immunotherapeutic advantages. Many bnAbs are polyfunctional and can eliminate infected cells. When bnAbs bind infected cells, they can initiate complement or antibody dependent cellular cytotoxicity, ADCC^26,27^ or antibody dependent cellular phagocytosis (ADCP)^28–30^. We collectively refer to these phenomena as *Fc-mediated killing*^30,31^. There is increasing evidence that bnAb binding to viral antigens also stimulates endogenous anti-HIV immune responses including CD8+ T cells^32^ and autologous antibodies^33^. Participants in several trials demonstrated prolonged control of viral load after ART treatment interruption, including some for weeks after bnAb levels mostly disappeared^34–40^. These cases suggest immunological control of viremia, potentially enhanced by bnAbs, and as such, bnAbs are increasingly investigated in HIV cure trials. When combined with latency reversal agents, therapeutic vaccines and strategically timed treatment interruptions of ART, bnAbs may contribute to control or removal of HIV reservoirs that persist despite suppressive ART^41–43^.

An optimal HIV targeting bnAb would exhibit sustained potent neutralization and Fc-mediated killing of infected cells, while also having a high barrier to drug resistance. While assays exist to estimate these metrics *in vitro*^44–46^, it is challenging to quantitate these effects in human trials, particularly as bnAb levels dynamically decay after infusion. bnAb neutralization potency has been quantified with *in vitro* TZM-bl assays, which use reporter cells to estimate inhibitory concentrations required to block pseudovirus infection (e.g., 50% inhibitory concentration, IC50)^47^. For HIV bnAbs tested for prophylactic efficacy, these assays significantly overestimate *in vivo* activity (∼600 fold)^48,49^. Fc-mediated killing is also difficult to measure^29,45^ and improved *in vitro* ADCC does not always correlate with *in vivo* activity^27,44^. Finally minor HIV variants that are resistant to bnAbs are frequently below detection thresholds prior to treatment^50–52^.

Here we build on ours and other mathematical models of HIV bnAbs^48,53–58^ to recapitulate detailed pharmacokinetic (PK), viral load and drug resistance data from 57 viremic clinical trial participants who received one of five different bnAbs (VRC01, VRC01LS, VRC07-523LS, 3BNC117, and 10-1074). We then provide a platform to rank bnAbs according to *in vivo* neutralization, Fc-mediated killing, and drug resistance emergence.

## Results

### HIV bnAb clinical trial data

We collected data from three phase 1 clinical trials testing five bnAbs conducted at the NIH Vaccine Research Center and Rockefeller University Hospital (**Fig. 1a**)^59–62^. In these trials, the bnAbs VRC01, VRC01-LS, VRC07-523-LS, 3BNC117, and 10-1074 were infused in PWH with unsuppressed HIV RNA viremia. 10-1074 targets the V3 loop of HIV envelope (*env*) while all the others target the HIV *env* CD4 binding site.

**Figure 1.**
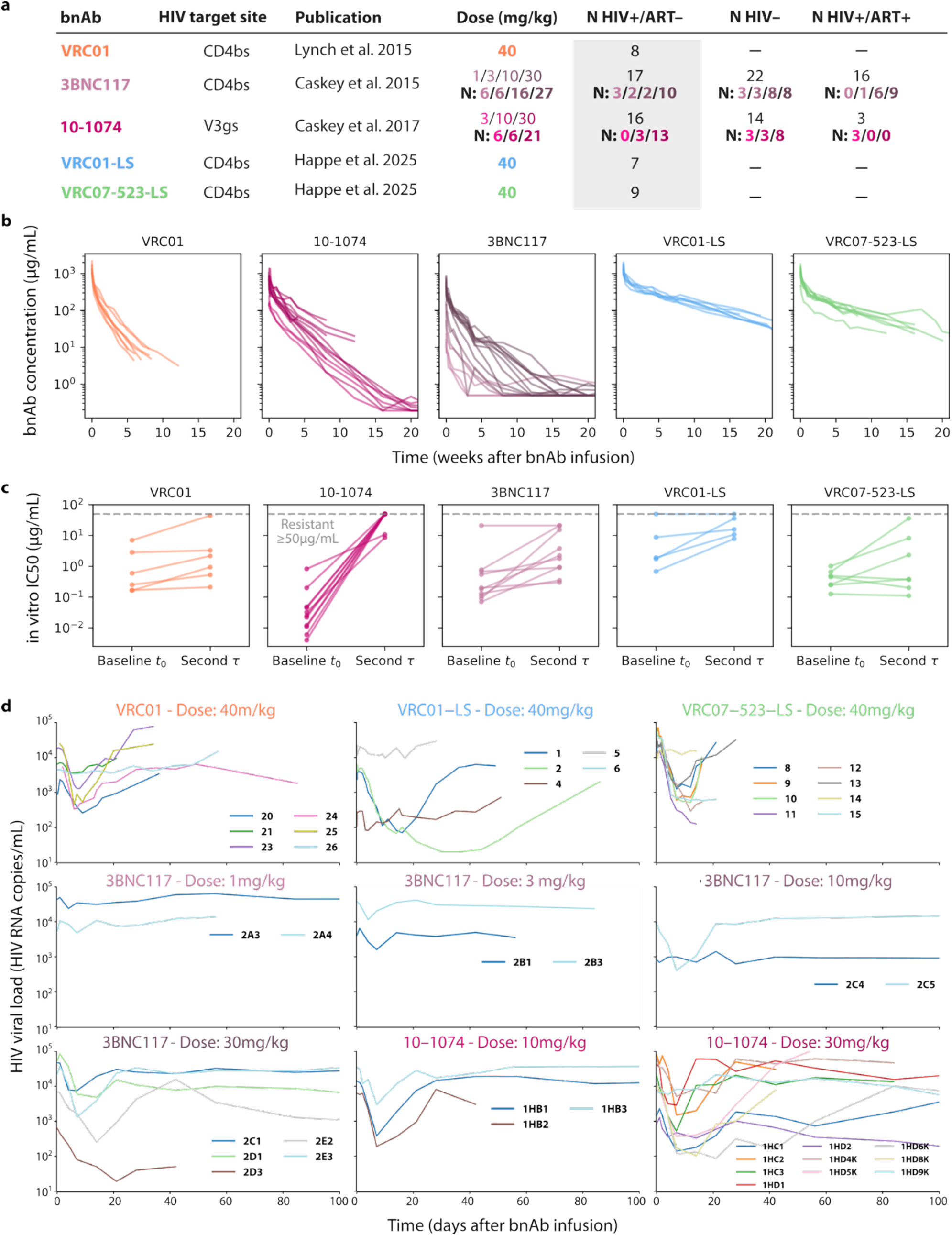
Pharmacokinetic (PK), pharmacodynamic (PD) and viral dynamic (VD) measurements from PWH receiving five HIV bnAbs. **a**, Study summary noting bnAb, HIV target site, original publication, and dose. In total, 112 individuals were infused with an HIV bnAb. Dose column indicates participant number for each dose. The shaded column is PWH who were not on ART (N=57). Studies of 3BNC117 and 10-1074 also included people without HIV (HIV-; N=36), and PWH on ART (HIV+/ART+; N=19). CD4bs: CD4-binding site on gp120; V3gs: V3-glycan supersite on gp120. Different colors correspond to specific bnAb types, and color shading indicates the administered dose. **b**, PK data for each bnAb: different colors correspond with different doses in (**a**). **c**, PD data for all participants who had two sequential measurements of sensitivity to circulating virus (quantified by geometric mean IC50) taken at baseline and a second time, τ. Second time point values were 4 weeks for all 10-1074 and 3BNC117 study participants, 4 (or 5 in one participant each) weeks for VRC01 and VRC01-LS, and 2 weeks for VRC07-523-LS participants. **d**, Viral dynamic data in PWH not on ART. Separate panels are shown for bnAb and dose.

A total of 112 participants were enrolled in these clinical trials, 57 of whom were not on ART at time of bnAb infusion and had detectable plasma viremia (**Fig. 1a**). The 3BNC117 and 10-1074 studies included arms testing bnAb safety and pharmacokinetics (PK) in HIV seronegative individuals (n=36) and HIV positive individuals on ART with suppressed viremia (n=19). There were variable dose arms for 3BNC117, and 10-1074 (doses and numbers of participants in each arm indicated in **Fig. 1a**).

### Inclusion criteria of data for mathematical modeling

We performed mathematical modeling of 43 participants based on adequate availability of three types of longitudinal data. Pharmacokinetic (PK) data consisted of longitudinal plasma bnAb concentrations (**Fig. 1b**). Participants were included if they had PK data for >15 days. bnAb levels generally exhibited biphasic decay patterns in the weeks following infusion (**Fig. 1b**).

Pharmacodynamic (PD) data included the sensitivity of circulating viruses to the infused bnAb quantified as geometric mean 50% inhibitory concentration using the TZM-bl assay (hereafter, *in vitro* IC50) obtained at serial timepoints (**Fig. 1c**). Participants were included if they had two measurements of bnAb IC50 at baseline and a second time point after infusion (all participant information in **Supplementary Data 1**). Second time point values were 4 weeks for all 10-1074 and 3BNC117 study participants, 4 (or 5 in one participant each) weeks for VRC01 and VRC01-LS, and 2 weeks for VRC07-523-LS participants. In some participants, baseline IC50 values were already high, indicating full resistance (**Fig. 1c**). For 10-1074, IC50 values increased substantially between baseline and post infusion whereas less dramatic and more variable increases occurred for other bnAbs. Notably, 10-1074 binds the V3-glycan supersite on gp120 versus the other bnAbs which bind the CD4 binding site.

Viral dynamic (VD) data consisted of serial plasma viral loads (HIV-1 RNA copies/mL) after bnAb infusion (**Fig. 1d**).

### Viral dynamic summary measures during treatment with five different bnAbs

Viral tra*j*ectories were notable for several patterns including no response (lack of or trivial viral decline), sharp viral decay followed by rapid re-expansion, and more prolonged viral suppression followed by slower re-expansion in viral load (**Fig. 1d**). To quantify these differences, we calculated multiple summary measures (see **Fig. 2, Extended Data Fig. 1, Supplementary Table 1**) including maximal viral load reduction (viral load drop), viral load downslope, timing of viral re-expansion, viral load upslope after re-expansion, and time to return to baseline viral load (**Fig. 2a**). Maximal viral load reduction (**Fig. 2b**) and downslope (**Fig. 2c**) differed across bnAb and by dose. VRC07-523-LS 40 mg/kg was associated with the largest viral load reduction and most rapid downslope. 10-1074 at 10 mg/kg (the lower of 2 doses) had the second largest viral load reduction. There was considerable variability within each group, particularly for VRC01-LS, VRC01 and 10-1074, suggesting that varying viral sensitivity might explain these results. Low doses of 3BNC117 (≤3 mg/kg) were associated with minimal HIV plasma RNA reductions (**Fig. 2b**).

**Figure 2.**
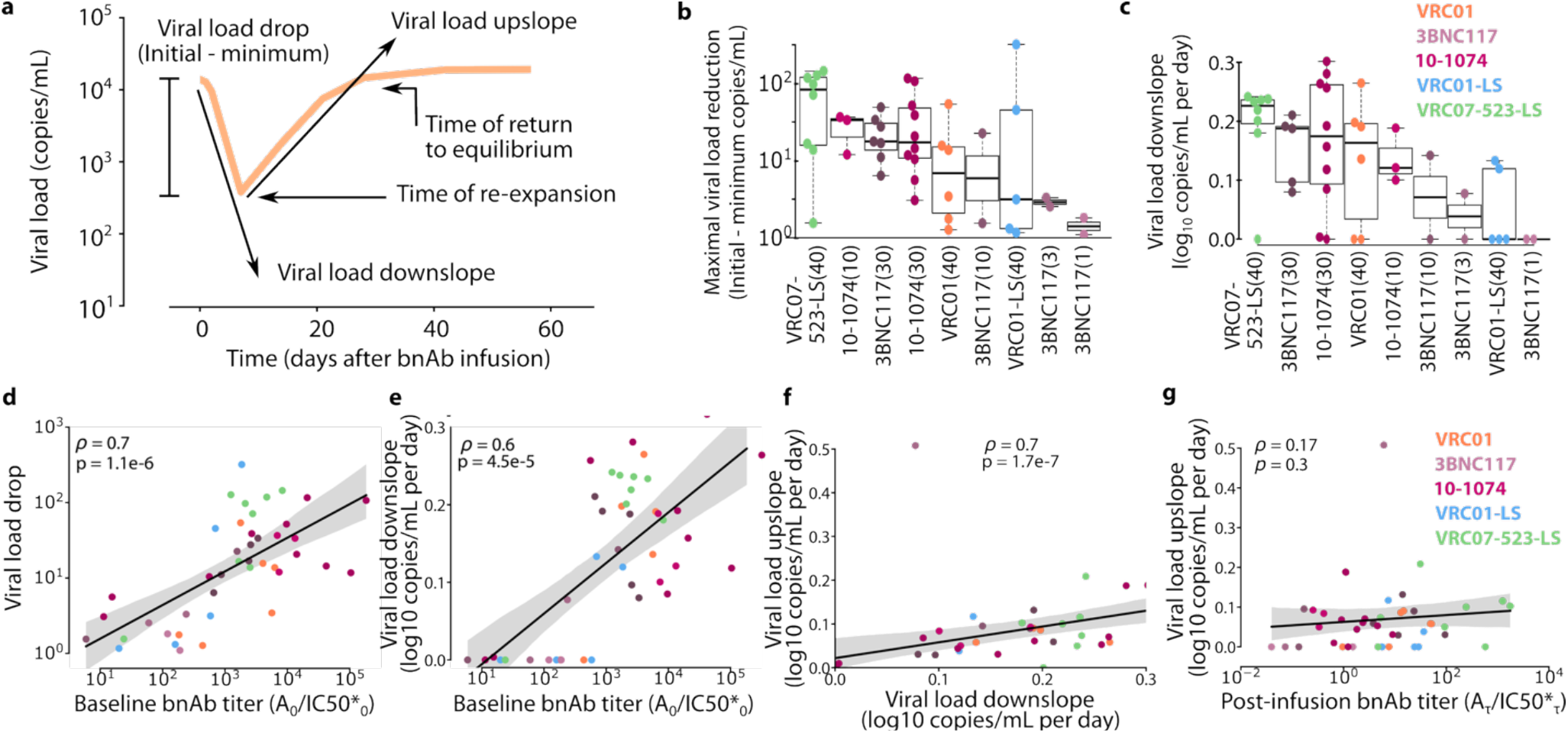
bnAb titer relates to viral load dynamics. **a**, Viral dynamic summary measures (see **Methods** for calculation details). **b**, Viral load drop for each bnAb and dose ranked from most to least **c**, Viral load downslope for each bnAb and dose ranked from fastest to slowest. Boxes indicate median and interquartile range (IQR) and whiskers indicate 1.5x IQR. **d-g**, Correlation analyses between viral dynamic summary measures and bnAb properties. IC50* denotes in vitro with subscript for time point. Marker color denotes bnAb and dose, solid line and shading indicates linear regression and 95% CI, respectively. Inset values denote Spearman correlation coefficient (ρ) and p-value. Titer is the ratio of bnAb concentration to in vitro IC50: **d**, Baseline bnAb titer vs viral load drop and **e**, Baseline bnAb titer vs viral load downslope. **f**, Viral load downslope vs upslope. **g**, Second time point in vitro IC50 vs viral load upslope.

### Baseline bnAb titers predict therapeutic responses and viral load drop

To determine how bnAb levels and virus sensitivity associated with viral dynamic summary measures, we computed baseline bnAb titer (initial bnAb concentration divided by *in vitro* IC50*). There was a strong positive correlation between titer and viral load drop (Spearman rho=0.7, **Fig. 2d**). bnAb titer was also correlated with observed downslope after bnAb infusion (p=4e-5, **Fig. 2e**). However, this association is harder to interpret due to the difficulty of estimating a downslope in weak responders. Only assessing individuals with therapeutic responses (>0.5 log10 drop), downslope was not associated with titer (p=0.8, data not shown). Most importantly, all individuals with titers above 1000 had therapeutic responses. These results imply titer predicts a therapeutic response, and that viral load drop increases with increasing titer.

There was strong correlation (rho=0.7, p=2e-7) between the HIV RNA upslope and downslope with upslope ∼2/3 slower than the downslope (**Fig. 2f**). Therefore, particularly rapid decay as seen in **Fig. 1d** was not predictive of more sustained lowering of HIV RNA. There was also no strong correlation (p=0.2) between second time point virus *in vitro* IC50 divided by concentration at that time point and the viral load upslope (**Fig. 2g**), indicating that a waning bnAb effect, whether by decreased concentration and/or more resistant variant emergence does not necessarily result in faster rebound.

### Pharmacokinetic (PK) modeling meta-analysis refines bnAb half-lives in people with and without HIV

PK modeling was performed for each bnAb during original phase 1 studies^63–65^. To harmonize modeling approaches across studies, we developed an optimal model for all study data simultaneously. We used a population nonlinear mixed effects modeling (pNLME) approach to fit a two compartment PK model to all observed bnAb levels across trials (**Methods, Fig. 3a, Supplementary Table 2**). The model required a covariate for each bnAb and provided an excellent fit, closely reproducing biphasic kinetics across all doses and bnAbs (**Fig. 3b, Extended Data Fig. 2, 3**, & **4, Supplementary Table 3**, & **Extended Data 2-Table 1**). Median half-lives were: ∼39 and ∼33 days for VRC01-LS and VRC07-523-LS, respectively, compared to ∼10 days for VRC01 and 3BNC117 and ∼13 days for 10-1074 (**Fig. 3c, Table 1**). These estimates sometimes differed from original reported values of 15 days for VRC01^63^, 17 days for 3BNC117^59^, 24 days for 10-1074^61^, 71 days for VRC01-LS^64^, and 38 days for VRC07-523-LS^65^, which may reflect differences in our combination model compared to each original separate bnAb model structure.

**Figure 3.**
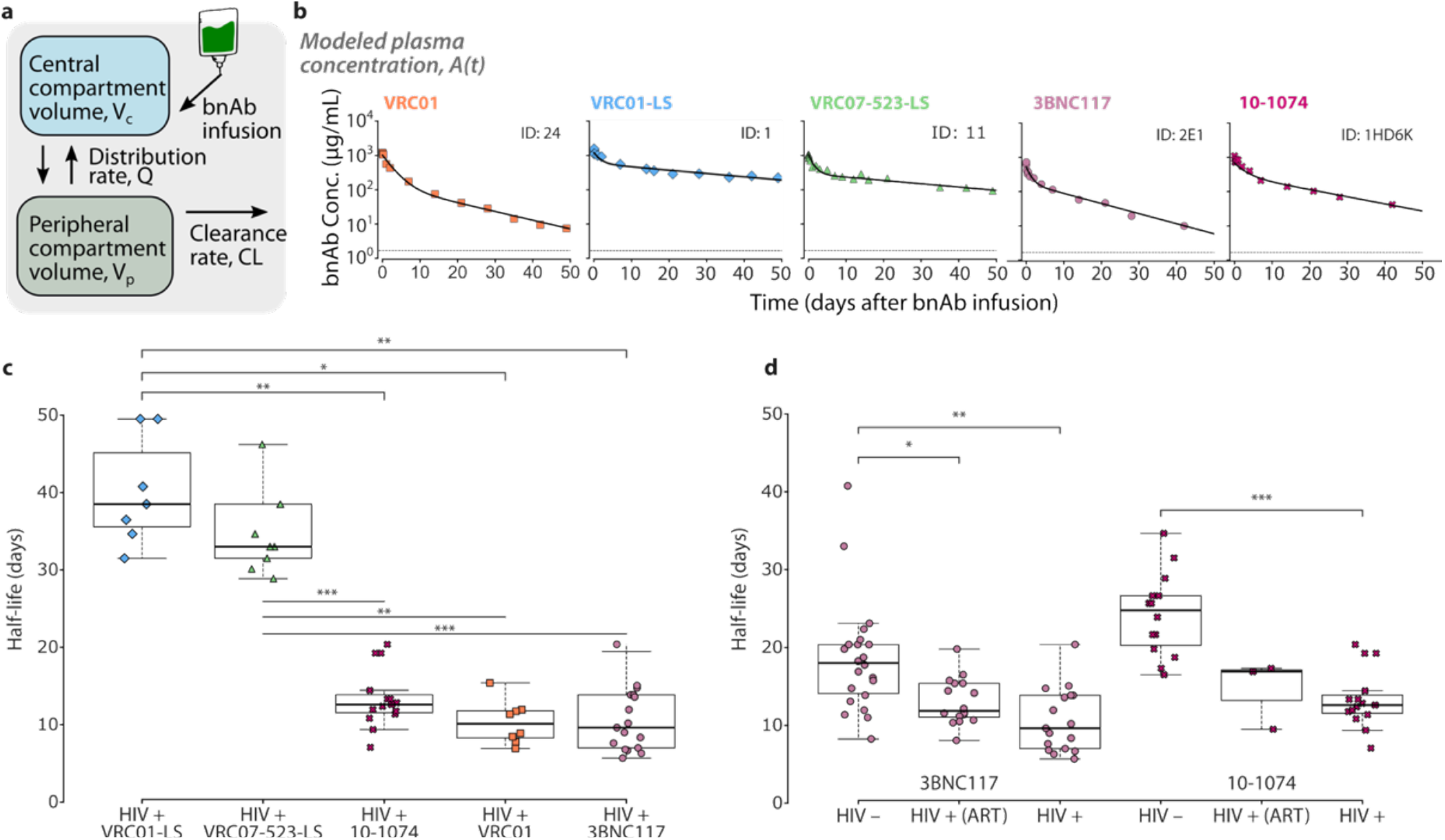
Combining trial data for accurate PK model of bnAb levels. **a**, Cartoon schematic for the two compartment PK model. **b**, Representative PK model fits (line) to observed bnAb concentration data for one person per bnAb (PWH off ART, PWH on ART, and HIV negative participant data and model fits in **Extended Data Fig. 2-4**, respectively). **c**, Model estimated half-lives for each bnAb in PWH off ART. **d**, Model estimated half-lives for each 3BNC117 and 10-1074 in PWH on and off ART, and in HIV negative individuals. Markers indicate individual participant values. Boxes indicate median and interquartile range (IQR) and whiskers indicate 1.5x IQR. Bonferroni corrected statistical comparisons denoted by *: p≤0.05, **: p≤0.01 ***: p≤0.001.

Both HIV infection and ART treatment impacted 3BNC117 and 10-1074 half-life. bnAb half-life in HIV negative individuals was a median of ∼18 days for 3BNC117, and ∼24 days for 10-1074. PWH who received 3BNC117 and 10-1074 while on ART had a median half-life of ∼12 and ∼17 days, respectively (**Fig. 3d**). Lastly, dose and initial bnAb concentration were not correlated with half-life (p=0.3).

### Modeling multi-strain HIV kinetics during bnAb treatment reveals in vivo neutralization potency, Fc-mediated killing, and resistance emergence

The complex correlations between bnAb titer and viral load upslope and downslope (**Fig. 2e&f**) imply that simple dose-response relationships do not fully explain individual virologic outcomes. There are many plausible, non-mutually exclusive and time varying drivers of viral load kinetics. For instance, bnAb neutralization likely decreases over time as bnAb level wane, as does any Fc-mediated killing. Resistant variant emergence likely depends on pre-existing frequencies of minor variants. Because these functions interact in a complex dynamical system, we sought to use mathematical modeling to determine necessary mechanisms in this system and then elucidate the relative impact of each mechanism on observed HIV RNA kinetics.

We developed eight competing and mechanistically distinct differential equation based mathematical models and assessed 350 versions of these models varying their statistical properties to optimize parameter estimate robustness and covariate structure. Using the optimal statistical formulation of each mechanistic model, we then ran 10 or more replicate assessments to ensure reproducibility of results (**Methods**). Using pNLME, corrected Bayesian Information Criteria (BICc) and constraints imposed by literature review, we selected the optimal model (**Supplementary Table 4**, details for statistical optimization of each mechanistic model in **Supplementary Data 2-Table 2**).

We achieved close model fits to drug levels (**Fig. 3, Extended Data Fig. 2-4**) and total viral load, denoted by *VL* = *V*_1_+*V*_2_, (**Fig. 4c, Supplementary Data 1**). We also evaluated whether at the second time point, the strain levels agreed with experimental observations of IC50. We defined the ratio of IC50s from second time point to baseline as the resistance factor *ρ*, such that *ρ*>1 implies rebound with more resistant virus. Then, we score models to ensure *V*_2_>*V*_1_ at the second time point for any individual observed to have *ρ*>1. Our best model’s predicted viral strain behavior was accurate, with only 1 of 43 individuals having an incorrect prediction (ID 20), whose experimental measure was *ρ*∼2 with both IC50s < 1 µg/mL, or a relatively similar IC50 between baseline and second time point viral variants.

**Figure 4.**
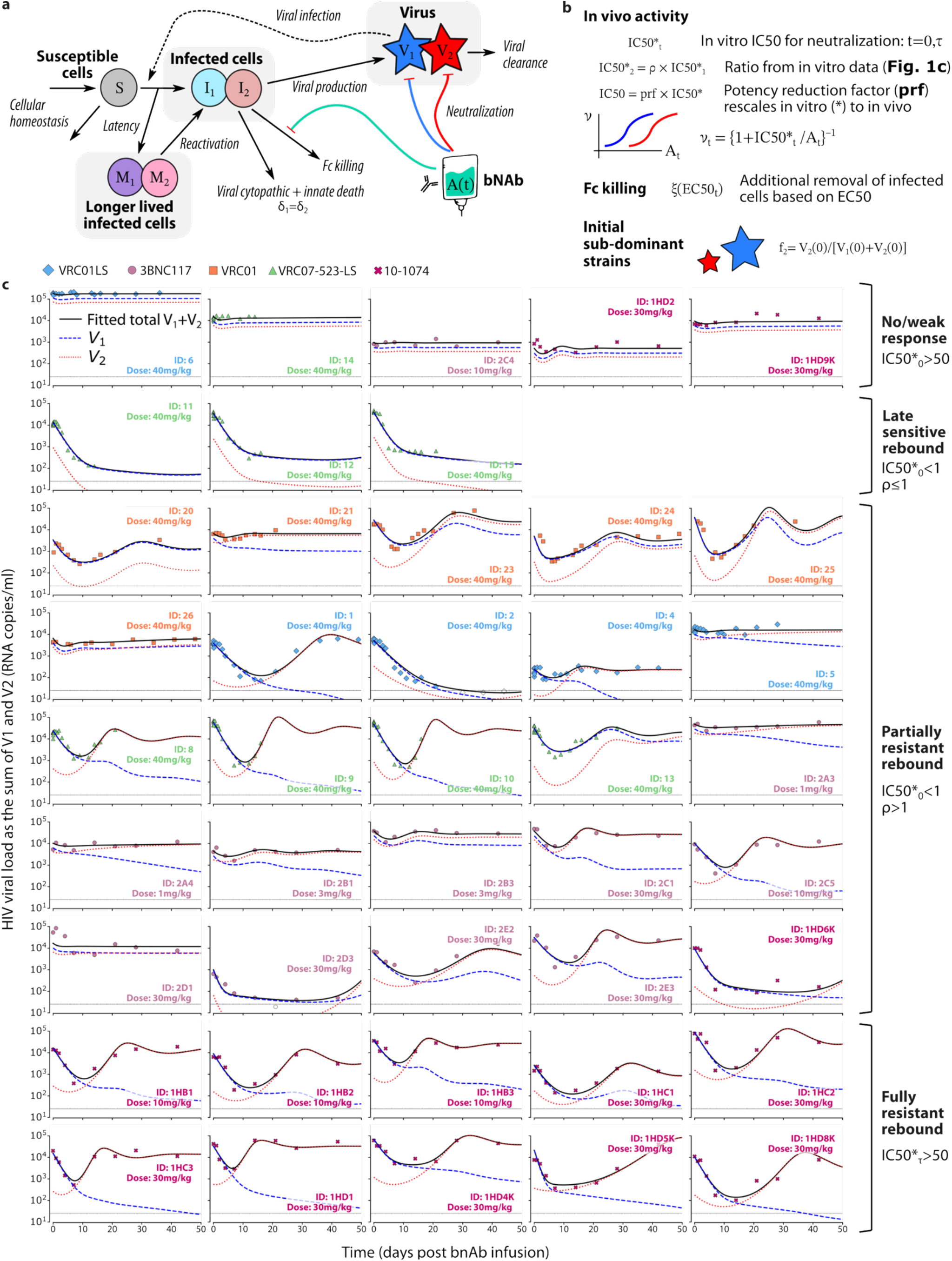
Mathematical model recapitulates viral kinetics after bnAb therapy during active HIV infection. **a**, Best mechanistic model of HIV infection dynamics after comprehensive model selection. The model tracks susceptible cells (S), actively virus-producing infected cells (I_j_), non-virus-producing longer-lived infected cells that can become active (M_j_), and viremia (V_j_). There are two viral strains to allow resistant variant emergence, j = {1,2}. Through the functions υ_1,2_, the model incorporates bnAb-mediated neutralization, Fc-mediated killing of infected cells that depends on bnAb concentration with EC50, and the emergence of initially sub-dominant variants (initial fraction f_2_) with potentially higher IC50 and EC50 for V_1_ than V_2_. **b**, Relationship of in vitro to in vivo IC50 in the model defined as the potency reduction factor. **c**, Model captures diverse kinetics of viral load decay and rebound following bnAb infusion in 43 PWH across clinical trials. Markers denote viral load data, shape and color indicate bnAb. Black line denotes the model fit (VL = V_1_+V_2_), while the blue and red dashed lines denote the two types of viral strains, V_1_ and V_2_ respectively. Panels are grouped by viral kinetic phenotype: non-responders, partially resistant rebounders, fully resistant rebounders and late sensitive rebounders (**Methods**).

All models included susceptible CD4+ T-cells and short-lived infected cells which produce HIV RNA. HIV RNA was assumed to infect cells dependent on viral load and target cell levels and to be cleared at a constant rate. Each model incorporated our PK model and assumed that bnAbs neutralize HIV in a concentration dependent fashion. We tested models with and without 1) Fc-mediated killing of infected cells, 2) pre-existing sub-dominant strains with partial or complete bnAb resistance, and 3) long-lived infected cells. To model viral load setpoint before therapy, we ensured (via analytical calculations, **Methods**) that each individual’s initial conditions were at a steady state self-consistent with estimated rates. **Fig. 4a** illustrates our final compartmental model after model selection (see **Methods** and **Supplementary Table 5 & Supplementary Data 2-Table 3**).

This best selected model provides mechanistic insights into the processes governing therapeutic responses (**Fig. 4b**). First, to match all data simultaneously, the model required explicit consideration of two strains of virus (*V*_1_ and *V*_2_), allowing for potentially resistant minor variants to pre-exist before therapy. We assumed that the ratio of baseline to second time point *in vitro* IC50 data (the resistance factor *ρ*, data in **Fig. 1c**) explains the relative difference in IC50 of *V*_1_ and *V*_2_. The model did not require ad*j*ustments to viral fitness differences between *V*_1_ and *V*_2_ because viral loads generally rebounded to pre-treatment levels regardless of change in IC50 over time (**Fig. 1d**). But the model required an *in vivo* IC50 for each participant that was generally larger than *in vitro* IC50 values. Models also improved when bnAbs exerted Fc-mediated killing that depended on bnAb levels and saturated above a certain level (the Fc-mediated killing EC50). Model fit improved further when the Fc-mediated killing EC50 was distinct from *in vivo* neutralization IC50 (**Fig. 4b**). Another assumption that improved model fit was inclusion of longer-lived infected cells (as previously shown with ART treatment modeling^67,68^). Together, the best fit model for each participant therefore estimates the bnAb’s *in vivo* IC50, the strength and saturation constant for Fc-killing, and the initial fraction of a potentially resistant minor variant.

### Classification of viral kinetics according to bnAb selection

To examine patterns of bnAb selection, we classified participants into four categories based on absolute values of IC50s (IC50 > 50 µg/mL means resistant) and resistance factor *ρ*. If *ρ*>1, we infer that rebound occurs with less sensitive minor variants sub-detectable before infusion and selected under bnAb pressure. 5 *non or weak responders* had baseline HIV strains that were resistant to the infused bnAb 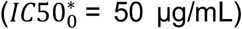. 10 *fully resistant rebounders* (all of whom received 10-1074), were sensitive at baseline but resistant at the second time point 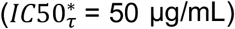. 25 *partially resistant rebounders* had viruses that were not completely resistant 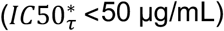 but were *ρ*=16-269 fold more resistant at the second time point. Finally, 3 *late sensitive rebounders* (all of whom received VRC07-523-LS) had *ρ* < 1, the second time point IC50 was similarly or more sensitive than baseline viruses. These participants were not observed to rebound. The model pro*j*ected that viral re-expansion would slowly start more than 50 days after infusion due to waning bnAb concentrations, but we cannot confirm this pro*j*ection. **Fig. 4c** and **Supplementary Data 3** show the model predicted behavior of ma*j*or and minor strains *V*_1_ and *V*_2_. For non-responders, *V*_1_ and *V*_2_ remain parallel. For resistant rebounding virus, *V*_2_ (red line) crosses *V*_1_ (blue line) typically 1-3 weeks after bnAb infusion, indicating strains *V*_2_ sweeping to dominance. In the three non-resistant cases, both strains follow similar decay kinetics though the initial concentration of *V*_2_ is not truly identifiable.

Lastly, our dataset included individuals with missing pre- or post-infusion in vitro IC50 measurements. We fit the model to this subset of individuals using posterior estimates from the full cohort as informative priors (**Supplemental Data 2-Table 3**). The model successfully captured viral load decay kinetics and strain dynamics while simultaneously estimating the missing IC50 values (**Supplementary Fig. 1 & Supplementary Table 6**).

### Quantifying the relationship between *in vivo* and *in vitro* IC50 estimates

To determine bnAb concentrations needed for *in vivo* neutralization and how these levels relate to *in vitro* potency estimates, we contrasted model estimated *in vivo* IC50 to TZM-bl assay measured *in vitro* IC50. We defined the potency reduction factor (prf), which is the fold increase between *in vitro* to *in vivo* IC50 value, effectively a quantitation of how much the *in vitro* assay overestimates *in vivo* potency, and importantly, a value that could pro*j*ect *in vivo* behavior after *in vitro* measurements. Median prf for CD4bs bnAbs were 191, 36, 88, and 115 for VRC01, 3BNC117, VRC01-LS, and VRC07-523-LS, respectively. But median value for 10-1074 was 462, indicating the largest discrepancy between *in vitro* and *in vivo* estimates (**Fig. 5a, Table 1**). There was variability across individuals, suggesting that differences between *in vitro* and *in vivo* neutralization are not wholly constant (**Extended Data Figure 5**), due to many possible host or virus differences. The interquartile range variability was generally within a factor of 2-4 for CD4bs bnAbs were 172-223, 20-114, 52-265, and 65-207, for VRC01, 3BNC117, VRC01-LS, and VRC07-523-LS, respectively. Thus, predictions of future *in vivo* neutralization from *in vitro* measurements could apply prf and prf ranges, or more conservatively, the upper end of these intervals. Again, 10-1074 had the widest range of prf 58-2919, indicative of further discrepancies between *in vitro* and *in vivo* neutralization for V3gs bnAbs.

**Figure 5.**
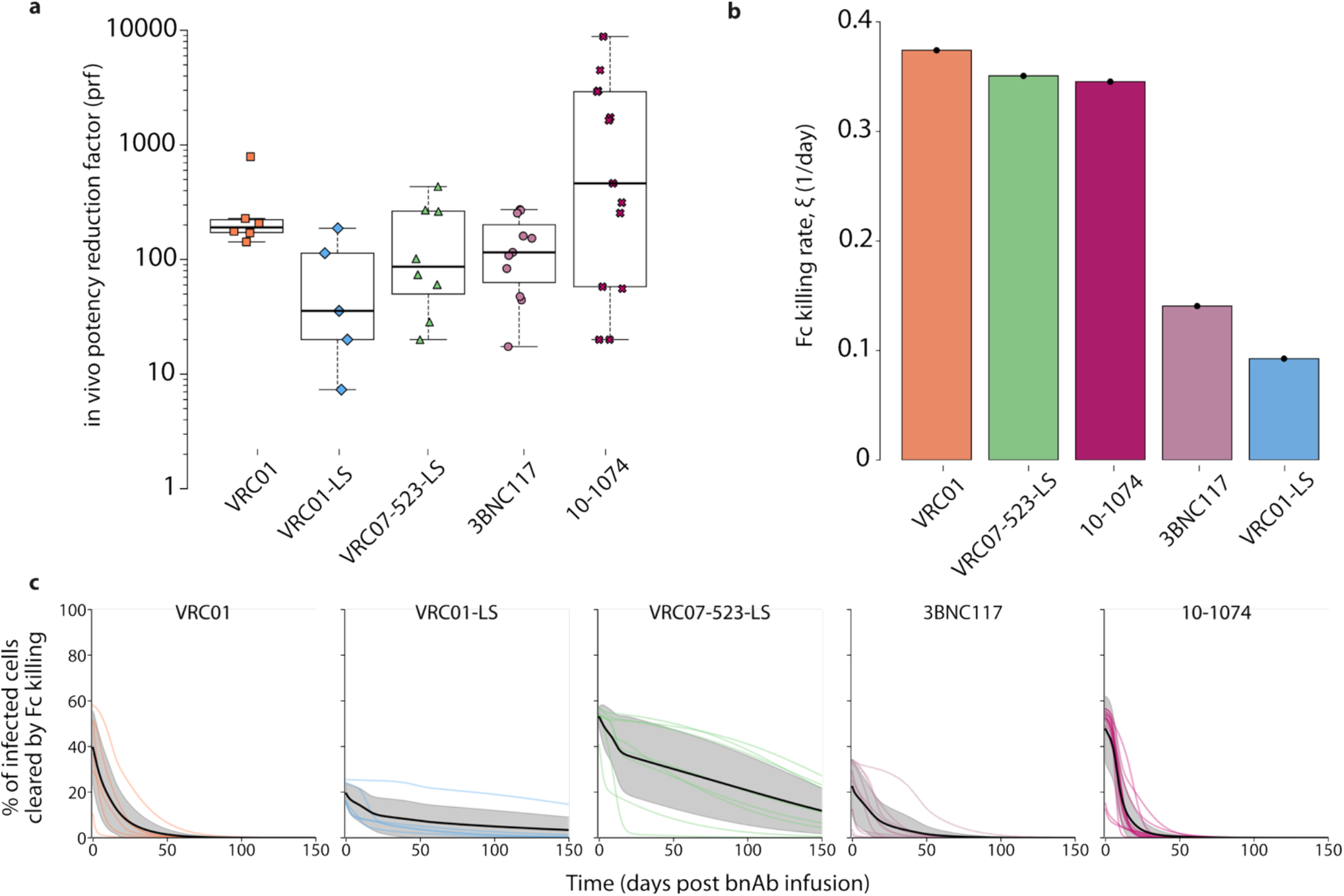
Model estimates of in vivo neutralization and Fc-mediated killing for each bnAb. **a**, Estimates of in vivo potency reduction factor (prf) for each bnAb (dots for each individual). **b**, Population estimates of maximum Fc-mediated killing rate for each bnAb. **c**, Time varying percentage of cells removed by Fc-mediated killing for each bnAb (panel). Each colored line is an individual. Solid black line is mean, shaded gray is the standard deviation interval.

**Table 1.** Summary of model estimated parameters for each bnAb. Half-life indicates long-term plasma clearance. Resistance factor indicates the ratio of final to initial in vitro IC50 from experimental data. In vivo neutralization IC50 and Fc-mediated killing EC50 indicate model estimated plasma concentration (µg/mL) needed to reach 50% activity of the mechanism, respectively. In vivo potency reduction indicates the fold increase of IC50 from experimentally observed in vitro IC50 to model estimated in vivo IC50. Fc killing proportion indicates the model estimated % of infected cell death in the first 30 days following infusion due to Fc-mediated killing.

| Model estimates<br>Median [25-75% range] | VRC01 | VRC01-LS | VRC07-523-LS | 3BNC117 | 10-1074 |
| --- | --- | --- | --- | --- | --- |
| Half-Life (days) | 10 [8–12] | 39 [36–45] | 33 [32–39] | 10 [7–14] | 13 [12–14] |
| Resistance factor (ρ) | 2.8 [1.5–5] | 5.6 [1.8–12] | 1.3 [0.8–8.3] | 6.1 [2.1–20] | 1100 [250–4200] |
| *in vivo* neutralization<br>IC50 | 133 [71–408] | 359 [62–1000] | 45 [29–93] | 19 [10–67] | 33 [12–73] |
| *in vivo* Fc killing EC50 | 317 [131–873] | 359 [62–1000] | 45 [29–93] | 19 [10–67] | 33 [12–73] |
| *in vivo* potency reduction<br>factor (prf) | 191 [172–223] | 36 [20–114] | 88 [52–265] | 115 [65–207] | 462 [58–2920] |

| Fc killing (%) | 13 [8–22] | 12 [10–15] | 38 [35–45] | 9 [6–15] | 10 [4–30] |
| --- | --- | --- | --- | --- | --- |

### Quantifying the strength of Fc-mediated killing of infected cells

To demonstrate the maximal cumulative effect of Fc-mediated killing for each bnAb, we plotted the strength of the antibodies Fc function activation parameter (ξ) (**Fig. 5b**). VRC01, 10-1074, and VRC07-523-LS demonstrated greater maximal cytolytic effects than 3BNC117 and 10-1074 (**Fig. 5b**). VRC07-523-LS demonstrated particularly intense cell killing during the first two weeks and eliminated the highest percentage of infected cells via Fc-mediated mechanisms over 150 days (**Fig. 5c**) by virtue of potency and prolonged half-life. VRC01 and 10-1074 Fc mediated killing rapidly declined during the first 30 days of treatment (**Fig. 5c**).

Our most accurate model estimated that the EC50 for Fc-mediated killing function was lower than the IC50 for neutralization, suggesting that Fc-mediated cell lysis function requires lower bnAb concentrations than neutralization. When we tested a model requiring that the in vivo Fc function EC50 was equal to or greater than the in vivo neutralization IC50, viral load tra*j*ectories were identical, suggesting that the lower limit of Fc killing EC50 is not identifiable. Because experimental evidence suggests that concentration thresholds are lower for neutralization than ADCC or ADCP^44^, we proceeded with the constraint that IC50≤EC50 during model fitting (**Supplementary Data 2-Table 2**).

The model suggests that Fc-mediated killing effects contributed to high absolute number (**Extended Data Fig. 6**) and relative percentages (**Extended Data Fig. 7**) of infected cells killed during the first 2 weeks of VRC01, 10-1074 and VRC07-523-LS therapy when bnAb levels were high. Confidence bounds were wider in later weeks because of variance among trial participants as well as uncertainty about the lower limit of *in vivo* EC50. Moreover, the impact of Fc-mediated effects and possible vaccinal effects are difficult to distinguish at these later stages.

### Verification of Fc-mediated effects modeling in mice and SHIV infected non-human primates who received bnAbs with natural and mutated Fc regions

To further estimate lower constraints on *in* vivo Fc-mediated EC50 levels in humans, we modeled available non-human primate data. Two different studies observed a 20-50% enhancement of HIV/SHIV viral load clearance in humanized mice and rhesus macaques treated with wild type bnAbs versus equivalent bnAbs lacking Fc-function^26,27^. We used a simplified version of our best model to analyze data from the macaque study, finding that the bnAb VRC07-523-LS enhanced the killing rate at similar magnitudes as in the human data. The model reproduced data showing less significant viral load reduction during the first 5 days of therapy in animals treated with either an Fc-deleted or mutated VRC07-523-LS. Our estimates of EC50 for Fc-mediated killing versus IC50 for neutralization also suggest the possibility of more potent Fc-mediated activity *in vivo* (**Supplementary Fig. 2**). This additional analysis did not allow further precise specification of the duration of Fc-mediated lysis more than a month after infusion

### Estimation of pre-existing resistant variant fraction

The initial fraction of the pre-infusion least abundant strain (*V*_2_) was estimated by the model to be a median of 1.5% for the 10-1074 cohort, 4.4% for the VRC07-523-LS cohort, 7.3% for the VRC01 and VRC01-LS cohorts, and 39% for the 3BNC117 cohort (**Supplementary Fig. 3**). The initial abundancy of *V*_2_ was heterogeneous across individuals. Despite a low initial abundance of *V*_2_ pre- and post-infusion, 10-1074 had the most consistent viral escape with fully resistant *V*_2_.

### Holistic ranking of bnAbs via model-estimated in vivo properties

Model output allowed unique comparisons of key characteristics across the five bnAbs. All antibodies had instances of selection for resistant mutants. Overall, we identified that VRC07-523-LS had among the highest *in vivo* neutralization potency (*in vivo* IC50), longest half-life, maximal Fc-mediated cytolytic potency, and highest barrier to resistance (**Table 1**). VRC01 had lower *in vivo* neutralization potency, short half-life, high maximal Fc-mediated cytolytic potency, and a somewhat higher barrier to resistance. VRC01-LS had lower *in vivo* neutralization potency, long half-life, low Fc cytolytic potency, and a relatively higher barrier to resistance. 3BNC117 had high *in vivo* neutralization potency, short half-life, low Fc-mediated cytolytic potency, and a lower barrier to resistance. Finally, 10-1074 had high *in vivo* neutralization potency, short half-life, brief Fc-mediated potency, and a very low barrier to resistance.

### Counterfactual simulations to interrogate mechanistic impact on viral kinetics

We next performed counterfactual simulations of the model to assess the relative effects of neutralization and Fc-mediated killing (**Fig. 6a**), the role of pre-existing resistant variants (**Fig. 6b**) and overestimation of neutralization potency *in vitro* (**Fig. 6c**) on the observed viral kinetics. In each case, we ad*j*usted a model parameter and explored counterfactual changes to viral dynamic metrics (**Fig. 2a**).

**Figure 6.**
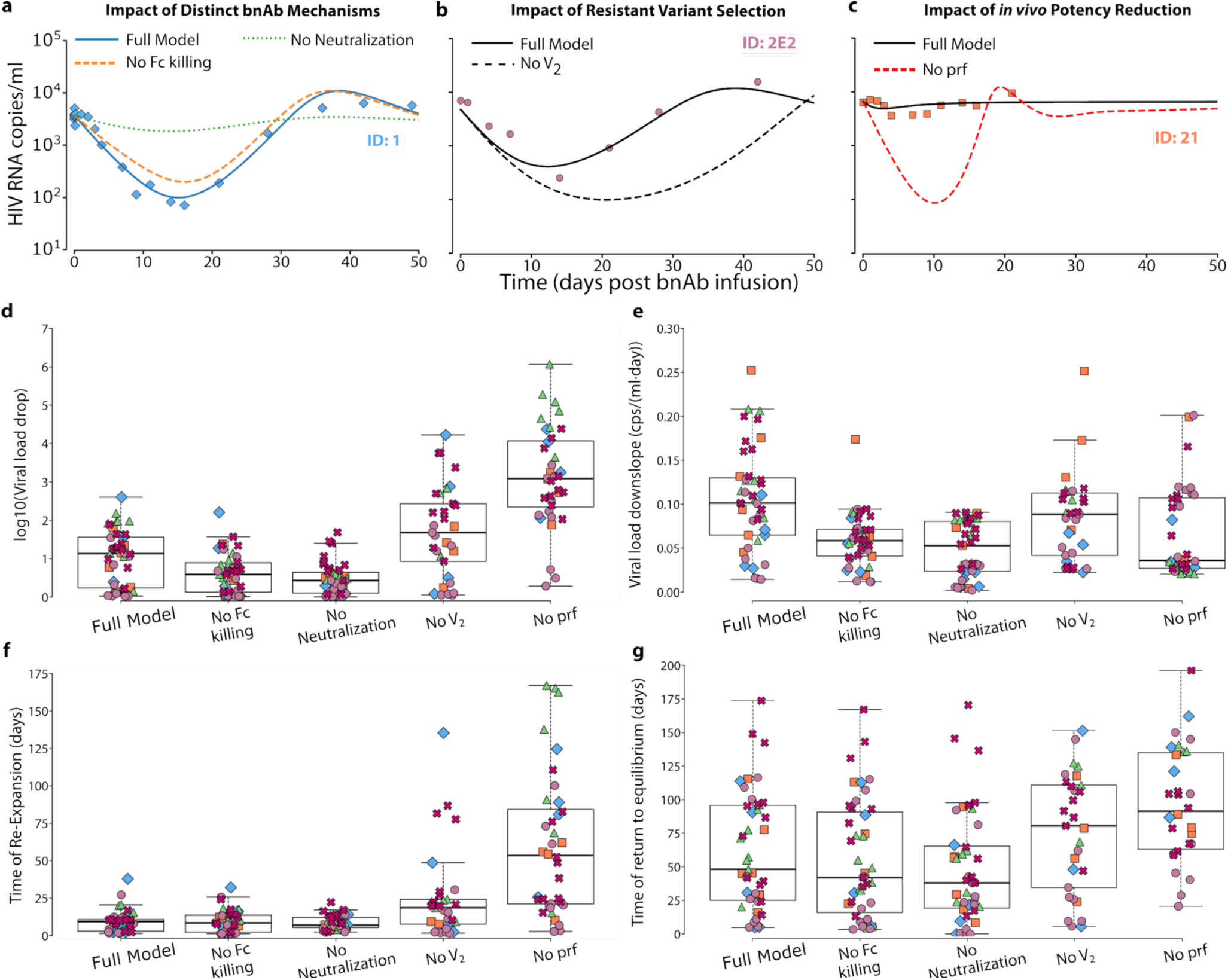
Model simulations of viral load kinetics assuming counterfactual knock out of bnAb mechanisms and/or resistant strains. **a**, Example simulations isolating bnAb neutralization (dashed orange) vs Fc killing (dashed green) compared to the full model (blue, and data markers). **b**, Example simulation removing selected virus (V2, dashed) vs full model (solid, and data markers) **c**, Example simulation removing the difference between in vivo and in vitro efficacy (prf, dashed orange) vs full model (solid, and data markers). **d-g**, Individual values (markers, colored by bnAb) and box plot across all bnAbs for various kinetic measures (including viral load drop, downslope, time of re-expansion, and time of return to equilibrium (set point), see **Fig. 2a** and **Methods** for definitions and calculations). Each box represents a scenario, from the full (best fit) model to counterfactual scenarios no Fc killing, no neutralization, no sub-dominant variants (V_2_), and no in vivo potency reduction.

Neutralization was responsible for a slighter higher proportion of viral load decay than Fc-mediated cell removal though both played important roles (**Fig. 6d**). The median rate of viral decline decreased by approximately 50% when either neutralization or Fc killing was removed from the model respectively (**Fig. 6e, Extended Data Fig. 8**). The median start time of viral re-expansion was consistent with exclusion of each mechanism (**Fig. 6f**). The addition of Fc-mediated effects only prolonged time to resuming viral setpoint slightly (**Fig. 6g**). There was marked variability for each of these metrics across individuals. Participants with sustained suppression over several weeks required Fc killing and neutralization to achieve this effect (**Extended Data Fig. 8**). Overall, our results suggest that neutralization and Fc killing work synergistically to more rapidly lower viral load to a lower nadir, but that time to setpoint is unchanged probably because of rapid rebound (**Fig. 2f**). These results also indicate that resistance decreases neutralization and Fc-function similarly.

To estimate the impact of bnAb resistance on HIV viral load, we performed simulations in which only the original dominant strain was present (**Fig. 6b**; see **Extended Data Fig. 9**). The rate of viral decline increased marginally (**Fig. 6e**) while the extent of viral load reduction increased without selection for resistance (**Fig. 6d**), while the start time of viral re-expansion increased by a median of 10 days (**Fig. 6f**). The simulated time to return to viral load steady state was delayed by a median of ∼32 days absent a resistant virus **(Fig. 6g**).

Our third counterfactual assumed that observed *in vitro* potency estimates were relevant *in vivo* (**Fig. 6c**; see **Extended Data Fig. 10**). The rate of viral decline was significantly lower, perhaps related to artefacts of the initial downslope rapidly re-equilibrating (**Fig. 6e**) and the start time of viral re-expansion significantly increasing (**Fig. 6f**). The viral load reduction increased dramatically by ∼2 logs (**Fig. 6d**) though viral load stabilization was delayed only by a median of ∼43 days likely due to rapid viral rebound (**Fig. 6g**). Some individuals had predicted sustained responses over 50 days (**Extended Data Fig. 10**).

## Discussion

We developed a comprehensive data-validated mathematical model to provide a meta-analysis level summary of five currently available HIV-1 targeting broadly neutralizing antibodies (bnAbs) tested in several clinical trials^59–62^. These intensive trials provided uniquely granular longitudinal viral loads, bnAb concentration, and two time point quantification of bnAb sensitivity for each participant. Despite highly heterogeneous outcomes, by coupling individual bnAb levels with antiviral activity and modeling across the entire population, we were able to meaningfully interpret each participant’s viral load tra*j*ectory. We specifically estimated values which cannot be directly measured such as the *in vivo* neutralization potency (IC50) of each bnAb, percentage of infected cell death due to bnAb Fc-mediated killing, and pre-treatment levels of bnAb resistant strains. We then performed counterfactual simulations with the validated model to quantify the relative impact of each mechanism on HIV viral kinetics over time during a single infusion trial.

While VRC07-523-LS emerged as the superior therapeutic bnAb across our holistic metrics, each bnAb had instances of drug resistance which manifested as occasional immediate non-response or, more commonly, as rapid selection of an apparently fit but sub-dominant resistant variant. Had there not been a pre-existing resistant mutant, our model suggests that many participants would have suppressed viremia for over a month. This result is in keeping with historical studies showing rapid emergence of drug resistance with use of ART monotherapy in HIV and SARS-CoV-2 targeting bnAbs^69,70^. Given the highly mutagenic nature of HIV *env*, the presence of sub-dominant resistant strains was unsurprising, but our model was uniquely able to estimate the kinetics of these strains as they underwent selection. Moreover, our results of particularly high rates of resistance for 10-1074 are consistent with sequencing studies which binds the V3 glycan supersite instead of the CD4 binding site like the other bnAbs^71^. Existing models suggest that optimal dosing of two or three bnAbs with separate molecular targets may limit resistance^22,23,25^, which generally agrees with human clinical trial data in which combination bnAbs slow viral rebound after ATI^24,39,40,72^.

Our model suggests the existence of fully and partially resistant variants. If fully resistant viruses predominated at therapy onset, then no viral load reduction was observed. If they existed as minority variants at treatment onset, then we observed rapid selection. In many cases, the replacing strains during bnAb therapy had *in vivo* IC50 levels which were only 2-10 times greater than the initial dominant strain. The viral load of these partially resistant strains decreased during the first week of treatment when bnAb levels were high but re-expanded sooner than the originally dominant strain. Whether partial resistance becomes an important clinical issue for bnAbs dosed in combination remains to be seen^73,74^.

Even absent resistance, bnAb redosing will likely need to occur frequently to prevent re-expansion with wild type virus. Our counterfactual simulations suggest that the presence of a resistant strain decreases the time to viral re-expansion in many cases, but only by a median of two weeks. The need for redosing is in part due to the 1-6 week half-lives of these bnAbs. VRC-07-523-LS and VRC-01-LS were successfully engineered to have meaningfully longer half-lives but still were predicted to drop below in vivo IC50 and become subtherapeutic 2-3 months of dosing despite having a median half-life exceeding 30 days.

Our model of all bnAb PK simultaneously also concluded that for 3BNC117 and 10-1074, bnAb half-life was shorter in PWH versus HIV negative individuals. The half-life was even shorter in viremic PWH who were not on ART. Our PK model was non-mechanistic and intended only to precisely reproduce bnAb levels. Our preliminary assessments relating viral load to bnAb half-life did not find strong associations, therefore, we did not explicitly model target-mediated drug disposition here. Further experimental work and modeling will be required to address the interplay between systemic inflammation, viremia, and bnAb clearance and could assist in the design of regimens which further sustain therapeutic bnAb levels.

Our model quantifies the extent to which *in vitro* assays and plasma concentrations underestimate the bnAb concentration required for neutralization and Fc killing. The median scaling factor (prf) between *in vitro* and *in vivo* neutralization varied from ∼40-200 for CD4bs bnAbs and was ∼500 for 10-1074 (a V3gs bnAb), though values also varied within participants receiving the same bnAb. This variation suggests host or viral factors further ad*j*ust *in vivo* neutralization. We did observe bnAb titer (peak bnAb concentration relative to baseline *in vitro* IC50) was moderately predictive of viral load reduction. This relationship indicates a signal in the raw data but also supports modeling as a valuable step in integrating several longitudinal data types to make a more comprehensive conclusion about bnAb dosing.

To be conservative, we suggest using the upper end of our estimates (roughly 200-fold for CD4bs bnAbs, **Table 1**) to pro*j*ect *in vivo* titers when *in vitro* IC50 can be measured. This result is remarkably similar to values estimated to titers required in the context of HIV prevention^48,75^. In absolute terms, assuming a non-fully resistant strain^73,74^, a bnAb concentration exceeding 100 µg/mL was typically suppressive in our model simulations.

Importantly, the prf value incorporates a wide range of mechanisms that differ between *in vitro* assays and *in vivo* neutralization, including but not limited to cell receptor levels, localization of bnAb, and competition with autologous antibodies. In particular, cell-to-cell spread is important for *in vivo* infection and not captured with current TZM-bl assays^76–78^. Additionally, prf also includes discrepancies between plasma concentrations and concentrations at key sites of infection. With respect to this point, theoretically, the LS mutation should only increase half-life. Yet, we observed smaller prf for VRC01-LS vs VRC01, indicating a closer correspondence between *in vitro* and *in vivo* IC50 for the LS variant (**Table 1**). That both LS variants had lower prf could be indicative of slightly less discrepancy between plasma concentrations and other sites.

Each bnAb was inferred to have some enhanced clearance, though these effects were most pronounced for VRC01, 10-1074, and VRC07-523-LS. Our model attributes this enhancement to some Fc-mediated killing activity against infected cells (ADCC, ADCP, and/or CDC). In many participants, cell killing via bnAb activity significantly enhanced viral load reduction. Counterfactual simulations with only neutralization often had a less pronounced decrease in viral load. Yet counterfactual simulations with only Fc-mediated killing sometimes barely lowered viral load, particularly following VRC01LS and 3BNC117 infusion. Still, Fc killing provided considerable synergy with neutralization in these cases. For many participants receiving VRC01 and 10-1074, Fc killing alone reduced viral load to a greater extent than neutralization alone.

The impact of Fc-mediated killing for VRC07-523-LS was sustained for many weeks due its prolonged half-life. However, the true duration of this effect is less certain as only three participants had prolonged suppression on VRC07-523-LS without selection of resistant variants. At the time of their last viral load sample, these participants were potentially starting to rebound but were offered ART as part of the trial. Moreover, our estimates for concentration requirements to activate Fc killing are less certain as values at or a below a certain EC50 level allowed equivalent fits to the data – which was corroborated by additional fitting to NHP data. Finally, bnAb infusions have been associated with potential vaccinal effects after full elimination of the bnAb, and we did not model these effects, which cannot be ruled out for long-term suppressors in the present cohorts^35,38,79,80^.

Given the massive clinical benefit of ART, it is somewhat doubtful that Fc-mediated killing would meaningfully enhance prevention of AIDS (and death). However, removal of infected cells may be vital for HIV cure^81,82^. An open question is whether the cell killing activity of bnAbs would be sustained over longer durations of treatment as infected cells convert from more to less activated with considerably less viral antigen presentation. Another hypothesis that could be tested is whether sustained CDC, ADCC or ADCP during prolonged bnAb treatment might decrease reservoir size relative to prolonged ART or rapidly eliminate reactivating cells. If so, then bnAb treatment may be a key part of multi-modality approaches to achieve functional cure.

There was an association between faster viral load drop and faster re-expansion. This result is intuitive from a mathematical modeling point of view, and points to set point equilibria that are maintained by higher absolute values of production and clearance such that removing the production leads to rapid clearance but correspondingly rapid rebound once therapy disappears. This balance also highlights the primacy of full viral suppression throughout the dosing interval when designing future bnAb regimens. This can be accomplished by redosing before bnAb concentrations drops below the *in vivo* IC50, and put in the context of our modeling, keeping bnAb titers >200 (200-fold greater than the *in vitro* IC50). Further technological gains in developing long acting bnAbs may increase possible dosing intervals for these agents.

Our model has limitations. We were not able to estimate possible fitness costs associated with selection of resistant strains^48,83,84^ that could impact evolution during re-expanding HIV^55^. However, the return of similar viral load set points indicates population fitness may not be strongly reduced in practice. We also do not consider other possible drawbacks of bnAbs including development of anti-drug antibodies (ADA), the theoretical possibility of antibody dependent enhancement of infection (ADE), and the considerable breadth needed to cover the global diversity of HIV strains. With respect to our estimates of Fc killing, other mechanisms like anatomic redistribution could also enhance plasma viral load clearance. On the other hand, direct experimental measurements of these phenomena remain difficult and the agreement of models with bnAb infusion data that included Fc-deletion in NHP suggests modeling may currently be the simplest way to estimate *in vivo* Fc-mediated killing. We also do not have a rigorous method to relate neutralization potency to Fc killing potency throughout the dosing interval and some studies conflict on the additive utility of Fc killing for prevention^85,86^. In *in vitro* studies, ADCC activity generally correlated with antibody binding to *env* on the surfaces of virus-infected cells and with viral neutralization^32^. However, neutralization was not always predictive of ADCC, as instances of ADCC in the absence of detectable neutralization, and vice versa, have been observed^44^.

Finally, the model output for a single individual (2D1) did not capture viral kinetics as accurately as for other participants. We predict that the model is missing a therapeutic effect during the first 10 days by over-weighing later datapoints which suggest a lower viral load setpoint: this may be because the replacing bnAb resistant strain was less fit than the pre-treatment predominant strain, or because this individual was not at steady state at bnAb initiation. Participant ID20’s model simulations did not predict strain replacement by our definition but this participant’s increase in IC50 between *V*_1_and *V*_2_was minimal.

In summary, our model provides an interpretive framework for trial participants receiving 5 different HIV bnAbs. Although we derived a single model after substantial investigation into necessary mechanisms, our work could serve as a living document as it is easily updated with new trial data. We identified that that *in vitro* assays overestimate *in vivo* bnAb potency intensely but also to highly varying degrees, that selection of resistant strains is common, and that Fc-mediated cell lysis is a critical mechanism which synergizes with neutralization. Future trials should thus focus on combination agents, which are re-dosed to allow more prolonged HIV suppression while not selecting for resistance.

## Acknowledgments

We sincerely thank the study participants and clinical teams. We are grateful to M Nussenzweig and M Caskey, and the Vaccine Research Center (VRC) teams for generously sharing their data. This research was funded by awards from the NIH NIAID including R01AI186721 to DBR, R01AI179457 to JTS, and R01OD011095, R01AI152703, and P01AI169615 to ASP. The content is solely the responsibility of the authors and does not necessarily represent the official views of the NIH.

## Data and code availability

All data and code used to perform analyses will be freely available on GITHUB upon publication.

## Materials and methods

### Data sources

We use several sources of data to simultaneously match antibody concentration and viral population dynamics. We incorporate studies using bnAbs to suppress HIV-1 viral load. The different bnAbs infused in each of the different studies are VRC01, VRC01LS, VRC01-523-LS, 3BNC117 and 10-1074^59–62^.

#### VRC01

Published data from the VRC 601 single-site, phase 1, open-label, dose escalation studies conducted at the NIH Clinical Center by the VRC Clinical Trials Program, NIAID, NIH (ClinicalTrials.gov NCT and 02840474). In this study, participants with HIV received a single infusion of 40 mg/kg of VRC01 (n=8). Serum VRC01 measurements were collected before infusion and at 0, 1, 2, 4, and 24 hours, and days 2, 7, 14, 21, 28, 35, 42, 49, 56, and 84 after infusion; and HIV RNA plasma samples were collected at baseline, 4 hours, as well as days 1, 2, 3, 5, 7, 14, 16, 21, 28, 35 and 42 after VRC01 administration for all participants, but also days 49, 56, and 84 for two participants for a total of 85 data points from all participants.

#### VRC01-LS and VRC01-523-LS

Published data from the VRC 607 single-site, phase 1, open-label, dose escalation studies conducted at the NIH Clinical Center by the VRC Clinical Trials Program, NIAID, NIH (ClinicalTrials.gov 02840474). In this study participants with HIV received a single infusion of 40 mg/kg of VRC01LS (n=8) or VRC01-523-LS (n=9). Serum antibody concentrations and HIV RNA plasma samples were obtained before infusion and at the end of infusion (0h), 30 min, 1 hr-4 hr, 24 hr, 48 hr and Days 7, 14, 21, 28, 35, 42, 49, 56, 84, 112, 140, 168, 252 and 336 (Week 48) post-infusion.

#### 3BNC117

Published data from a phase 1, dose escalation clinical trial, where the bnAbs 3BNC117 was administered to 22 participants without HIV and 33 participants living with HIV. Participants were given a single infusion of 1mg/kg, 3mg/kg, 10mg/kg or 30mg/kg. Serum antibody concentrations and plasma samples were obtained to measure HIV-1 RNA levels at screening, before infusion and at the end of infusion (0h), 30 min, 6h, 9h, 12h and days 1, 2, 4, 7, 14, 21, 28, 42, 56, 84, 112, 140, and 168 (Week 24) post-infusion.

#### 10-1074

Published data from an open-label, dose escalation clinical trial, where the bnAb 10-1074 was administered to 19 participants living with HIV (3 of these while on ART) and 14 participants without HIV-1. Participants were given a single infusion of 3mg/kg, 10mg/kg, or 30mg/kg. Serum antibody concentrations and plasma samples were obtained to measure HIV-1 RNA levels at screening, before infusion, on days 1, 2, 4, 7, 14, 21, 28, 42, 56, 84, 112, 140 and 168 (Week 24) post-infusion.

#### Pharmacokinetic (PK) Data

Longitudinal samples were collected of antibody concentration in plasma (*µ*g/ml) in each participant ranging from 6 to 48 weeks. The minimum observations per individual is 13, with a maximum of 24 observations.

#### Pharmacodynamic (PD) Data

*In vitro* IC50 was measured twice, once pre-infusion 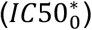 and post bnAb infusion 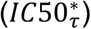 at either day 14, 28, or 35. We classified the participants in four groups according to their two *in vitro* IC50 measurements. The classification is as follows:

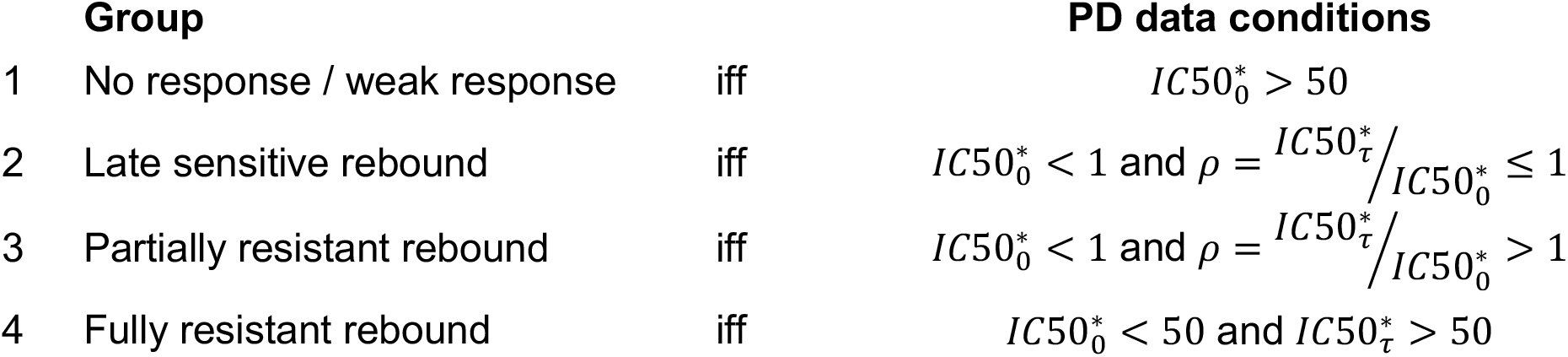

#### Viral Load (VL) Data

Longitudinal samples were collected of HIV RNA (copies/ml) in each participant ranging from 6 to 48 weeks. The minimum observations per individual is 8, with a maximum of 23 observations.

### PD and VL summary metrics

In this study we made use of a set of defined metrics derived from the data for data analysis or from computational simulations for model result analysis. The metrics include initial viral load, final viral load, minimum viral load, time of minimum, start time of re-expansion, maximal viral load reduction, viral load downslope, viral load upslope, time to viral set point, maximal viral load rebound,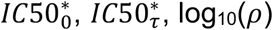. The detailed measures and descriptions are in **Supplementary Table 1**.

### Data clustering

We clustered individuals with HIV viral load into three dynamic groups using k-means clustering as implemented in the Python package scikit-learn 1.8.0^87^. As input features we used *V*_0_, *V*_*min*_, *t*_*min*_, *V*_*f*_, maximal viral load reduction, maximal viral load rebound,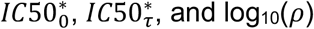.

We tested values of k from 2 to 20 for 3 possible interpolation methods, linear, quadratic, or cubic spline. Comparing these scenarios, linear interpolation had the lowest within cluster sum of squares for some values of *k* ≤ 11. Additionally, comparing the cluster centers using the input features for 3 ≤ *k* ≤ 8, suggests the optimal k values are either 3 or 4. Scatter plots in PCA space showed that *k* = 3 provided the best cluster separation.

### Pharmacokinetic (PK) model

After infusion, the serum antibody concentration displayed a biphasic decay. We use a two compartmental model derived for VRC01 pharmacokinetics in Cardozo-O*j*eda et al. 2021^53^. The model has the form,

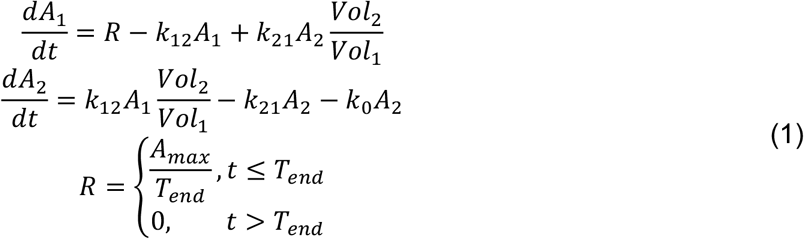

The model assumes the biphasic decay results from antibody distribution from the blood into the tissue followed by elimination from the body. In the model, antibody is infused at rate *R* for the time of infusion (0 < *t* < *T*_*end*_) into *A*_1_ with volume *Vol*_1_ and *A*_*max*_ describing the maximum antibody serum concentration. The infused antibody enters compartment *A*_2_ at a rate *k*_12_, and clears at a rate *k*_0_. Antibody is transported back to *A*_1_at a rate *k*_21_. Now, it is assumed that *T*_*end*_ = 1, and at the end of the infusion, *t* = *T*_*end*_, the antibody concentration in both compartments are

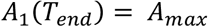

and

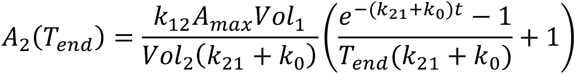

After the end of the infusion, *t*>*T*_*end*_, then *R* = 0. Then the solution of the two-dimensional system is given by the general form

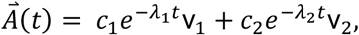

where λ_1,2_ are eigenvalues and v_1,2_ are the corresponding eigenvectors. It is easily verified that the eigenvalues and eigenvector are

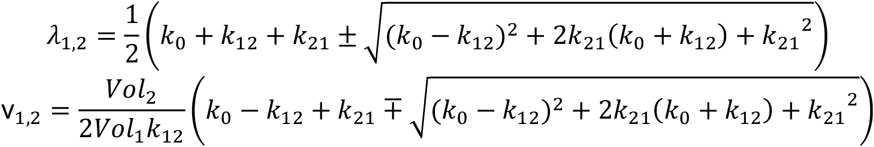

We assume our initial conditions are *A*_1_(0) = *A*_1_(*T*_*end*_) = *A*_*max*_, and *A*_2_(0) = *A*_2_(*T*_*end*_). Thus, the decay of antibody after the end of infusion is given by,

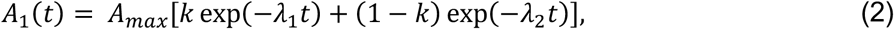

where 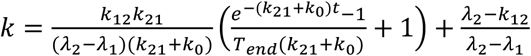

Since the recorded antibody concentration was recorded in plasma, then we fit the data to *A*(*t*) = *A*_1_(*t*).

### Viral dynamic model

We begin with a modified version of our prior PKPD viral dynamics (PKPD-V) model^48^. Let *S*(*t*) susceptible healthy CD4 T-cells, *V*(*t*) plasma detected viremia, and *I*(*t*) HIV infected CD4 T-cells at time *t*. The mathematical model is derived based on the following assumptions:

1. It is assumed the time varying neutralization *υ*(*t*) is mathematically described by a Hill-type logistic function:

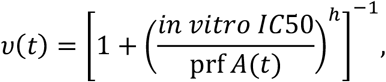

where pr*f* is the potency reduction factor. This term accounts for the overestimation of the 50% titer (*A*(*t*)*⁄in vitro IC*50).
2. It is assumed the susceptible CD4 T-cells have a constant birth rate, *α*, and a natural death rate of *δ*_9_. Susceptible cells can become infected by virus at a rate *β*.
3. The bnAb is not capable to neutralize 100% of the virus. Thus, there is emergence of escape or resistance of *V*, which will be denoted by (1 − *υ*(*t*)). Therefore, the infection term, ((1 − *υ*(*t*))*βVS*), represents the susceptible cells infected by the virus.
4. Infected cells are cleared though natural death at rate *δ*_*I*_.
5. Infected cells, *I*_*j*_, produce virus, *V*_*j*_, at a rate *π*. Virus naturally dies at a rate *γ*. Thus, the basic viral model that include PKPD is as follows:

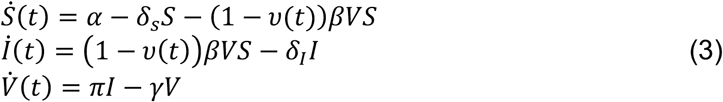

We modify this model with the following additional assumptions:

1. We assume there is a population of infected cells that have longer life span, denoted by *M*(*t*).
2. Within host HIV infection is a diverse quasi-species. Related to our prior models^87^, we assume that, prior to therapeutic intervention, a sample taken from an individual living with HIV will predominantly contain the most abundant strains. Thus, we add two subpopulations for *V*_*j*_(*t*) plasma detected viremia, *I*_*j*_(*t*) HIV infected CD4 T-cells, and *M*_*j*_(*t*) long-lived HIV infected cells at time *t* where *j* = {1,2}. Accordingly, *V*_1_(*t*) denotes the most abundant HIV strains pre-infusion, and *I*_1_(*t*) and *M*_1_(*t*) correspond to the cells infected with the dominant strains.
3. We modify the pharmacodynamic to estimate the in vivo 50% titer. Let *in vitro IC*50_0_ denote the *in vitro* IC50 measurement pre-infusion at day 0 and *in vitro IC*50_*τ*_ the *in vitro* IC50 measured *τ* = {14,28,35} days after the infusion. We assume each subpopulation of HIV strains can be neutralized, which can be described by the following functions.

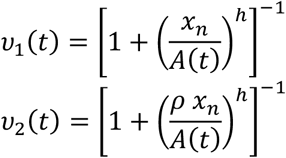

Here *x*_*n*_ = *IC*50_*n*_ denotes the neutralization *in vivo* IC50 and *ρ* = *in vitro IC*50_*τ*_ *⁄in vitro IC*50_0_. Thus, the infectivity term, (1 – *υ*_*j*_(*t*))*jβ*_*s*_*V*_*j*_ *S*), represents the susceptible cells infected by the subpopulations of virus *j*.
4. Let (1 − *µ*) with *µ* ∈ (0,1] be the proportion of infected CD4 T-cells that produce virus. Thus, the term ((1 − *µ*) (1 – *υ*_*j*_(*t*)) *β*_*s*_ *V*_*j*_ *S*) represent the virus producing infected cells that enter compartment *I*_*j*_.
5. We assume that bnAbs also activate Fc-effector function activity with a logistic function (like for neutralization) but with a distinct *EC*50. Thus, the Fc-effector function can be described by the following functions.

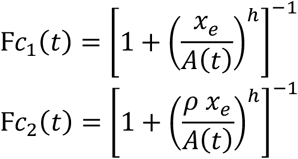

Here *ρ* = *in vitro IC*50_*τ*_ *⁄in vitro IC*50_0_ and 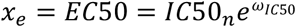 denotes the Fc-effector function *in vivo* IC50 with *EC*50 ≥ *IC*50_*_. Thus, infected cells are cleared via Fc killing with intrinsic rate of ξ. Thus, the effective Fc-effector function activity response can be expressed with the term F*c*_*j*_(*t*)ξ*I*_*j*_.
6. The term (*µ* 1 – *υ*_*j*_(*t*)) *β*_*s*_*V*_*j*_*S*) describes the infected long-lived cells that enter the *M*_*j*_ compartment. Additionally, long-lived infected cells become virus producing cells at a rate Φ.
7. We assume virus has an additional clearance rate. As virus infects the susceptible cells viral particles are removed from the active virus from population *V*_*j*_ at a rate ((1 – *υ*_*n,j*_(*t*)*jβ*_*v*_*V*_*j*_*S*). Lastly, we assume *β* = *β*_*s*_ = *β*_*v*_.

Based on the assumptions above, our final mechanistic model is

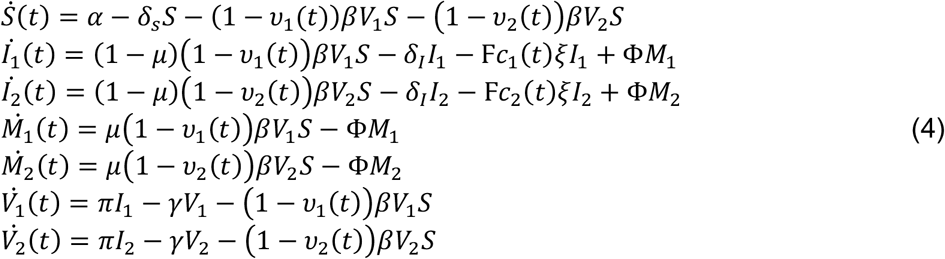

Where total viral load *V*(*t*) = *V*_1_+*V*_2_. See **Supplementary Table 7** for a description and units of each parameter in the mechanistic model.

We assume that at time *t* = 0 the model is at steady state in the absence of treatment. Thus, equation (4) simplifies to

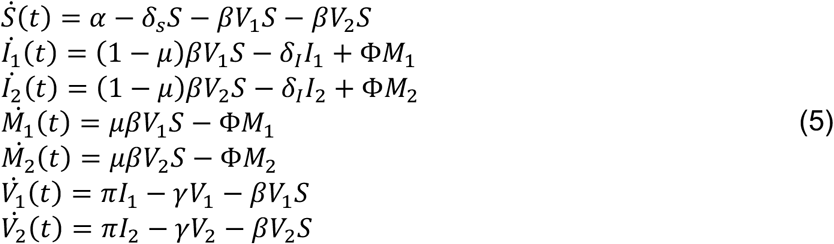

Let *V(t)=V*_*1*_*(t) + V*_*2*_*(t)*. Thus, the initial condition for the system at steady state is

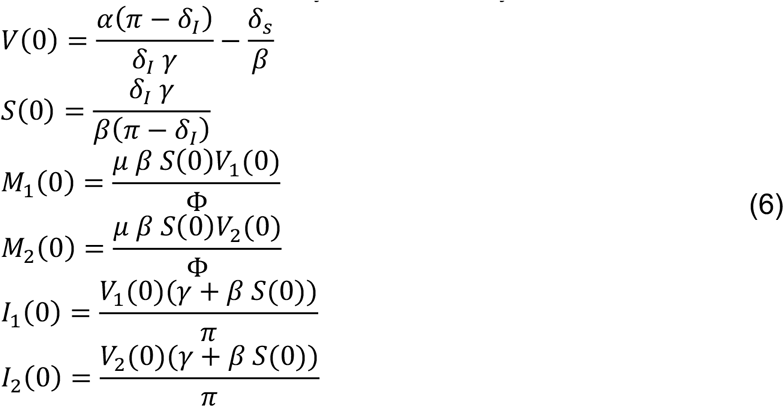

Furthermore, we assume that a proportion (*f*_*s*_) of the initial viral load, *V*(0), are the most abundant strains sampled from each participant. Thus, we have the following initial conditions for the subpopulations of viremia,

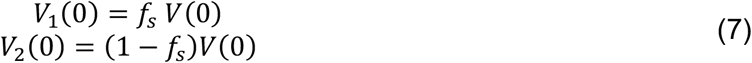

### PK model fitting and selection

We first fit our PK model in equation (2) using a population nonlinear mixed effect modeling (pNLME) approach^89^. With this approach, an antibody concentration measurement from an individual *i* at a time point *T* is modeled as log_10_ (*y*_*iT*_) = *f* _*𝒱* =_(*t*_*iT*_, *θ*_*i*_)+*ϵ*, where *f* _*𝒱* =_ represents the numerical value of *A*(*t*) in equation (2) describing the plasma antibody concentration, *θ*_*i*_ is the parameter vector for individual *i*, and *ϵ*∼*N*(0, *σ*^2^) is the measurement error for the log_10_-transformed bnAb concentration data. In pNLME, each individual’s parameters are the sum of the average population value *θ*_*pop*_ and a random effect encompassing their deviation from the average, *η*_*i*_; or *θ*_*i*_ = *θ*_*pop*_+*η*_*i*_. We fixed *σ* = 0.07 log_10_ bnAb plasma concentration *µ*g/mL when comparing model fits, so that any differences in likelihood of the full model occur due to a change in agreement between model simulations and data rather than estimated magnitude of the measurement error.

For PK model selection, we analyzed the 111 individuals who received a single infusion of one bnAb. Across the cohorts, individuals enrolled in the clinical trials include PWH off ART, PWH on ART, and individuals with no infection. The dose for infusion were 1 mg/kg, 3 mg/kg, 10 mg/kg, 30 mg/kg or 40 mg/kg (further details in original clinical trials^59–62^). The candidate models that we considered are listed in **Supplementary Table 2**. For each candidate model, we used the Stochastic Approximation of the Expectation Maximization (SAEM) algorithm^88^ embedded in the Monolix software (Monolix 2024R1, Lixoft SAS, a Simulations Plus company) to obtain the maximum likelihood estimation (MLE) of the vector of fixed effects, *θ*_*pop*_, and the MLE of the vector of standard deviation of the random effects, Ψ_*θ*_, for the model parameters *A*_*max*_, *k*, λ_1_, and λ_2_. We assumed a log-normal distribution for parameter values *A*_*max*_ and λ_2_ ; and a logit-normal distribution for *k* and λ_1_. The parameter *k* was assumed to fall between 0 and 1, and parameter λ_1_ was assumed to fall between values 0.15 and 3 to avoid fits that result in switching the interpretation of λ_1_ and λ_2_.

Using the parameter set with the highest likelihood, we computed the Akaike Information Criterion, *AIC* = −2 log (*max L*)+2*m* for each model, where *L* is the likelihood that the data were generated by this model with these parameter values and *m* is the number of model parameters. Smaller AIC scores indicate that a model is statistically more likely to explain the data. The model with the smallest AIC score assumes there is a correlation between *k*, λ_1_, and λ_2_, parameter *A*_*max*_ have dose as covariate, *k* has antibody type as covariate, λ_2_ has HIV/ART status (PWH off ART, PWH on ART, and HIV negative) and antibody type as covariates, where VRC01-LS and VRC07-523-LS are grouped. VRC01-LS and VRC07-523-LS are grouped since by design, they will have a slower clearance/longer half-life (**Supplementary Table 3**; see **Supplementary Data 2-Table 1** for detailed information of pNLME parameters). All likelihood and information criterion scores are recorded in **Supplementary Table 2**.

### Viral dynamics model selection

Once we selected the best PK model, we used each individual’s estimated PK parameters as fixed parameters to fit the viral dynamic model in equation (4) also using the pNLME framework. We also fit eight other versions of viral dynamics models described in subsection *Viral dynamics model choices and selection score* below and **Supplementary Table 4**. The viral load measurement from an individual *i* at a time point *T* is modeled as log_!%_(*y*_*iT*_) = *f*_*𝒱*_(*t*_*iT*_, *θ*_*i*_)+*ϵ*, where *f*_*𝒱*_ represents the solution of the ODE model for the state variable describing the total amount of virus (*V* = *V*_1_+*V*_2_). For each of the nine ODE models we first let the SAEM algorithm estimated *σ* and was later fixed when comparing model fits, so that any differences in likelihood of the full model occur due to a change in agreement between model simulations and data rather than a drastic increase in the estimated magnitude of the measurement error.

For model selection, we analyzed data from 43 PWH who received a single infusion of one bnAb while off ART, and had their PK, PD, and VL data completely measured (see **Supplementary Data 1** for full details on which participants were included in the fitting). The infused dosage was 1, 3, 10, 30, or 40 mg/kg (further details in original clinical studies^59–62^). The candidate ODE and statistical models that we considered are listed in **Supplementary Data 2-Table 2**. For each candidate model, we used the Stochastic Approximation of the Expectation Maximization (SAEM) algorithm^89^ embedded in the Monolix software (Monolix 2024R1, Lixoft SAS, a Simulations Plus company) to obtain the maximum likelihood estimation (MLE) of the vector of fixed effects, *θ*_*pop*_, and the MLE of the vector of SDs of the random effects, Ψ_*θ*_, for the model parameters *log*_10_(*f*_*s*_), *IC*50_*n*_, *ω*_*IC*50_, *log*_10_(*β*), *µ*, ξ, *π, δ*_*I*_ and *α*. We assumed a log-normal distribution for parameter values ξ and *α* ; logit-normal distribution for *log*_10_(*f*_*s*_), *IC*50_*n*_, *ω*_*IC*50_, *µ, π*, and *δ*_*I*_; and a normal distribution for *log*_10_ (*β*). The parameter with logit-normal distributions were assumed to have the following interval *log*_10_ (*f*_*S*_) ∈ (−0.22, 0), *IC*50_*n*_ ∈ (0, 1000), *ω*_*IC50*_ ∈ (0, 1000), *µ* ∈ (0,0.49), *π* ∈ (10, 10000), and *δ*_*I*_ ∈ (0.26, 0.85). We assumed fixed values for parameters *δ*_*s*_ = 0.01, *γ* = 23, *h* = 1.1, and *ϕ* = 0.05 for all individuals, that is no random effect was estimated in these parameters. Additionally, we fixed the parameter *δ*_*I*_ = 0.5 population value while allowing random effects to be estimated.

To select the best model for the data, we developed a strategy with several steps. We first fit a mechanistic model with pNLME. Next, we iterated by removing random effects on parameters that were clearly not identifiable (RSE > 50%) and adding covariates to certain parameters to determine the best statistical formulation for that mechanistic model. Then, we assessed the biological plausibility of the model by comparison of parameter values to previous estimates in HIV literature. Lastly, we compared the model predicted ratio between *V*_1_ and *V*_2_ to determine whether “takeover” occurs in agreement with IC50 observations that changed substantially during an individual’s study period. In summary, we rank models by BICc, the takeover number (meaning matching the change or not of IC50 in each of 43 individuals), and made some qualitative choices about plausibility regarding parameter values with respect to additional published data. In **Supplementary Data 2-Table 2**, we present the model scores from 350 tested models.

The best model ultimately had the second lowest BICc, 42 out of 43 correct strain takeovers, and biologically plausible parameter values (**Supplementary Table 5 & Supplementary Data 2-Table 3**). The chosen model assumes there is a correlation between *β* and *α*, parameter *f*_*s*_ and *π* has the cluster type as covariate, parameter *IC*50_*n*_ has the *in vitro* IC50 pre-infusion value as a continuous covariate, the rate of Fc killing (ξ) has antibody type as covariate, and *β* has the cluster type as covariate, where cluster 2 and 3 are grouped (**Supplementary Table 5**; see **Supplementary Data 2-Table 3** for detailed information of estimated population parameters). PD, PK and VD data with their respective fitting results for each individual included in this study can be seen on **Supplementary Data 3**.

### Covariates on PK and viral dynamics parameters

We used seven different covariates to iteratively improve the statistical model fitting. Six of the covariates are categorical and one was taken as a continuous measure. We use two covariates (*bnAb Cat 1, bnAb Cat 2*) representing the bnAb types in this study. The first covariate, *bnAb Cat 1*, considers each of the five bnAb as a distinct group, while the second covariate, *bnAb Cat 2*, groups VRC01-LS and VRC07-523-LS due to their engineered longer half-life. The third categorical covariate is representative of the dose infused in the participants, and thus, this covariate includes four groups for the dosages: 1 mg/kg, 10 mg/kg, 30 mg/kg, and 40 mg/kg. The fourth categorical covariate pertains to the HIV and ART status of the participants, and thus includes PWH off ART, PWH on ART, and HIV negative individuals. The fifth and sixth categorical covariate pertains to the clustered individuals through their viral load dynamics. The fifth covariate has three groups, one for each cluster, while the sixth covariate groups Cluster 2 and 3. The final covariate uses the pre-infusion *in vitro* IC50 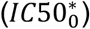 as a continuous measure.

### Viral dynamics model choices and selection score

The models described in this subsection (M1–M8) represent the candidate models considered during mechanistic model selection, which Eq. (4) was chosen from. Note that equation (4), presented earlier, reflects our final selected model. We started with our baseline viral dynamics model (M1), which includes susceptible CD4+ T cells, as well short- and longer-lived infected cells which produce HIV RNA and are necessary to capture two-phase decay. HIV RNA infects cells in a density dependent fashion allowing a steady state pre-bnAb dosing. We allow for two strains of virus in the model for the selection of bnAb resistant viruses. We assume that short- and long-lived infected cells can be infected with either *V*_1_ or *V*_2_. The model includes bnAb neutralization function that decreases rate of infection and bnAb Fc killing that removes infected cells. Both these terms are time dependent and related to bnAb levels and potency. The model (M1) assumed that a fraction *f* of infected cells that become actively infected cells (*I*_1_, *I*_2_), while the remaining fraction (1 − *f*) dies before producing any new virus.

With this initial model, we have previously considered that two types of viruses/infected cells (*V*_1_ and *V*_2_) are not required to match observed viral load data but are clearly needed to explain changes in viral sensitivity over time. This implies that models with only one type that are not matched to time varying IC50 data could also be used to explain the data, but that ultimately conclusions from such a model would likely be inaccurate. This reality supports our choice to include takeover scores in our model selection exercise, as explained in the *Model fitting and selection* subsection.

Model M1

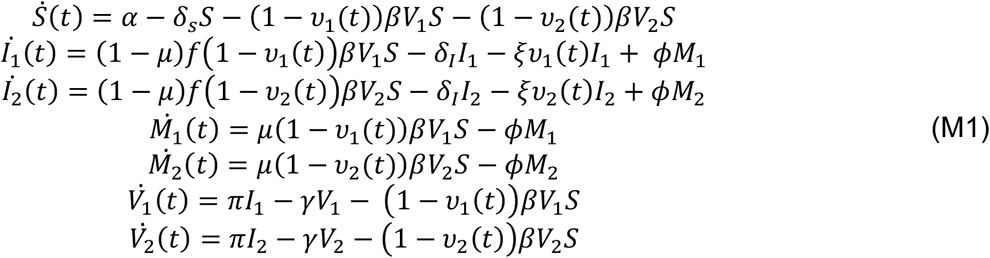

Model (M2) introduces the parameter *f* for the long-lived infected cell compartments (*M*_1_, *M*_2_), representing the possibility that some infection events are abortive, and therefore remove susceptible cells without producing infected cells. This model reduced the BICc score by 265 relative to Model (M1) and improved prediction of the correct strain takeover dynamics (see **Supplementary Table 4**). We next used the mathematical framework to investigate whether the SAEM algorithm could instead distinguish differences in infectivity and viral production between *V*_1_ and *V*_2_.

Model M2

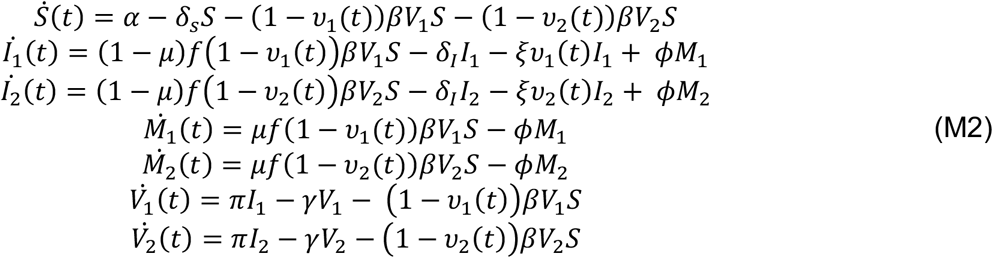

For the next model (M3), we removed the long-lived infected cells compartments (*M*_1_, *M*_2_) to determine if a simpler ODE model was sufficient, but without these compartments, BICc score was 114 points worse than model (M2). Thus, we deemed *M*_1,2_cells necessary.

Model M3

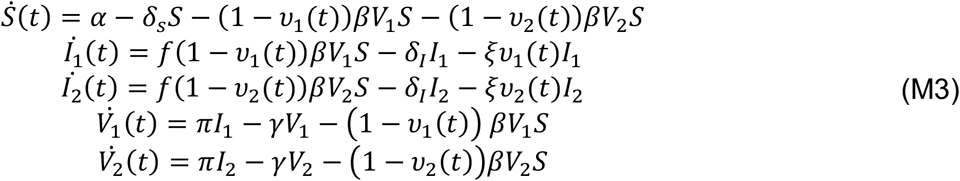

Model (M4) reintroduces the long-lived infected cell compartment. Although, we assume that these cells reactivate to become *I*_1,2_ cells and do not have a death rate in all models. We deemed using an extra death rate for the *M*_1,2_ unnecessary due to the short timescale of the studies, which could lead to the unidentifiability of such parameter. In this model we introduce distinct virion production rates (*π*_1,2_) for each of the viral strains. Thus, this model includes the assumption that different viral strains may be more advantageous and have a better fitness to compete and overtake other strains that do not produce as many virions.

Model M4

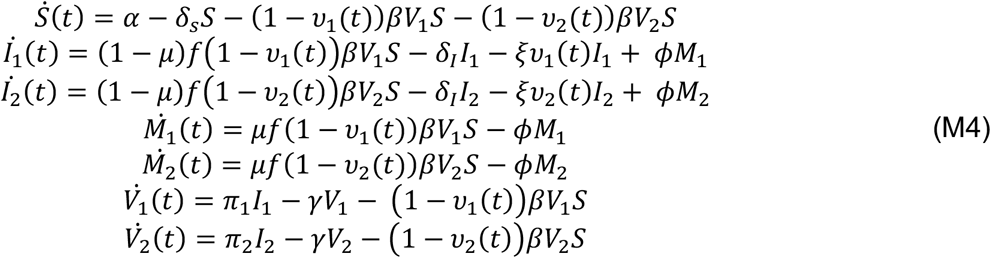

In Model (M5) we wanted to test the assumption that different viral strains have different fitness and can escape the bnAb therapy at different rates. Thus, we introduced distinct infectivity parameters (*β*_1,2_). We additionally assumed that infected cells associated with *V*_2_ (*I*_2_), do not exhibit different neutralization dynamics from *I*_1_, and therefore used a single neutralization function in the model. However, comparison of Models (M4) and (M5), which allow infectivity and viral production to vary between *V*_1_ and *V*_2_, showed no improvement in model fit. This suggests the data is not sufficiently granular to infer obvious fitness costs to emergent viruses.

Model M5

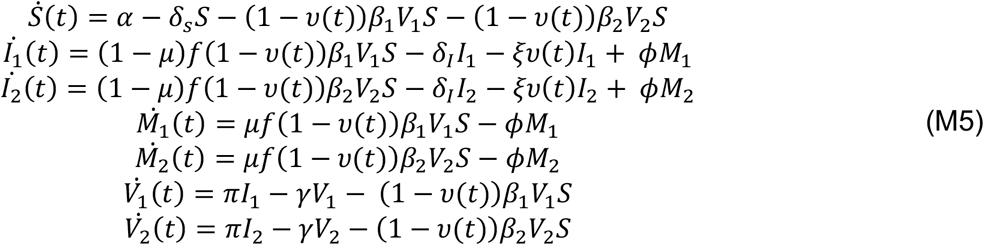

Models (M3), (M4) and (M5) exhibited poorer fits, higher BICc scores and reduced accuracy in predicting strain takeover dynamics compared with Model (M2). Therefore, we reverted to Model (M2) as the baseline framework. Model (M6) then reintroduced distinct neutralization functions for the two types of infected cells while removing the possibility that some fraction of susceptible cells dies before becoming actively infected cells, (or equivalently, *f*=1). This model improved the BICc score by 254 points relative to Model (M2).

Model M6

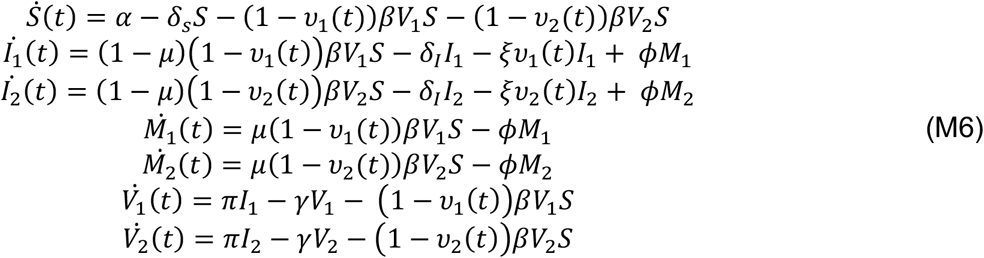

We were interested in testing whether Fc killing could be assumed to be constant through time in Model (M7). Thus, we removed the *υ*_1,2_ functions from the Fc killing of *I*_1,2_ compartments. This model had a BICc score 52 points worse than model (M6), suggesting Fc function is indeed time varying.

Model M7

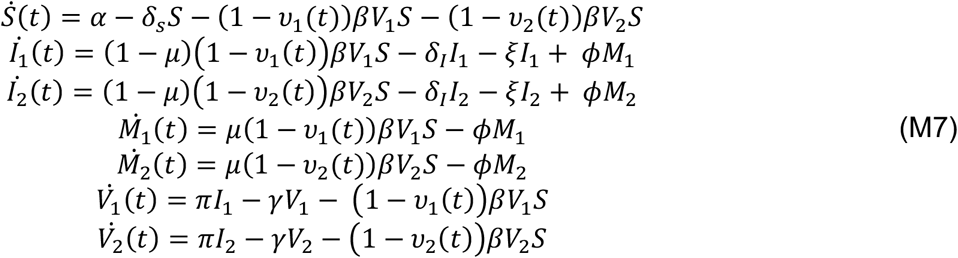

In previous models we assumed that the Fc-function 50% threshold (*EC*50) was equal to the IC50 for neutralization. To relax this assumption, we introduce and estimate a separate parameter (*EC*50) for Fc killing, invoked in the model through a logistic function *Fc*_1,2_. Results from this model (M8) shows that Fc killing likely has a distinct *in vivo EC*50. Additionally, this model shows that *in vivo* sensitivity (for neutralization and Fc killing) appears to differ across bnAbs, but also between individuals receiving the same bnAb. This suggests the viral and/or host context in which therapy occurs changes the efficacy. Finally, the absolute best version of (M8) (**Supplementary Table 4**) inferred that Fc killing was more sensitive than neutralization (i.e., *IC*50>*EC*50), which is unsupported by prior data^44^. When we required that Fc killing sensitivity was less than or equal to neutralization sensitivity (i.e., *IC*50 ≤ *EC*50), this model still provided excellent fit despite not being best by our quantitative model selection (**Supplementary Data 2-Table 2**) but was nevertheless accepted given our prior knowledge about the biology of the system.

Model M8

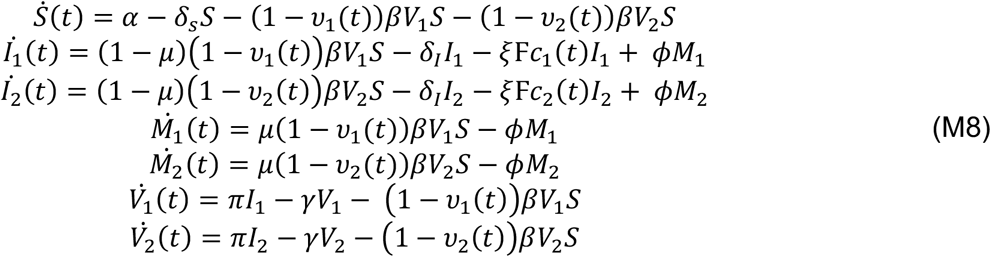

