## Extended Data for "Modeling quantifies in vivo neutralization, Fc-mediated killing, and resistance in human clinical trials of five anti-HIV broadly neutralizing antibodies"

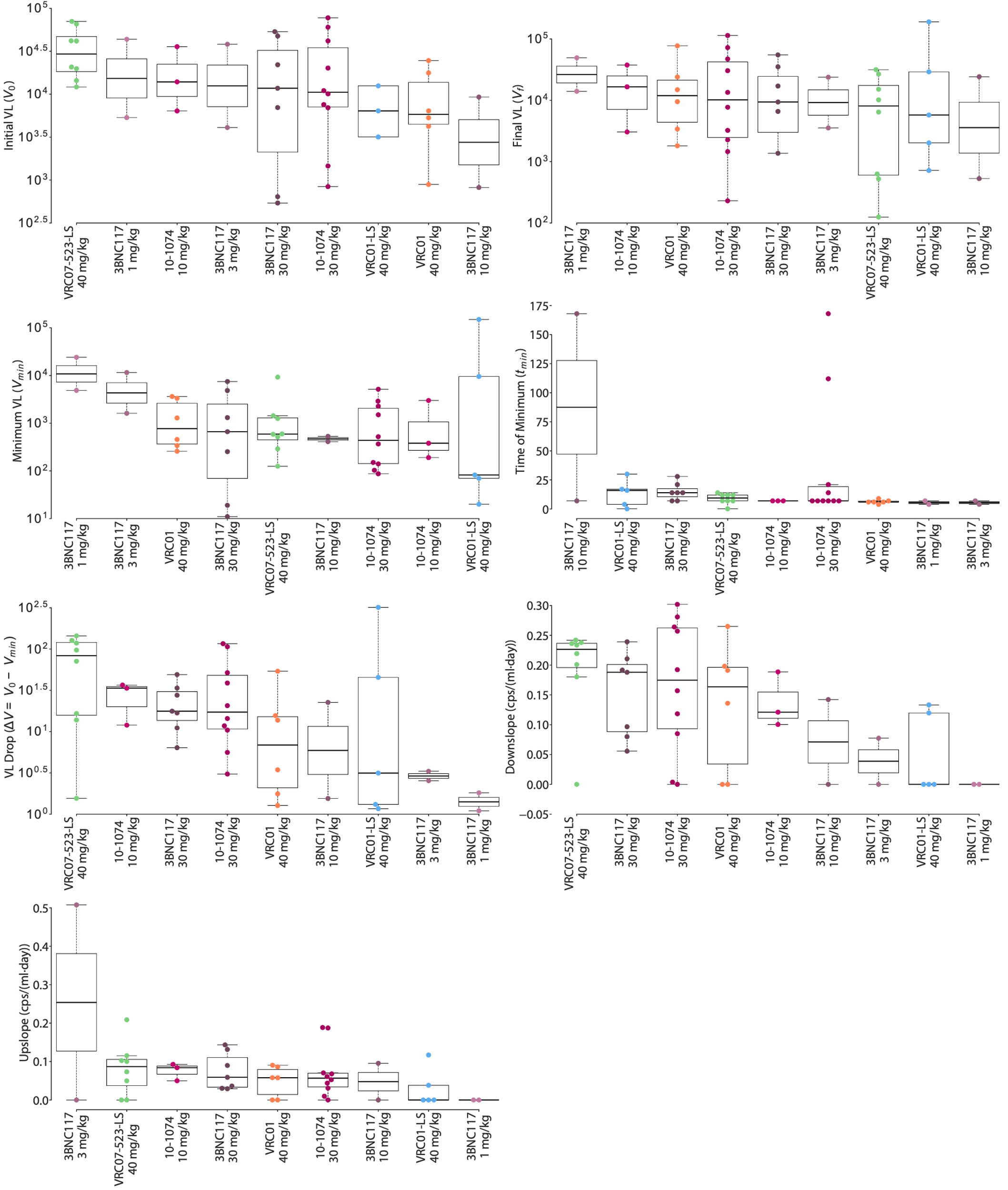

**Extended Data Figure 1.** **Individual viral load metrics according to bnAb and dose.** Metric calculations are described in Methods and Figure 2 in main text.

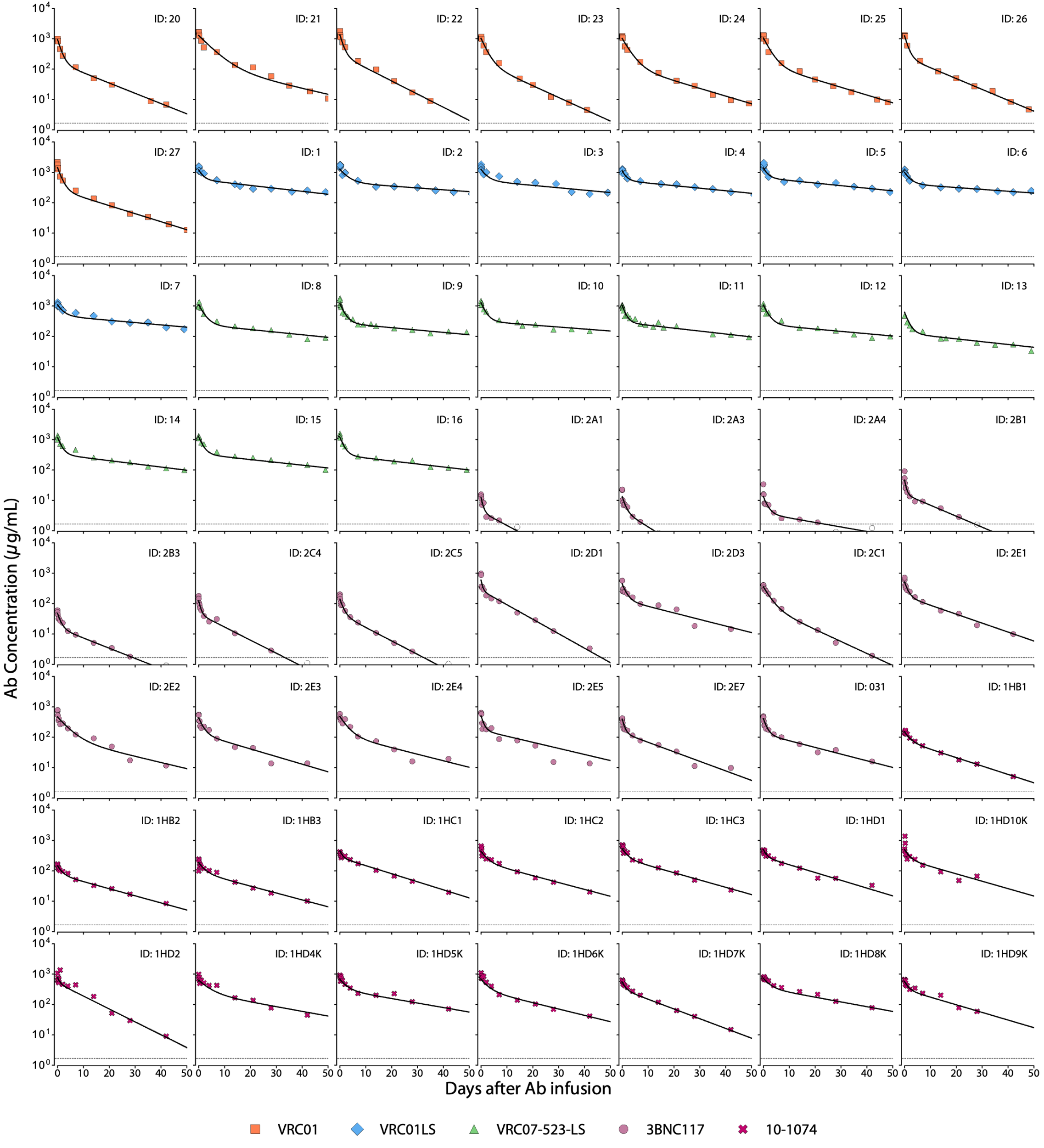

**Extended Data Figure 2.** **PK model fit to bnAb concentration data in PWH off ART.** Different markers and colors indicate bnAb infused (see legend). Black solid line represents the best PK model fit for each person living with HIV who was not taking ART.

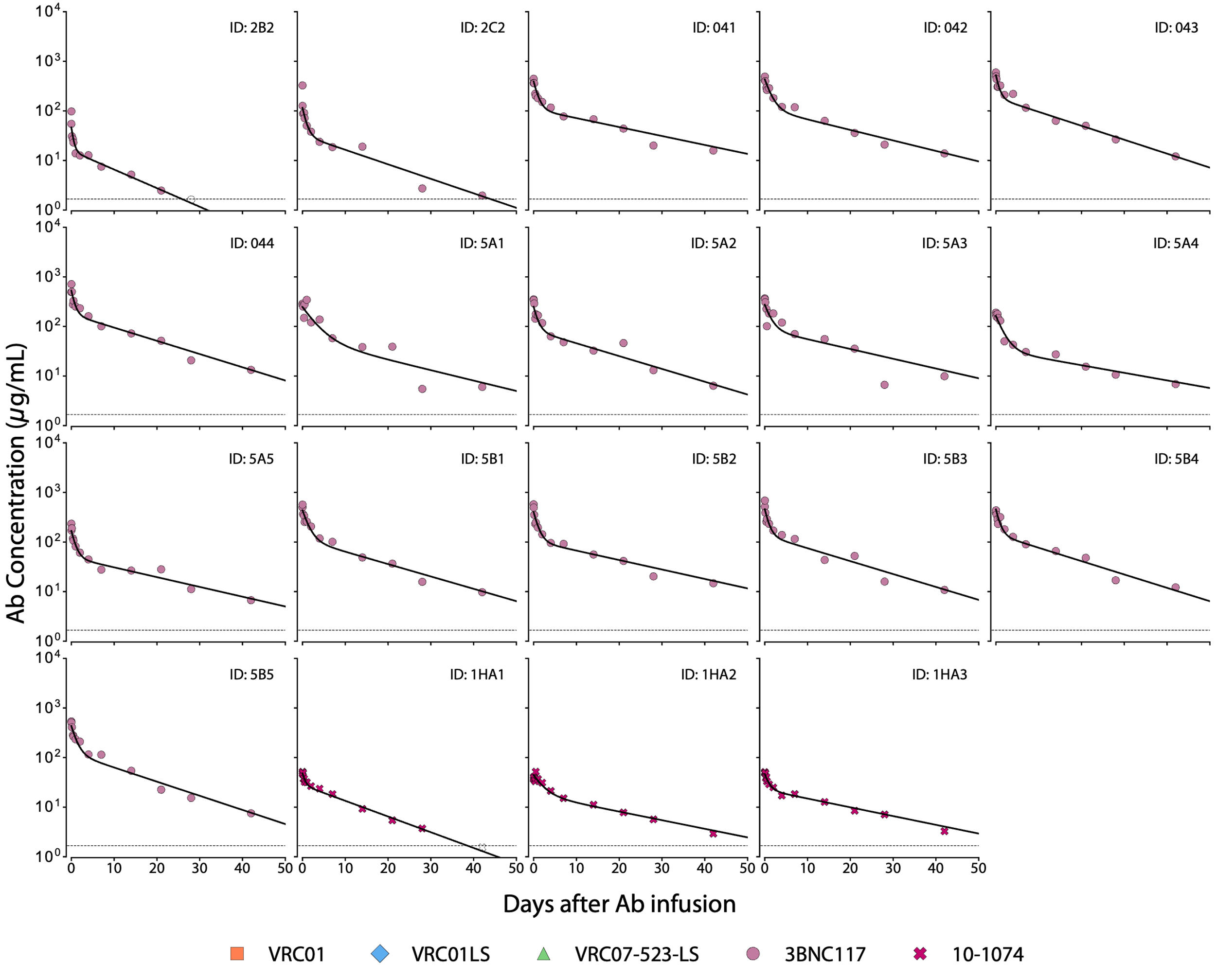

**Extended Data Figure 3.** **PK model fit to bnAb concentration data in PWH on ART.** Different markers and colors indicate bnAb infused (see legend). Black solid line represents the best PK model fit for each person living with HIV who was taking ART.

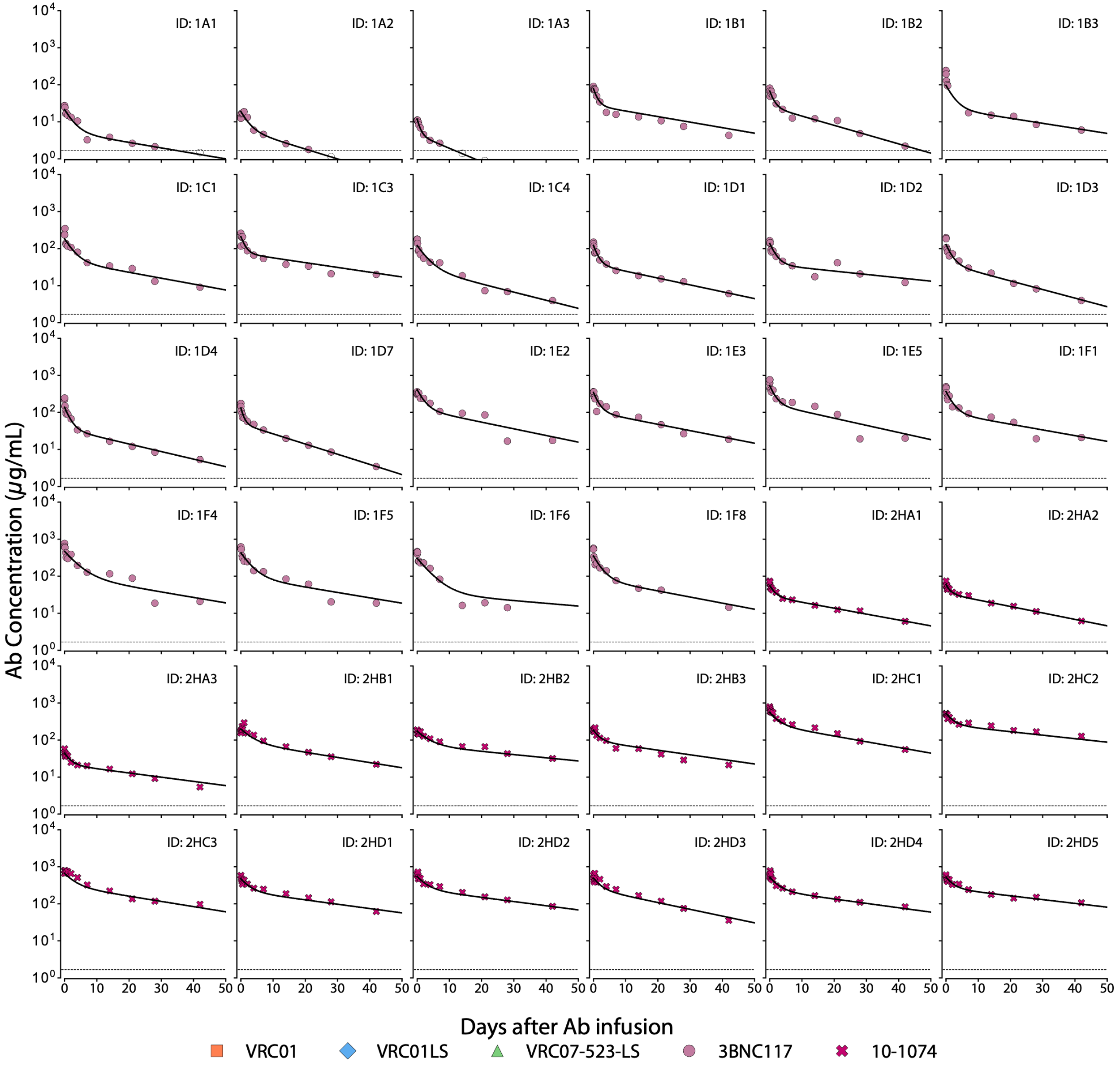

**Extended Data Figure 4.** **PK model fit to bnAb concentration data in HIV negative individuals.** Different markers and colors indicate bnAb infused (see legend). Black solid line represents the best PK model fit for each HIV negative person.

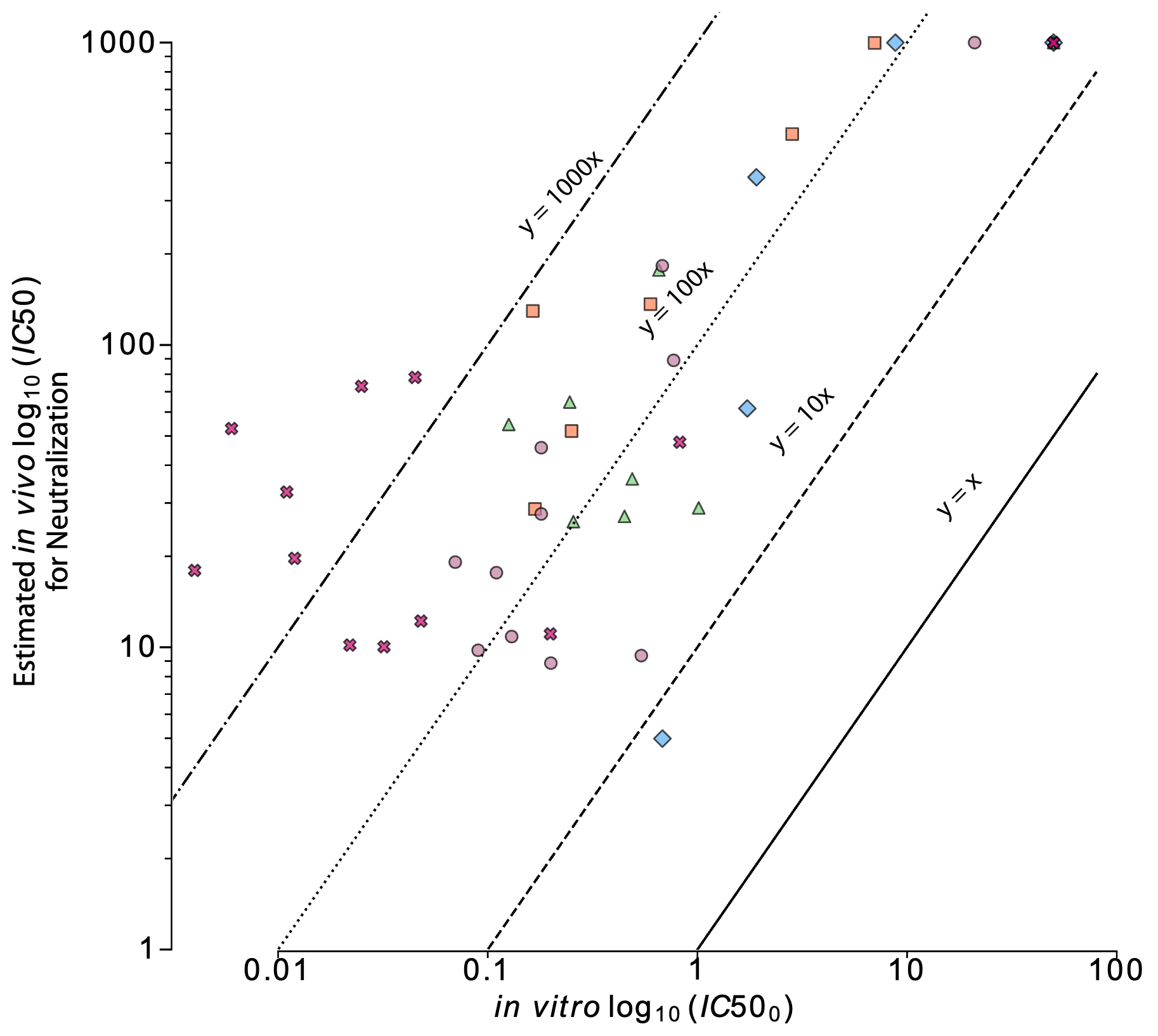

**Extended Data Figure 5. Correlation plot between model estimated neutralization *in vivo* IC50 and *in vitro* IC50 pre-infusion.** The plot shows the estimated fold-change in the bnAb potency *in vivo* to that measured i*n vitro*. Each marker represents an individual. Orange squares are individuals given VRC01, blue diamonds are individuals given VRC01-LS, green triangles are individuals given VRC07-523-LS, purple circles are individuals given 3BNC117, and pink are individuals given 10-1074.

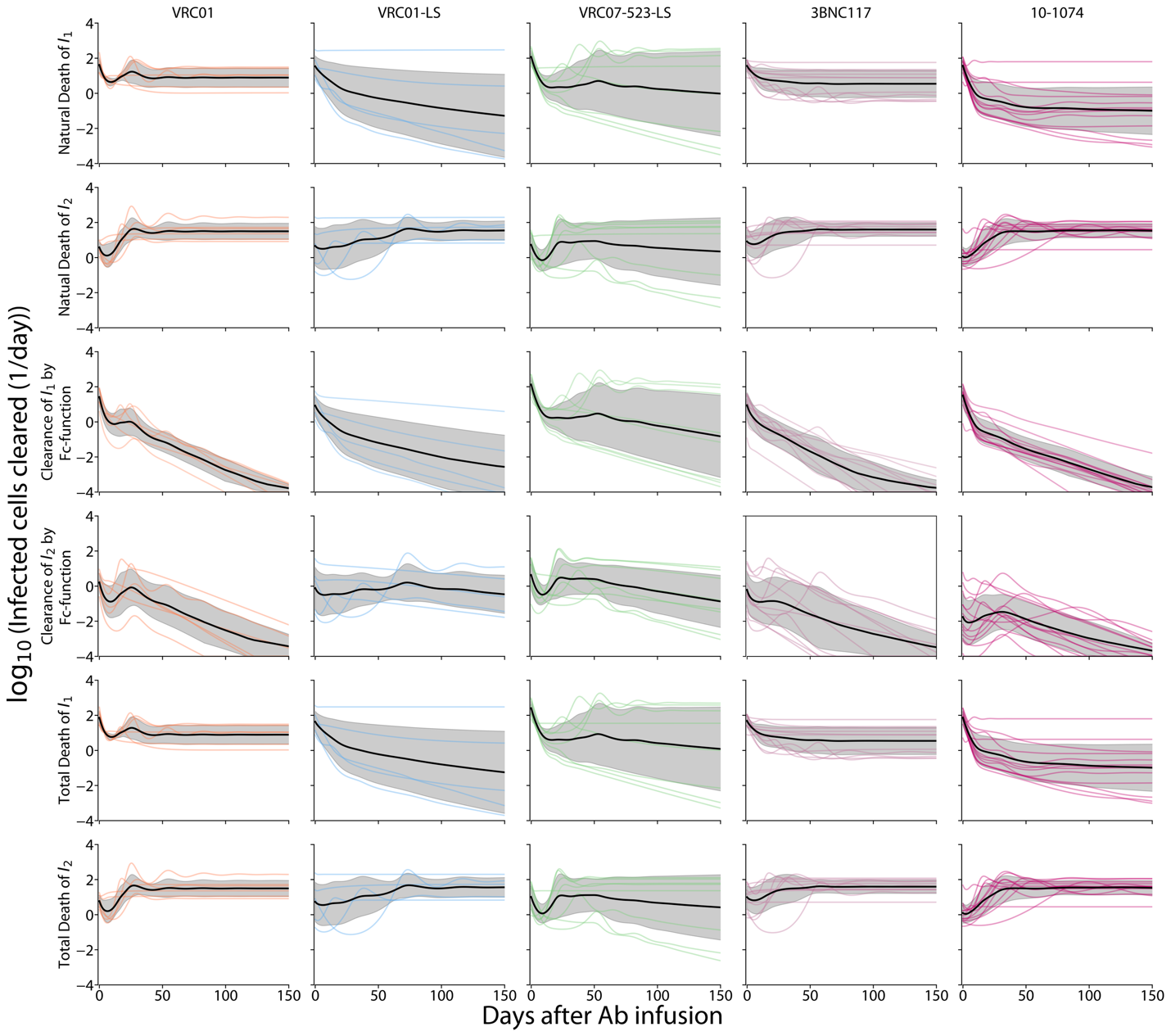

**Extended Data Figure 6.** **Model estimates of the number of infected cells cleared per day by mechanism.** Each column shows individuals infused with one of five bnAbs. Each row indicates the viral-mediated death of $I_{1}$ and $I_{2}$, death by Fc- mediated lysis in $I_{1}$ and $I_{2}$, and the total infected cell death (viral-mediated and Fc-mediated) of each $I$ compartment. Each colored line represents an individual, the black solid line is the mean of the group, and shaded area is the standard deviation.

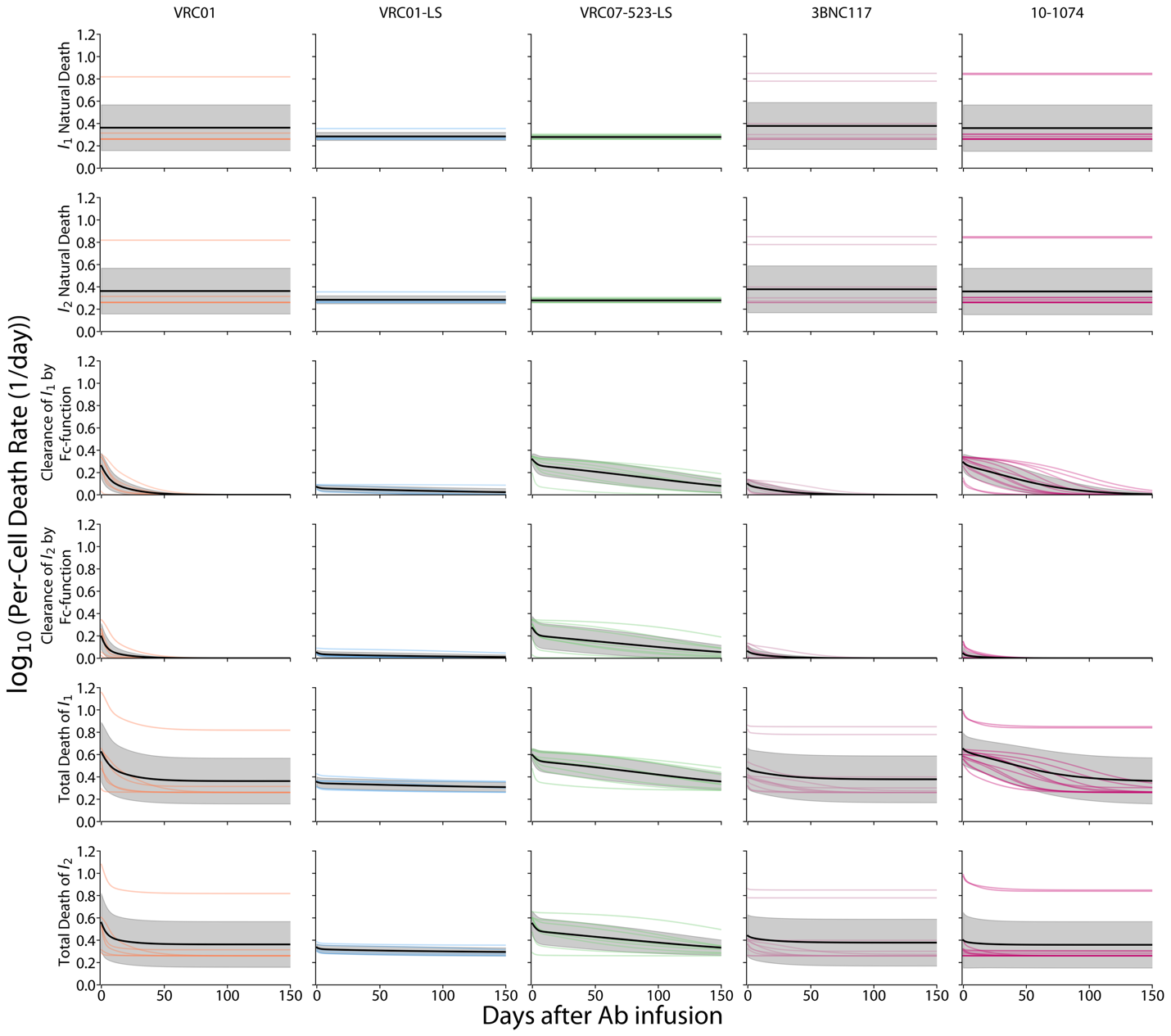

**Extended Data Figure 7.** **Model estimates of per cell death rates by mechanism.** Each column shows individuals infused with one of five bnAbs. Each row indicates the viral-mediated death of $I_{1}$ and $I_{2}$, death by Fc- mediated lysis in $I_{1}$ and $I_{2}$, and the total infected cell death (viral-mediated and Fc-mediated) of each $I$ compartment. Each colored line represents an individual, the black solid line is the mean of the group, and shaded area is the standard deviation.

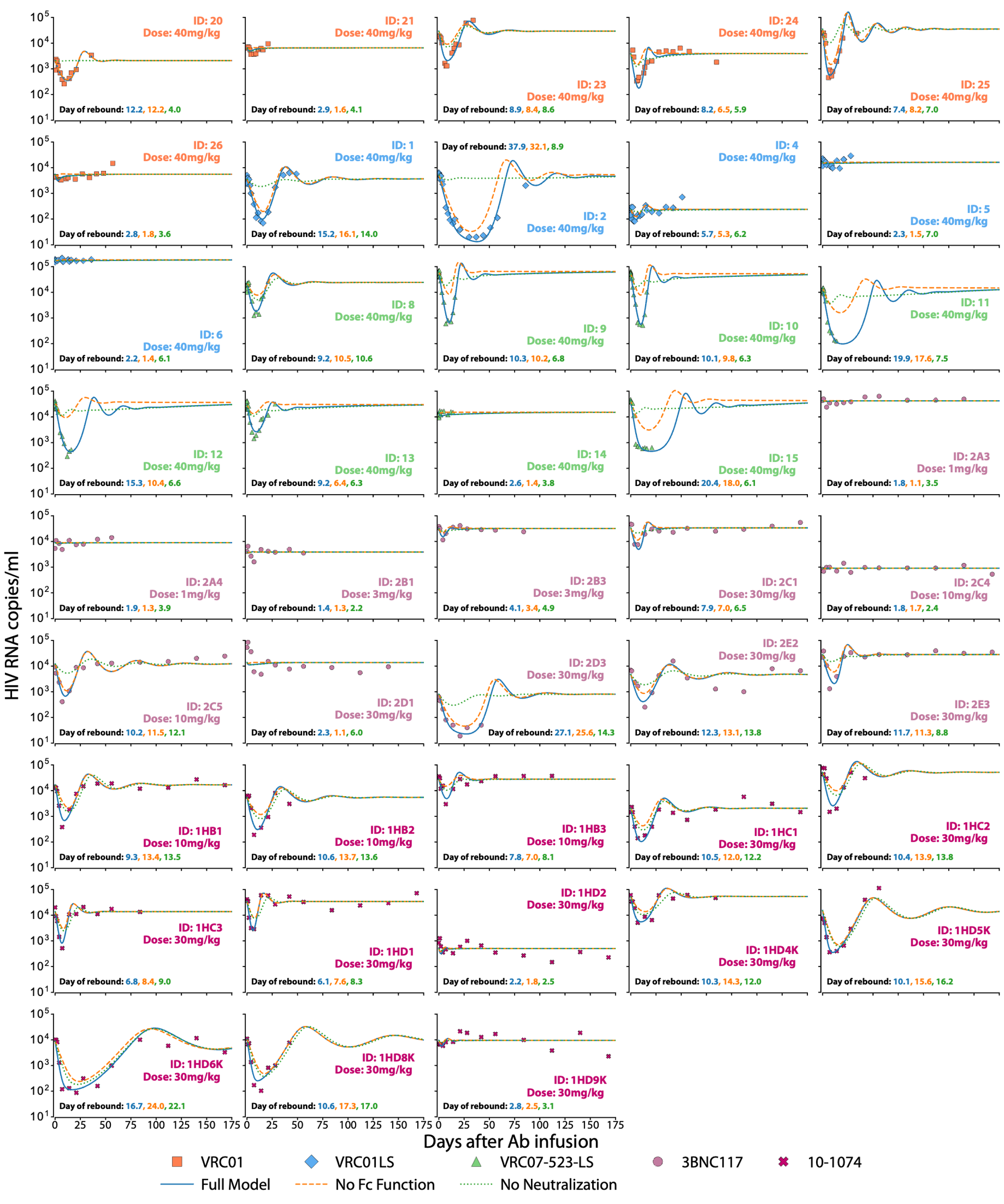

**Extended Data Figure 8.** **Impact of neutralization and Fc-mediated cell lysis on HIV RNA dynamics.** Counterfactual simulations when the effect of Fc-effector functions or neutralization is removed from the VD model. Blue solid line represents the full model, orange dashed line simulates the model without Fc-mediated cell lysis and green dotted line simulates a model without neutralization. Longitudinal HIV RNA (cps/ml) of each enrolled individual is shown by the markers. Different marker and color represent the type of bnAb infused (see legend).

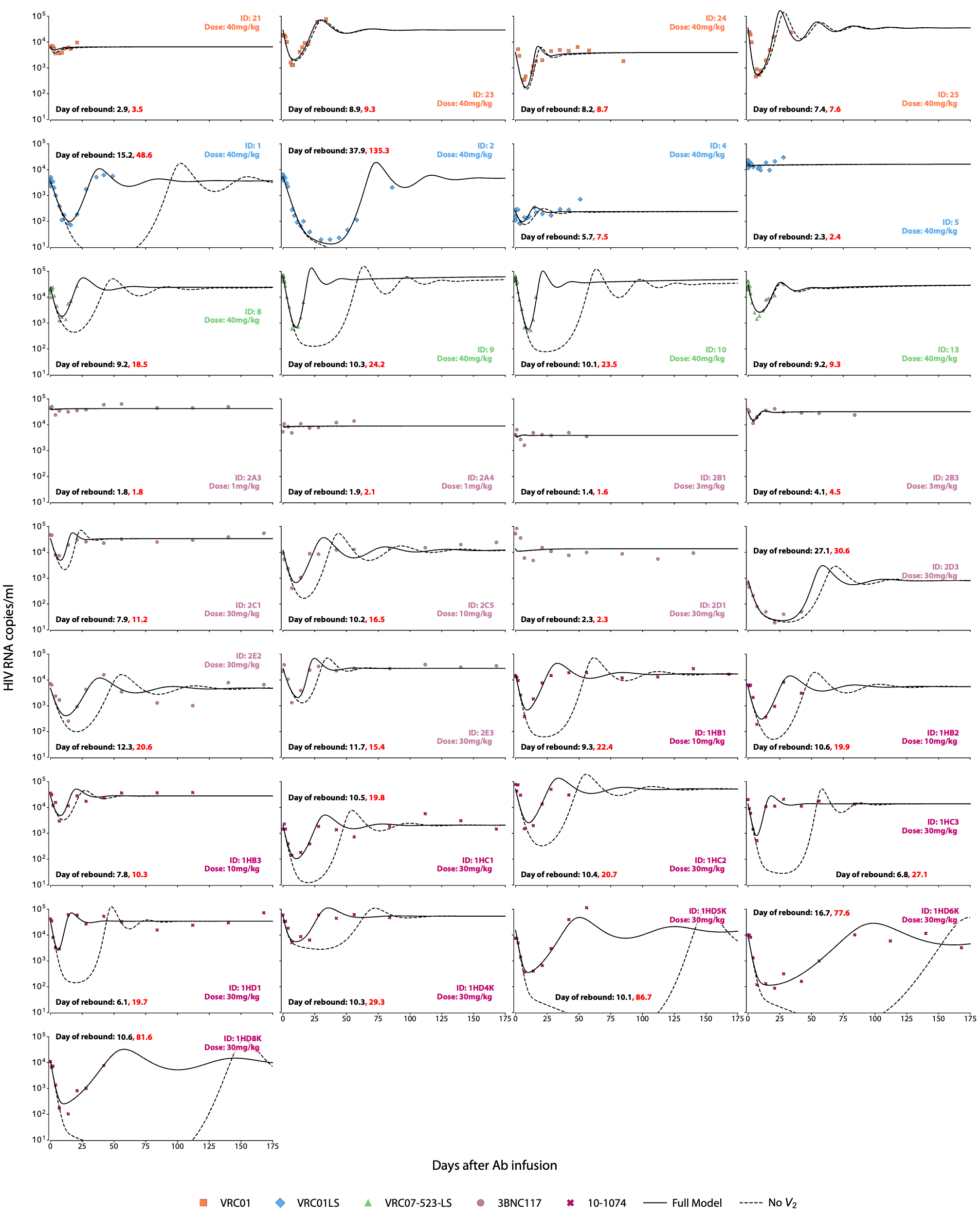

**Extended Data Figure 9. Impact of resistance strain selection on HIV RNA dynamics.** Counterfactual simulations when the sub-detectable strain ($V_{2}$) is removed from the model. Black solid line represents our full model. The dashed line is a model without $V_{2}$ strain. The rebounding VL is due to lack of sufficient antibody to control strain $V_{1}$. Longitudinal HIV RNA (cps/ml) of each enrolled individual is shown by the markers. Different marker and color represent the type of bnAb infused (see legend).

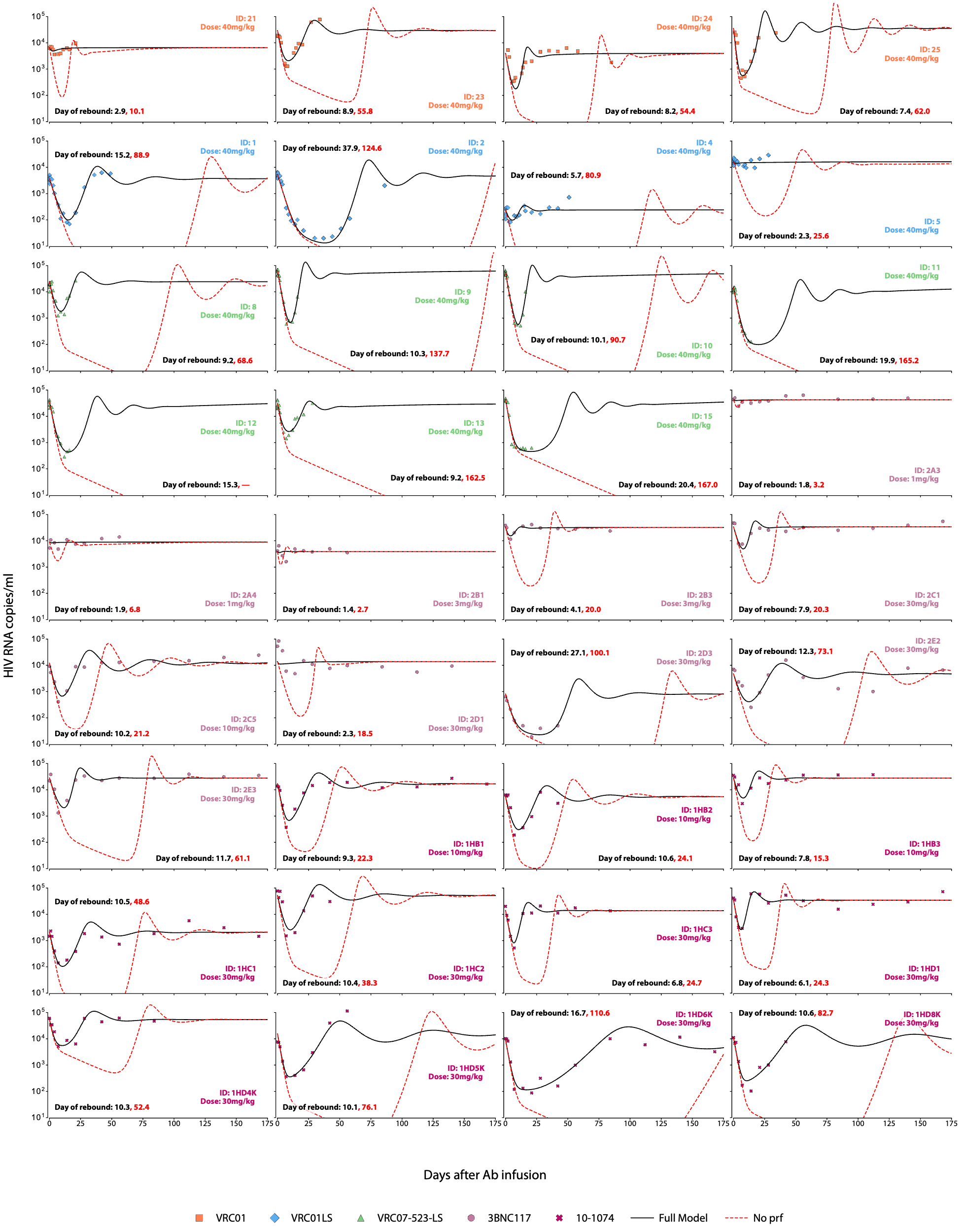

**Extended Data Figure 10. Impact of decreased bnAb potency *in vivo* on HIV RNA dynamics**. Counterfactual simulations when the potency reduction factor (prf) is removed from the VD model. Black solid line represents the full model, dashed red line simulates the model where the *in vivo* bnAb potency is equivalent to *in vitro* estimates. Longitudinal HIV RNA (cps/ml) of each enrolled individual is shown by the markers. Different marker and color represent the type of bnAb infused (see legend).
