## Supplementary Information for "Modeling quantifies in vivo neutralization, Fc-mediated killing, and resistance in human clinical trials of five anti-HIV broadly neutralizing antibodies"

**Supplementary Table 1.** Derived metrics for the analysis of data, and model simulation outcomes.

| **Name** | **Description** |
| --- | --- |
| Initial viral load [log_10_($V_{0})]$ | Viral load (copies/ml) at the first time point |
| Final viral load [log_10_ ($V_{f}$)] | Last viral load (copies/ml) recorded |
| Minimum viral load [log_10_ ($V_{min}$)] | Lowest viral load (copies/ml) recorded (nadir). |
| time of minimum [$t_{min}$] | Time point (day) when nadir occurs. |
| Start time of re-expansion | The day after minimum, one timestep (0.045 days) after $t_{min}$. |
| Maximal viral load reduction / Viral load drop ($\Delta V$) | Initial minus minimum viral load recorded  [log_10_( $V_{0}$) - log_10_ ($V_{min}$)] |
| Viral load downslope | Th rate of viral load decline: estimated as the slope of a linear regression fitted between the maximum viral load preceding the nadir and the nadir time point viral load |
| Time to viral set point / Time of return to equilibrium, $t_{eq}$ | First time after rebound at which the rate of change of log_10_ total viral load begins to stabilize or stays within a tolerance VL of ± log_10_(0.5) log10 VL for 5 days. |
| Viral load upslope | Rate of increase in viral load following the nadir: Estimated as the slope of a linear regression fitted between the nadir time $t_{min}$ and set point time $t_{eq}$ |
| Maximal viral load rebound [log_10_] | Magnitude of rebounding viral load after reaching the nadir [log_10_( $V_{f}$) - log_10_ ($V_{min}$)] |
| ${IC50}_{0}^{*}$ | Geometric mean *In vitro* IC50 pre-infusion at the baseline time point ($0$) |
| ${IC50}_{\tau}^{*}$ | Geometric mean *In vitro* IC50 post-infusion at the second time point ($\tau$) |
| Resistance factor $\rho$ | Ratio of in vitro IC50 from second to baseline time points: ${IC50}_{\tau}/{IC50}_{0}$ |

**Supplementary Table 2.** List of statistical models that were tested for the pharmacokinetic model. The table includes if there was any assumption of correlation between parameter and which parameter was assigned a covariate. Normalized $\Delta$-2LL, $\Delta$BICc, and $\Delta$AIC scores are reported relative to the best model (Run 19 V2). Values in green indicate the best score in each column, and the best-performing model is highlighted in green.

| **Run** | **Parameter** | **Parameter Correlation** | **Covariate** | | | | $\boldsymbol{\Delta}$**-2LL** | $\boldsymbol{\Delta}$**BICc** | $\boldsymbol{\Delta}$**AIC** |
| --- | --- | --- | --- | --- | --- | --- | --- | --- | --- |
|  |  |  | bnAb Cat 1 | bnAb Cat 2 | Dose | HIV/ART Status |  |  |  |
| **Run 19 V2** | $A_{m}$ | $k,\lambda_{1},\lambda_{2}$ |  |  |  |  | 0 | **0** | **0** |
|  | $k$ |  |  |  |  |  |  |  |  |
|  | $\lambda_{1}$ |  |  |  |  |  |  |  |  |
|  | $\lambda_{2}$ |  |  |  |  |  |  |  |  |
| Run 19 | $A_{m}$ | $k,\lambda_{1},\lambda_{2}$ |  |  |  |  | 1.2 | 1.2 | 1.2 |
|  | $k$ |  |  |  |  |  |  |  |  |
|  | $\lambda_{1}$ |  |  |  |  |  |  |  |  |
|  | $\lambda_{2}$ |  |  |  |  |  |  |  |  |
| Run11 | $A_{m}$ | $k,\lambda_{1},\lambda_{2}$ |  |  |  |  | **-1.9** | 7.5 | 2.1 |
|  | $k$ |  |  |  |  |  |  |  |  |
|  | $\lambda_{1}$ |  |  |  |  |  |  |  |  |
|  | $\lambda_{2}$ |  |  |  |  |  |  |  |  |
| Run 11 V2 | $A_{m}$ | $k,\lambda_{1},\lambda_{2}$ |  |  |  |  | -0.9 | 8.6 | 3.1 |
|  | $k$ |  |  |  |  |  |  |  |  |
|  | $\lambda_{1}$ |  |  |  |  |  |  |  |  |
|  | $\lambda_{2}$ |  |  |  |  |  |  |  |  |
| Run 12 | $A_{m}$ | $k,\lambda_{2}$ |  |  |  |  | 19.7 | 19.8 | 19.7 |
|  | $k$ |  |  |  |  |  |  |  |  |
|  | $\lambda_{1}$ |  |  |  |  |  |  |  |  |
|  | $\lambda_{2}$ |  |  |  |  |  |  |  |  |
| Run 10 | $A_{m}$ | $k,\lambda_{1}$ |  |  |  |  | 27.8 | 27.8 | 7.8 |
|  | $k$ |  |  |  |  |  |  |  |  |
|  | $\lambda_{1}$ |  |  |  |  |  |  |  |  |
|  | $\lambda_{2}$ |  |  |  |  |  |  |  |  |
| Run 4 | $A_{m}$ | NA |  |  |  |  | 39.8 | 35.2 | 37.8 |
|  | $k$ |  |  |  |  |  |  |  |  |
|  | $\lambda_{1}$ |  |  |  |  |  |  |  |  |
|  | $\lambda_{2}$ |  |  |  |  |  |  |  |  |
| Run 18 | $A_{m}$ | NA |  |  |  |  | 35.9 | 40.6 | 37.9 |
|  | $k$ |  |  |  |  |  |  |  |  |
|  | $\lambda_{1}$ |  |  |  |  |  |  |  |  |
|  | $\lambda_{2}$ |  |  |  |  |  |  |  |  |
| Run 3 | $A_{m}$ | NA |  |  |  |  | 90.3 | 76.2 | 84.3 |
|  | $k$ |  |  |  |  |  |  |  |  |
|  | $\lambda_{1}$ |  |  |  |  |  |  |  |  |
|  | $\lambda_{2}$ |  |  |  |  |  |  |  |  |
| Run 2 | $A_{m}$ | NA |  |  |  |  | 104.5 | 81.0 | 94.5 |
|  | $k$ |  |  |  |  |  |  |  |  |
|  | $\lambda_{1}$ |  |  |  |  |  |  |  |  |
|  | $\lambda_{2}$ |  |  |  |  |  |  |  |  |
| Run 13 | $A_{m}$ | NA |  |  |  |  | 105.5 | 82.0 | 95.5 |
|  | $k$ |  |  |  |  |  |  |  |  |
|  | $\lambda_{1}$ |  |  |  |  |  |  |  |  |
|  | $\lambda_{2}$ |  |  |  |  |  |  |  |  |
| Run 14 | $A_{m}$ | NA |  |  |  |  | 92.2 | 87.5 | 90.2 |
|  | $k$ |  |  |  |  |  |  |  |  |
|  | $\lambda_{1}$ |  |  |  |  |  |  |  |  |
|  | $\lambda_{2}$ |  |  |  |  |  |  |  |  |
| Run 15 | $A_{m}$ | NA |  |  |  |  | 95.8 | 91.1 | 93.8 |
|  | $k$ |  |  |  |  |  |  |  |  |
|  | $\lambda_{1}$ |  |  |  |  |  |  |  |  |
|  | $\lambda_{2}$ |  |  |  |  |  |  |  |  |
| Run 17 | $A_{m}$ | NA |  |  |  |  | 144.7 | 158.8 | 150.7 |
|  | $k$ |  |  |  |  |  |  |  |  |
|  | $\lambda_{1}$ |  |  |  |  |  |  |  |  |
|  | $\lambda_{2}$ |  |  |  |  |  |  |  |  |
| Run 16 | $A_{m}$ | NA |  |  |  |  | 215.2 | 229.3 | 221.2 |
|  | $k$ |  |  |  |  |  |  |  |  |
|  | $\lambda_{1}$ |  |  |  |  |  |  |  |  |
|  | $\lambda_{2}$ |  |  |  |  |  |  |  |  |
| Run 1 | $A_{m}$ | NA |  |  |  |  | 441.7 | 399.3 | 423.7 |
|  | $k$ |  |  |  |  |  |  |  |  |
|  | $\lambda_{1}$ |  |  |  |  |  |  |  |  |
|  | $\lambda_{2}$ |  |  |  |  |  |  |  |  |

**Supplementary Table 3.** Summary results for the population parameter values for the selected best PK model fit. See **Supplementary Data 2-Table 1** for complete information of estimated population parameters.

| Parameter | Population Parameter (%RSE) | | |
| --- | --- | --- | --- |
| $\boldsymbol{A}_{\boldsymbol{max}}$ | 1mg/kg group: | | 15.15 (10.2) |
|  | 3mg/kg group: | | 55.13 (7.10) |
|  | 10mg/kg group: | | 161.88 (5.63) |
|  | 30mg/kg group: | | 484.05 (3.97) |
|  | 40mg/kg group: | | 1146.54 (7.20) |
| $\boldsymbol{k}$ | VRC01 group: | | 0.84 (2.7) |
|  | VRC01-LS group: | | 0.54 (6.94) |
|  | VRC07-523-LS group: | | 0.77 (4.50) |
|  | 3BNC117 group: | | 0.72 (1.39) |
|  | 10-1074 group: | | 0.49 (3.18) |
| $\boldsymbol{\lambda}_{\boldsymbol{1}}$ | 0.62 (9.85) | | |
| $\boldsymbol{\lambda}_{\boldsymbol{2}}$ | VRC01 group: | HIV Neg group: | 0.043 (12.4) |
|  |  | HIV+ (ART) group: | 0.058 (12.9) |
|  |  | HIV + group: | 0.071 (10.1) |
|  | VRC01-LS and VRC07-523-LS groups: | HIV Neg group: | 0.011 (10.7) |
|  |  | HIV+ (ART) group: | 0.015 (11.4) |
|  |  | HIV + group: | 0.019 (8.11) |
|  | 3BNC117 group: | HIV Neg group: | 0.039 (9.09) |
|  |  | HIV+ (ART) group: | 0.053 (9.84) |
|  |  | HIV + group: | 0.065 (5.82) |
|  | 10-1074 group: | HIV Neg group: | 0.032 (9.24) |
|  |  | HIV+ (ART) group: | 0.043 (9.98) |
|  |  | HIV + group: | 0.053 (6.04) |

| **Model** | **ODE Sub-models in Group** | **Key Structural Difference** | **# Math Models Runs** | **Fit params** | **Best −2LL** | **Best AIC** | **Best BICc** | **Correct Strain Takeover prediction (#/43)** | **Best Run** |
| --- | --- | --- | --- | --- | --- | --- | --- | --- | --- |
| **M1** | - Model11 | Has both V1*S ordering and the f factor in viral equations | 13 | 18 | 715.94 | 751.94 | 814.92 | 34 | Run6_Model11_Assessment_006 |
| **M2** | - Model1 - ModelV1 - ModelV2 - ModelV2a - ModelV2b | Uses f (fraction infectious) factor in M compartments | 26 | 25 | 496.79 | 516.79 | 549.69 | 43 | Run7_ModelV2a |
| **M3** | - ModelV3 | Reduced 5-state model - no M2/M1 memory cells | 18 | 13 | 580.66 | 614.66 | 664.00 | 37 | Run21_ModelV3_Assessment_007 |
| **M4** | - ModelV2b1 - ModelV4 | Separate virion production rates (pi1, pi2) for two viral strains | 31 | 13 | 462.95 | 496.95 | 546.29 | 30 | Run121_ModelV4_Assessment_008 |
| **M5** | - ModelV5 | Single nu - no two viral strain distinction in neutralization. Separate infection rate (beta1, beta2) | 37 | 35 | 332.33 | 374.33 | 428.07 | 36 | Run28_ModelV5_Assessment_010 |
| **M6** | - ModelV2c | Uses nu1, nu2 neutralization without the f factor | 63 | 15 | 223.38 | 253.38 | **295.61** | 41 | Run17_ModelV2c_Assessment5 |
| **M7** | - ModelV2_constantADCC | Constant ADCC term Xi*pIr without neutralization factor | 46 | 28 | 211.27 | 271.27 | 347.83 | 37 | Run23_ModelV2_ConstantADCC_Assessment_008 |
| **M8** | - ModelV2c_2IC50 - ModelV2c_2IC50V2 - ModelV2c_2IC50V3 | Two IC50s: separate functions for neutralization and Fc function activation | 115 | 31 | **198.87** | **250.87** | 325.65 | 42 | Run243_ModelV2c_2IC50_V3_2IC50_V3_Assessment10 |

**Supplementary Table 4. Summary of 8 distinct ODE models that were tested.** Rows are ordered by earliest run execution date. Two files are in the same model group if and only if they share identical dynamic equations. Files within a group may still differ in auxiliary definitions, initial conditions, or constants. A total of 350 distinct mathematical model where run. # Math Models Runs is the number of mathematical models tested of each model group (out of 350), which accounts for different mechanistic models, statistical models and assessments, # Fit Params counts MLE-estimated parameters only; fixed parameters are excluded. Best −2LL, AIC, and BICc are all taken from the single minimum BICc run in each group. See **Supplementary Data 2-Table 2**, for detailed scores from 349 tested models. Best Run is selected by BICc, strain takeover prediction, parameter identifiability, and biologically significant statistical model.

**Supplementary Table 5.** Summary results for the estimated and fixed population parameter values on the selected best mechanistic model fit. See **Supplementary Data 2-Table 3** for information of estimated population parameters.

| Parameter | Population Parameter (%RSE) | |
| --- | --- | --- |
| $\boldsymbol{log}_{\boldsymbol{10}}\left( \boldsymbol{f}_{\boldsymbol{s}} \right)$ | Cluster 1: | -0.0066 (0.000088) |
|  | Cluster 2: | -0.033 (0.00061) |
|  | Cluster 3: | -0.22 (0.0000000009) |
| ${\boldsymbol{IC}\boldsymbol{50}}_{\boldsymbol{n}}$ | 17.52 (11.32) | |
| $\boldsymbol{\omega}_{\boldsymbol{IC}\boldsymbol{50}}$ | 0.00013 (34.83) | |
| $\boldsymbol{log}_{\boldsymbol{10}}\left( \boldsymbol{\beta} \right)$ | Cluster 1: | -5.29 (0.16) |
|  | Cluster 2 & 3: | -4.07 (0.3) |
| $\boldsymbol{\mu}$ | 0.011 (0.022) | |
| $\boldsymbol{\xi}$ | VRC01 group: | 0.37 (NaN) |
|  | VRC01-LS group: | 0.093 (NaN) |
|  | VRC07-523-LS group: | 0.35 (NaN) |
|  | 3BNC117 group: | 0.14 (NaN) |
|  | 10-1074 group: | 0.35 (NaN) |
| $\boldsymbol{\pi}$ | Cluster 1: | 2767.74 (0.0000000047) |
|  | Cluster 2: | 411.82 (134.12) |
|  | Cluster 3: | 1696.41 (1259.99) |
| $\boldsymbol{\alpha}$ | 60.05 (20.41) | |
| $\boldsymbol{\delta}_{\boldsymbol{S}}$ (fixed) | 0.01 | |
| $\boldsymbol{\gamma}$ (fixed) | 23 | |
| $\boldsymbol{\delta}_{\boldsymbol{I}}$ (fixed with random effects) | 0.5 | |
| $\boldsymbol{\Phi}$ (fixed) | 0.05 | |
| $\boldsymbol{h}$ (fixed) | 1.1 | |

**Supplementary Figure 3. Model fitting to subset of PWH enrolled in the clinical trials.** Mathematical model captures viral load decay and viral strain dynamics of individuals initially removed from this study due to missing data. Viral load model includes total (solid black line) as well as V_1_ (dashed blue, dominant variant at baseline), and V_2_ (dotted red lines, potentially resistant minor variant).

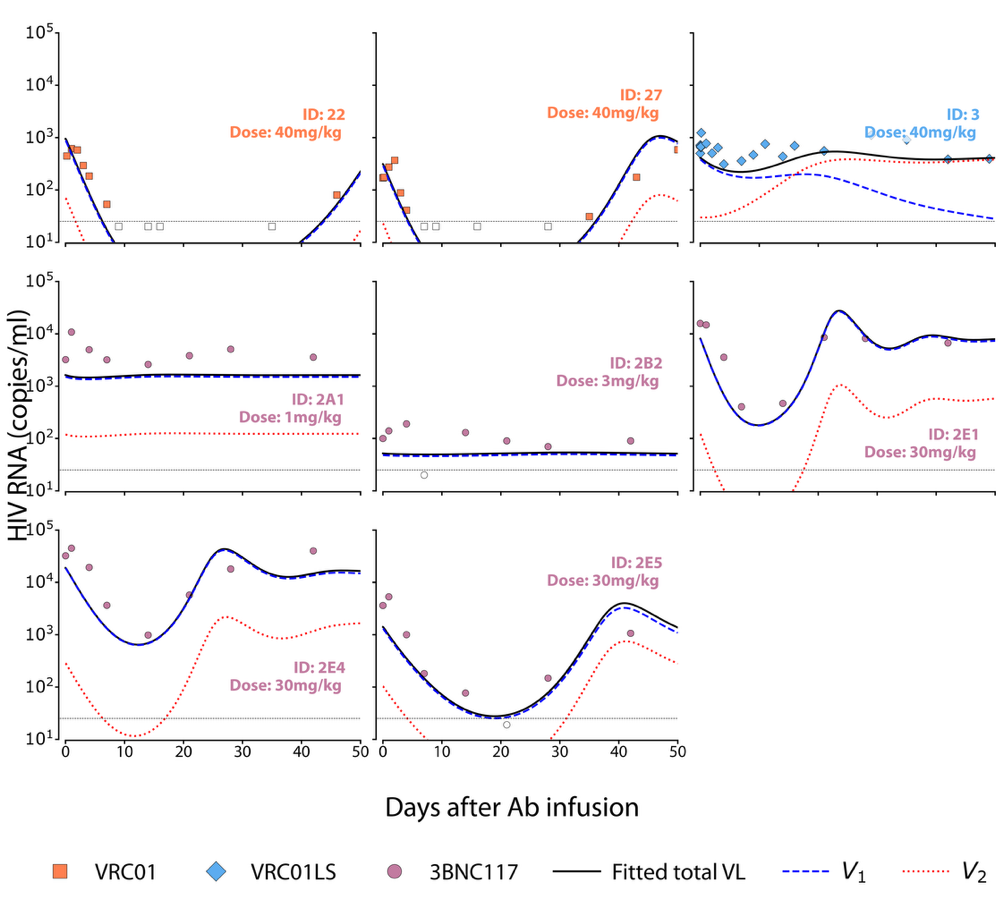

**Supplementary Table 6.** Summary results for the estimated proportion of *in vitro* IC50 measurements at pre- and post- infusion of bnAb ($\rho$). The remainder of model parameters were fixed at population level while allowing for random effects. **Supplementary Table 5** was used to fix the population mode.

| Parameter | Population Parameter (%RSE) | |
| --- | --- | --- |
| $\boldsymbol{\rho}$ | VRC01 group: | 1 (590) |
|  | VRC01-LS group: | 20 (35.1) |
|  | 3BNC117 group: | 14.05 (60.2) |
| FIXED Parameters with Random effects | | |
| $\boldsymbol{log}_{\boldsymbol{10}}\left( \boldsymbol{f}_{\boldsymbol{s}} \right)$ | Cluster 1: | -0.0066 |
|  | Cluster 2: | -0.033 |
| ${\boldsymbol{IC}\boldsymbol{50}}_{\boldsymbol{n}}$ | 17.52 | |
| $\boldsymbol{\omega}_{\boldsymbol{IC}\boldsymbol{50}}$ | 0.00013 | |
| $\boldsymbol{log}_{\boldsymbol{10}}\left( \boldsymbol{\beta} \right)$ | Cluster 1: | -5.29 |
|  | Cluster 2 & 3: | -4.07 |
| $\boldsymbol{\mu}$ | 0.011 | |
| $\boldsymbol{\xi}$ | VRC01 group: | 0.37 |
|  | VRC01-LS group: | 0.093 |
|  | 3BNC117 group: | 0.14 |
| $\boldsymbol{\pi}$ | Cluster 1: | 2767.74 (fixed with random effects) |
|  | Cluster 2: | 411.82 (fixed with random effects) |
| $\boldsymbol{\alpha}$ | 60.05 (20.41) | |
| $\boldsymbol{\delta}_{\boldsymbol{S}}$ (fixed no random effects) | 0.01 | |
| $\boldsymbol{\gamma}$ (fixed no random effects) | 23 | |
| $\boldsymbol{\delta}_{\boldsymbol{I}}$ | 0.5 | |
| $\boldsymbol{\Phi}$ (fixed no random effects) | 0.05 | |
| $\boldsymbol{h}$ (fixed no random effects) | 1.1 | |

**Supplementary Figure 2.** **Simplified modeling of VRC07-523-LS with and without Fc-functions in non-human primates suggests potent Fc-mediated lysis of infected cells.** Data were digitally extracted from Asokan et al. PNAS 2020. Left: bnAb pharmacokinetics (PK) for wild type (wt) and two engineered mutant forms of this bnAb without Fc-function (lala) and with *in vitro* enhancement to Fc-function (del). Data are dots and line is biphasic decay PK model. Right: Mean viral kinetics (dots) and viral load model fits (lines) using a simplified version of the best model selected for the human data. control and placebo data represent no infusion and infusion of sham bnAb so should remain stable.

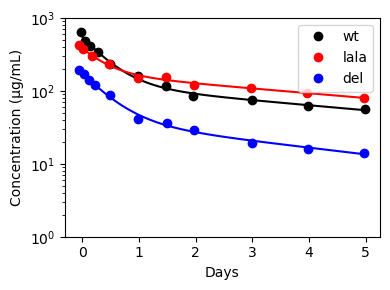

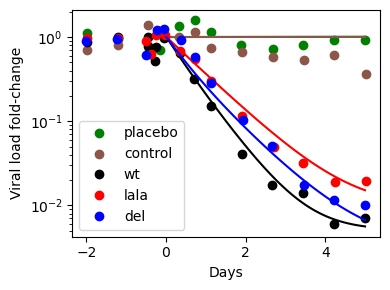

**Supplementary Figure 3.** **Estimates of the percentage of potentially resistant minor variants prior to bnAb treatment.** The initial fraction of viral load in strain 2. By definition of minor variant, values were constrained to not go above 40%.

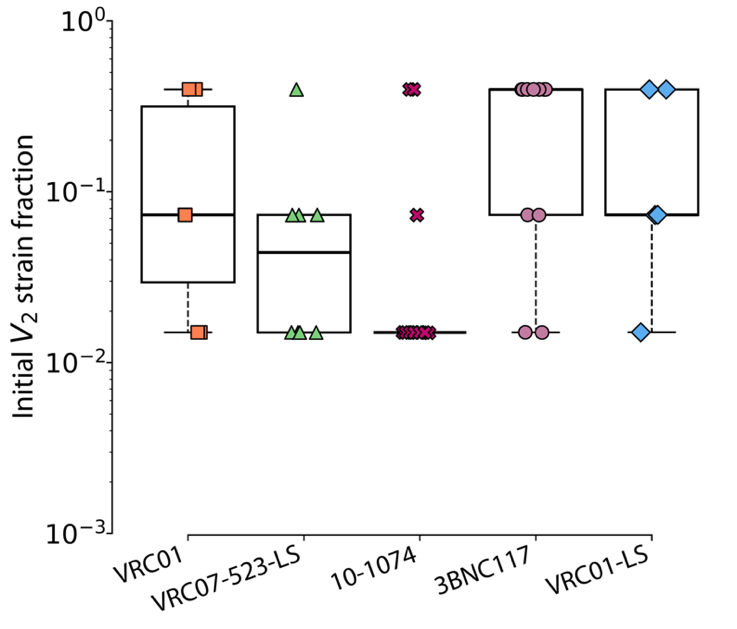

**Supplementary Table 7.** Biological description and units of the parameters included in model (4).

| **Parameter** | **Description** | **Units** |
| --- | --- | --- |
| $f_{s}$ | Proportion of abundant strains ($V_{1}$) at $t_{0}$. | unitless |
| $\alpha$ | Generation rate of susceptible cells | $\frac{cell}{time}$ |
| $\delta_{s}$ | Natural death rate of susceptible cells | ${time}^{-1}$ |
| $\beta_{S}$ | Infection rate of susceptible cells. | ${(virus\cdot time)}^{-1}$ |
| $\beta_{V}$ | Anchoring rate of virus | ${(cell\cdot time)}^{-1}$ |
| $\delta_{I}$ | Natural death rate of virus producing infected cells | ${time}^{-1}$ |
| $\pi$ | Rate of viremia replication | $\frac{virus}{cell \cdot time}$ |
| $\gamma$ | Natural death rate of virus | ${time}^{-1}$ |
| $\mu$ | Proportion of infected cells that are long-lived cells | unitless |
| $\xi$ | Intrinsic Fc-effector function activation rate | ${time}^{-1}$ |
| $\Phi$ | Conversion rate from long-lived infected cells to virus producing cells. | ${time}^{-1}$ |
| $x_{n}$ | *In vivo* IC50 for neutralization (${IC50}_{n}$) | $\frac{\mu g}{ml}$ |
| $\omega_{IC50}$ | Log-scale ratio between ${IC50}_{n}$ and *in vivo* IC50 for Fc functions (${EC50}_{e}$) | unitless |
| $\rho$ | Proportion of *in vitro* IC50 measurements at two time points. | unitless |
| $h$ | Hill exponent from the *in vivo* neutralization curve | unitless |
