## Supplementary Data 3 for "Modeling quantifies in vivo neutralization, Fc-mediated killing, and resistance in human clinical trials of five anti-HIV broadly neutralizing antibodies"

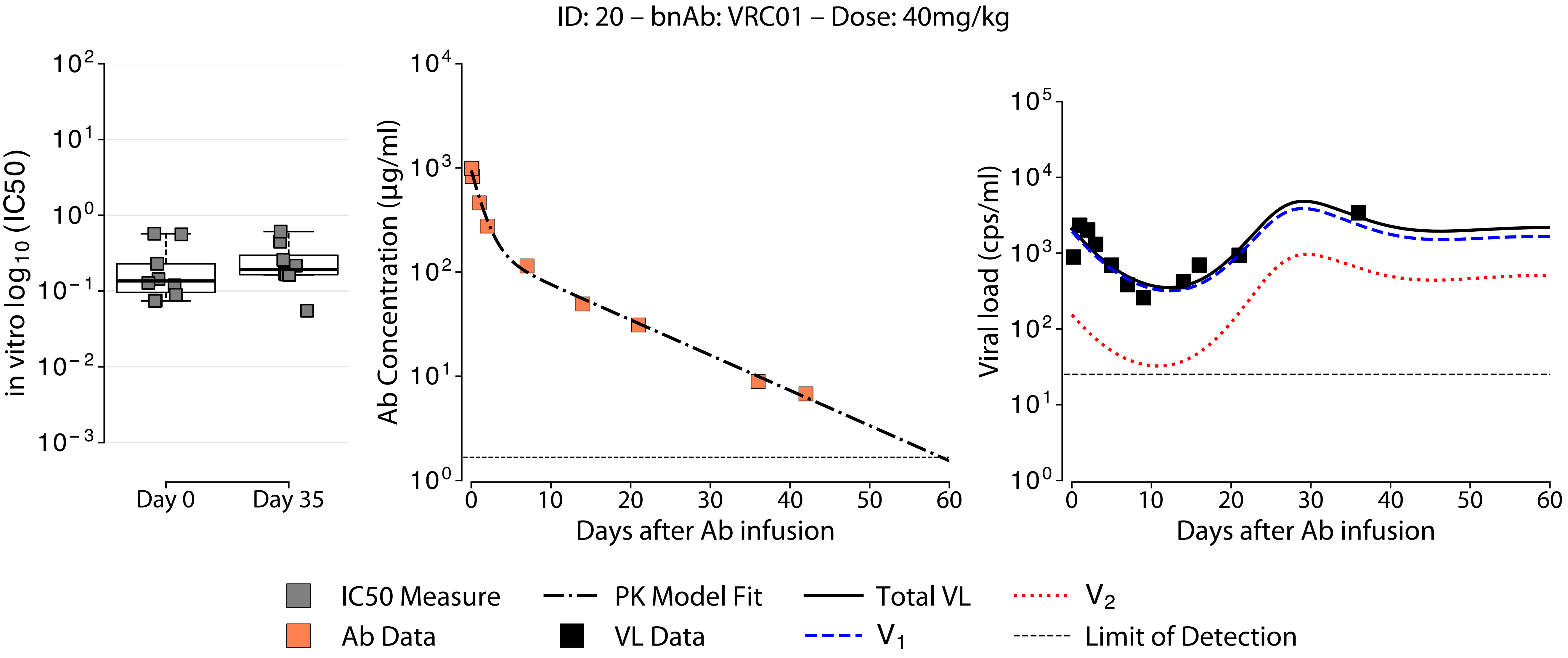
ID: 20 – bnAb: VRC01 – Dose: 40mg/kg
in vitro log10 (IC50)
Day 0
Day 35
Ab Concentration (µg/ml)
Days after Ab infusion
Viral load (cps/ml)
Days after Ab infusion
IC50 Measure
Ab Data
PK Model Fit
VL Data
Total VL
V1
V2
Limit of Detection

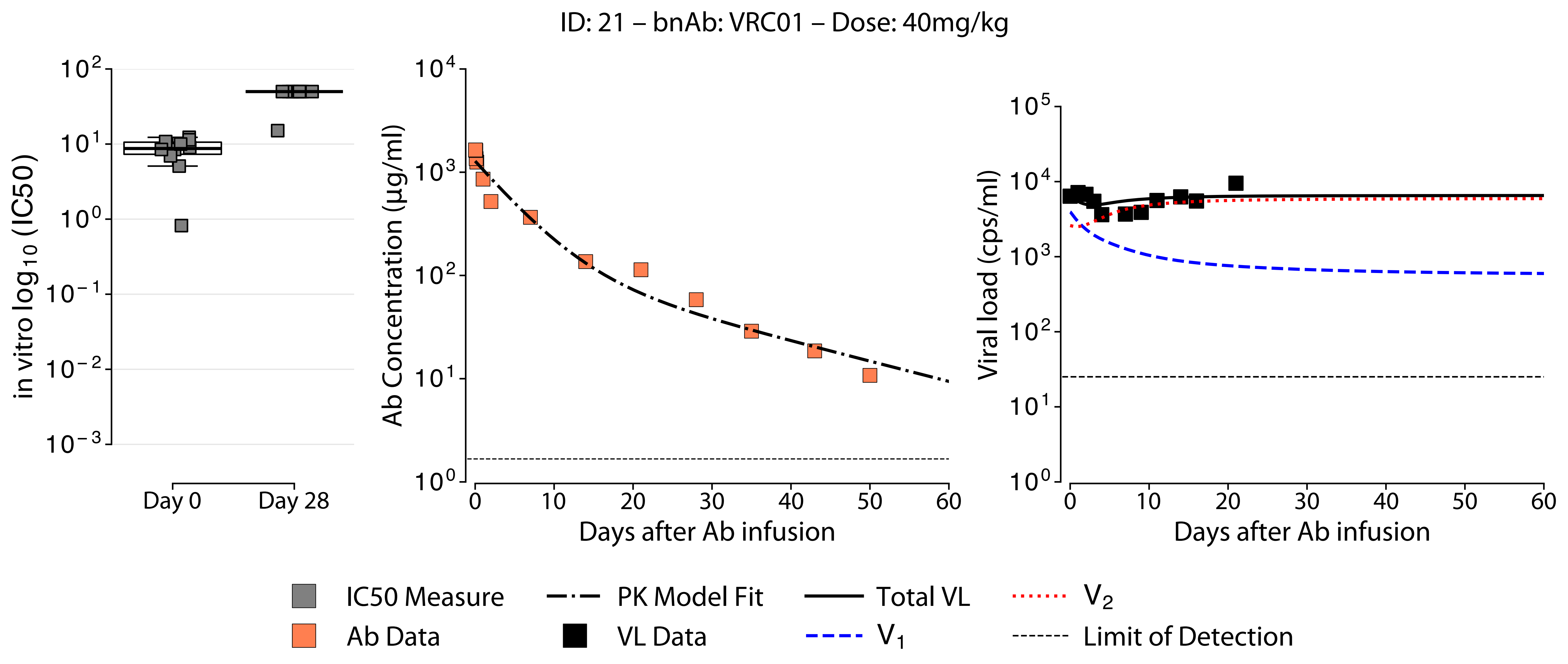

ID: 21 – bnAb: VRC01 – Dose: 40mg/kg
in vitro log10 (IC50)
Day 0
Day 28
Ab Concentration (µg/ml)
Days after Ab infusion
Viral load (cps/ml)
Days after Ab infusion
IC50 Measure
Ab Data
PK Model Fit
VL Data
Total VL
V1
V2
Limit of Detection

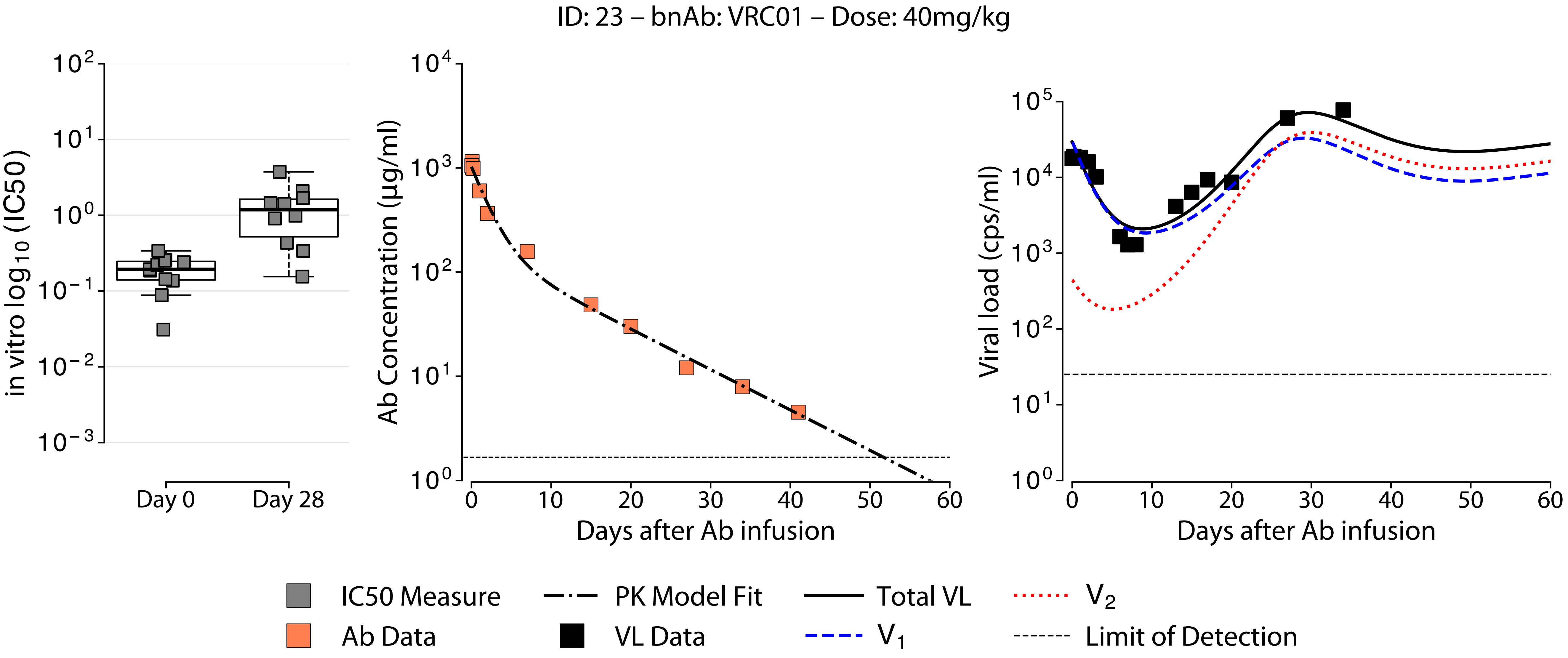
ID: 23 – bnAb: VRC01 – Dose: 40mg/kg
in vitro log10 (IC50)
Day 0
Day 28
Ab Concentration (µg/ml)
Days after Ab infusion
Viral load (cps/ml)
Days after Ab infusion
IC50 Measure
Ab Data
PK Model Fit
VL Data
Total VL
V1
V2
Limit of Detection

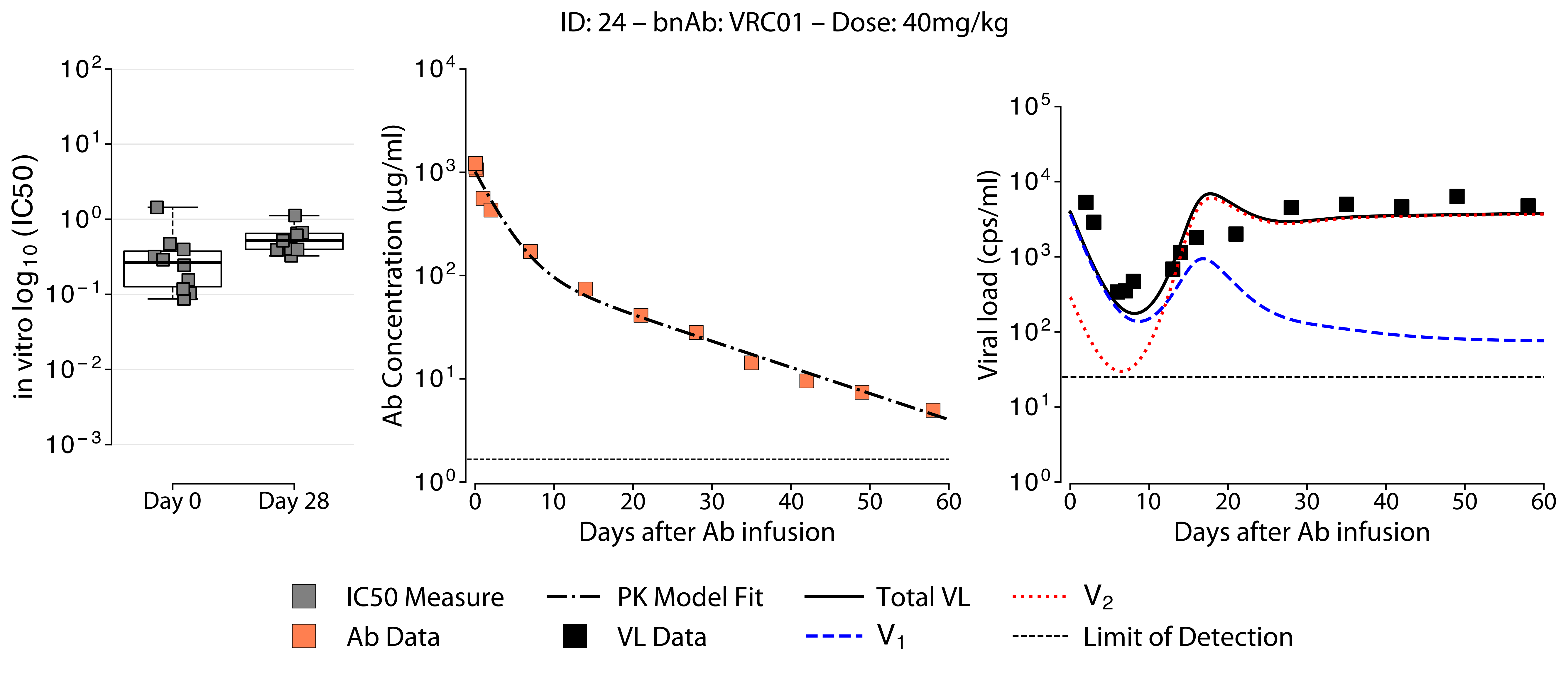

ID: 24 – bnAb: VRC01 – Dose: 40mg/kg
in vitro log10 (IC50)
Day 0
Day 28
Ab Concentration (µg/ml)
Days after Ab infusion
Viral load (cps/ml)
Days after Ab infusion
IC50 Measure
Ab Data
PK Model Fit
VL Data
Total VL
V1
V2
Limit of Detection

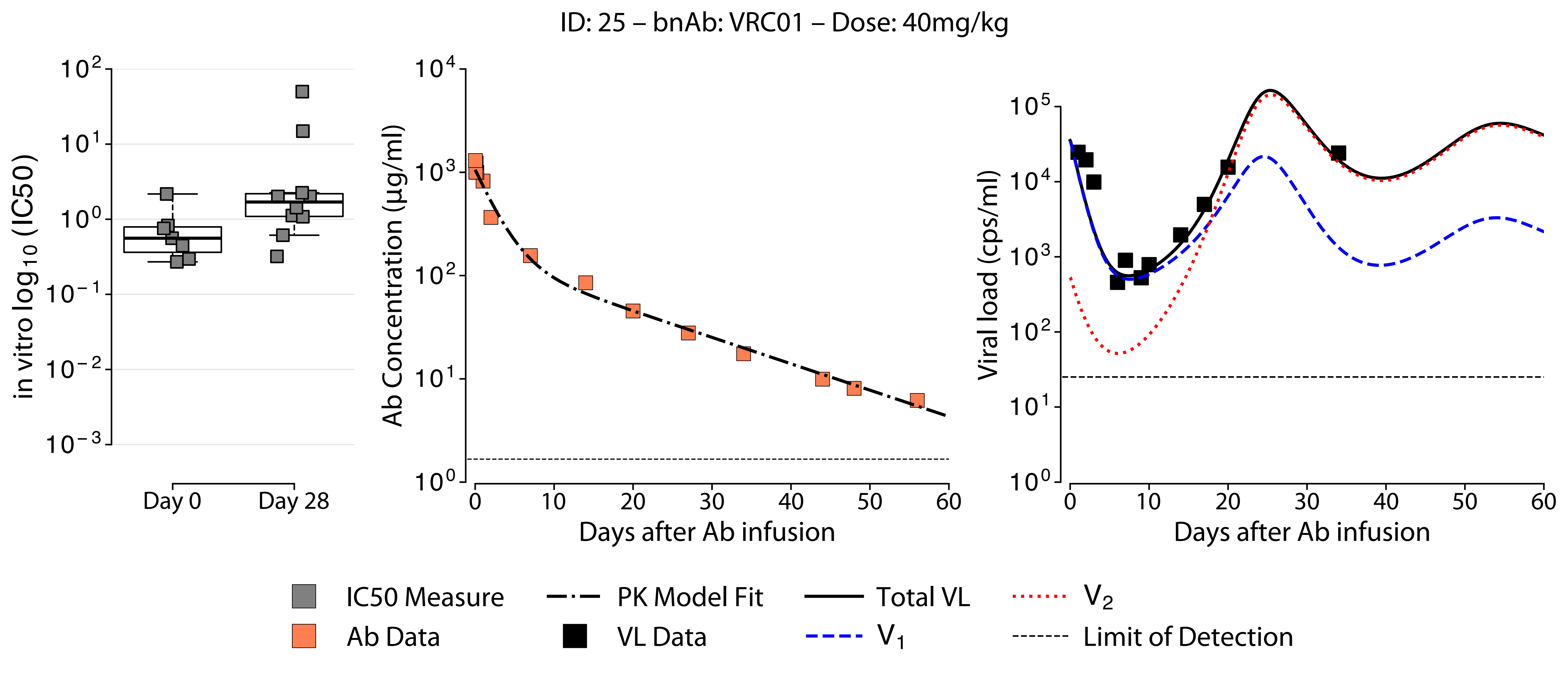

ID: 25 – bnAb: VRC01 – Dose: 40mg/kg
in vitro log10 (IC50)
Day 0
Day 28
Ab Concentration (µg/ml)
Days after Ab infusion
Viral load (cps/ml)
Days after Ab infusion
IC50 Measure
Ab Data
PK Model Fit
VL Data
Total VL
V1
V2
Limit of Detection

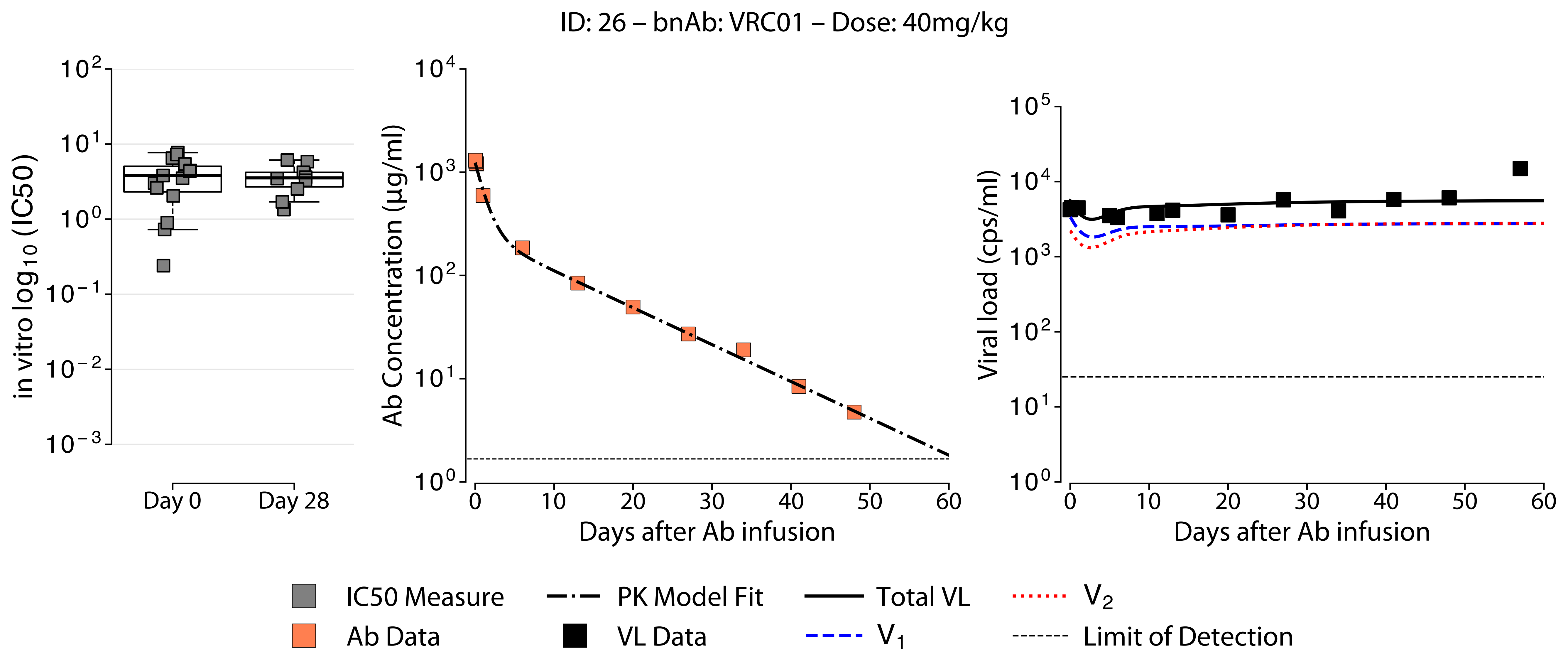
ID: 26 – bnAb: VRC01 – Dose: 40mg/kg
in vitro $\log_{10}$ (IC50)
$10^{2}$
$10^{1}$
$10^{0}$
$10^{-1}$
$10^{-2}$
$10^{-3}$
Day 0
Day 28
Ab Concentration (μg/ml)
$10^{4}$
$10^{3}$
$10^{2}$
$10^{1}$
$10^{0}$
0 10 20 30 40 50 60
Days after Ab infusion
Viral load (cps/ml)
$10^{5}$
$10^{4}$
$10^{3}$
$10^{2}$
$10^{1}$
$10^{0}$
0 10 20 30 40 50 60
Days after Ab infusion
IC50 Measure
Ab Data
PK Model Fit
VL Data
Total VL
$V_1$
$V_2$
Limit of Detection

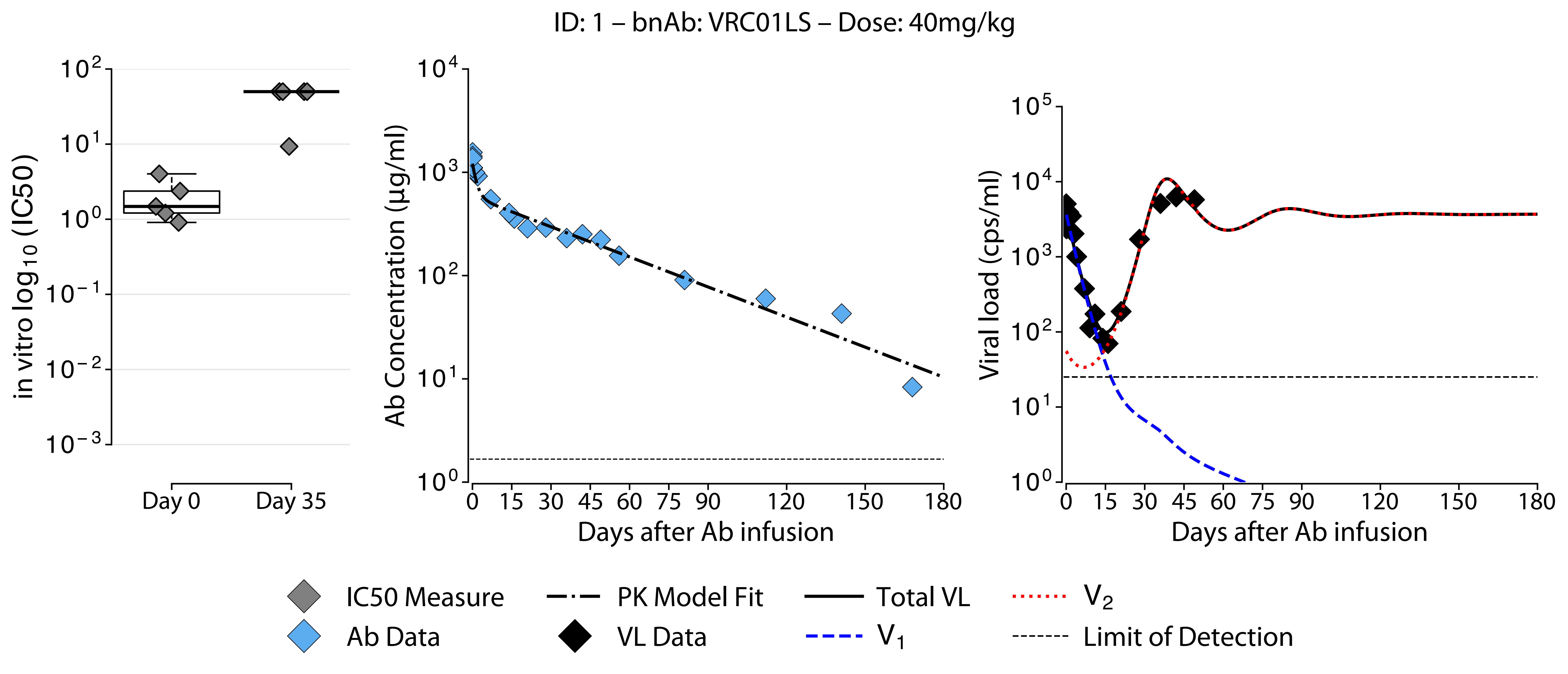
ID: 1 – bnAb: VRC01LS – Dose: 40mg/kg
in vitro $\log_{10}$ (IC50)
$10^2$
$10^1$
$10^0$
$10^{-1}$
$10^{-2}$
$10^{-3}$
Day 0
Day 35
Ab Concentration (μg/ml)
$10^4$
$10^3$
$10^2$
$10^1$
$10^0$
0 15 30 45 60 75 90 120 150 180
Days after Ab infusion
Viral load (cps/ml)
$10^5$
$10^4$
$10^3$
$10^2$
$10^1$
$10^0$
0 15 30 45 60 75 90 120 150 180
Days after Ab infusion
IC50 Measure
Ab Data
PK Model Fit
VL Data
Total VL
$V_1$
$V_2$
Limit of Detection

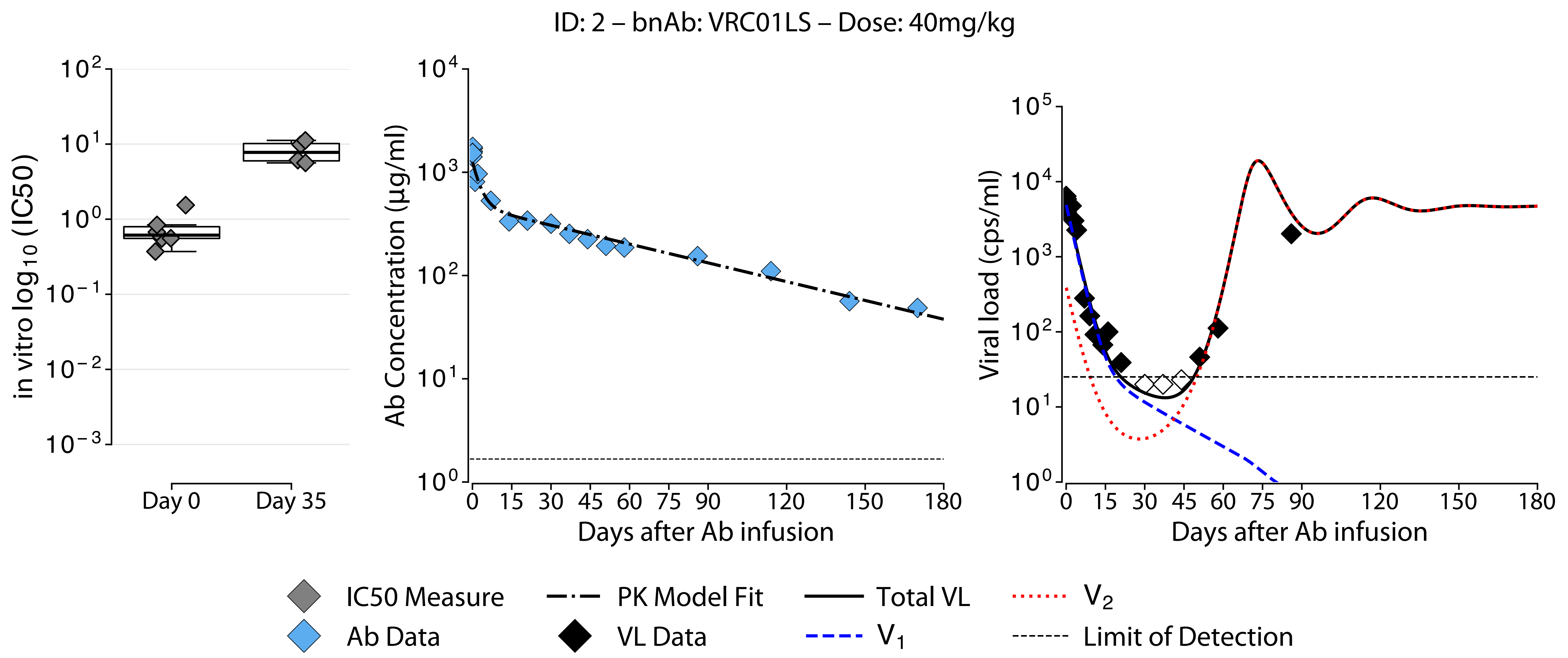

ID: 2 – bnAb: VRC01LS – Dose: 40mg/kg
in vitro log10 (IC50)
Day 0
Day 35
Ab Concentration (µg/ml)
Days after Ab infusion
Viral load (cps/ml)
Days after Ab infusion
IC50 Measure
Ab Data
PK Model Fit
VL Data
Total VL
V1
V2
Limit of Detection

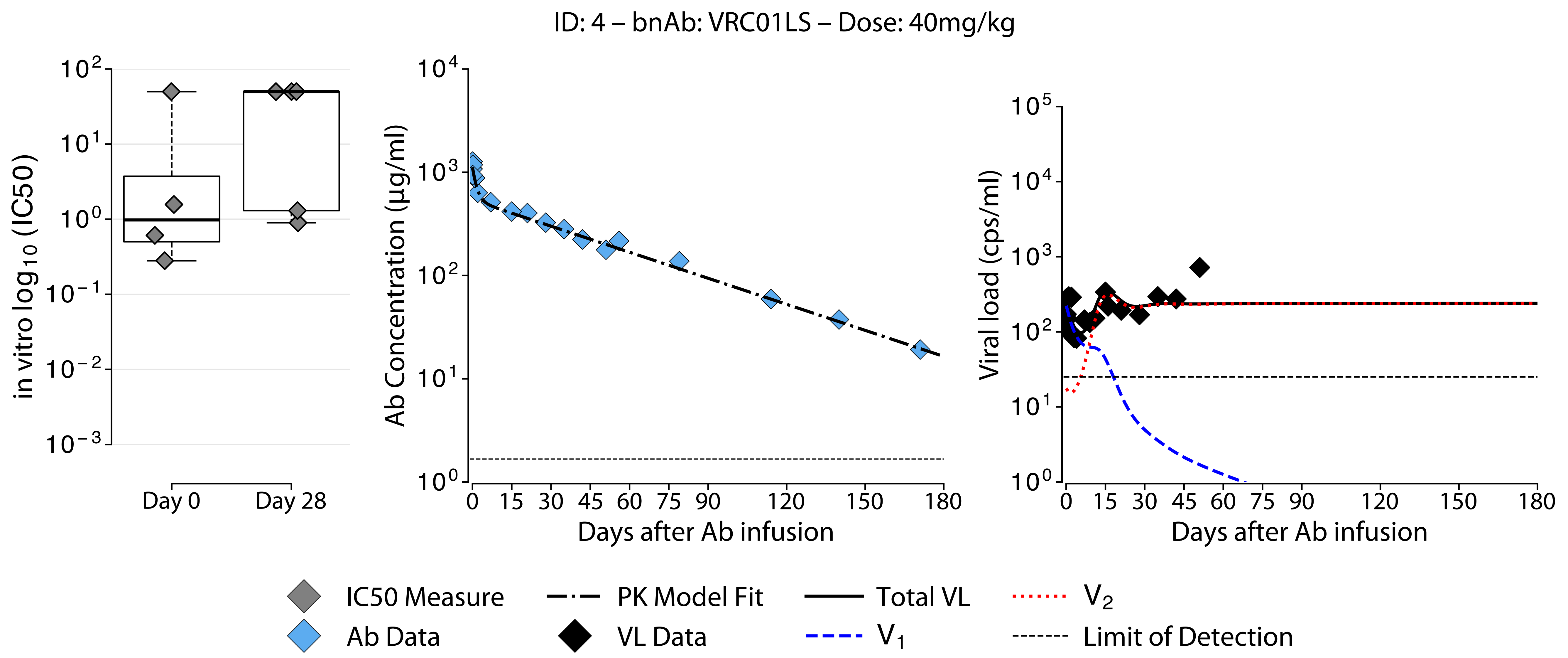
ID: 4 – bnAb: VRC01LS – Dose: 40mg/kg
in vitro log10 (IC50)
Day 0
Day 28
Ab Concentration (µg/ml)
Days after Ab infusion
Viral load (cps/ml)
Days after Ab infusion
IC50 Measure
Ab Data
PK Model Fit
VL Data
Total VL
V1
V2
Limit of Detection

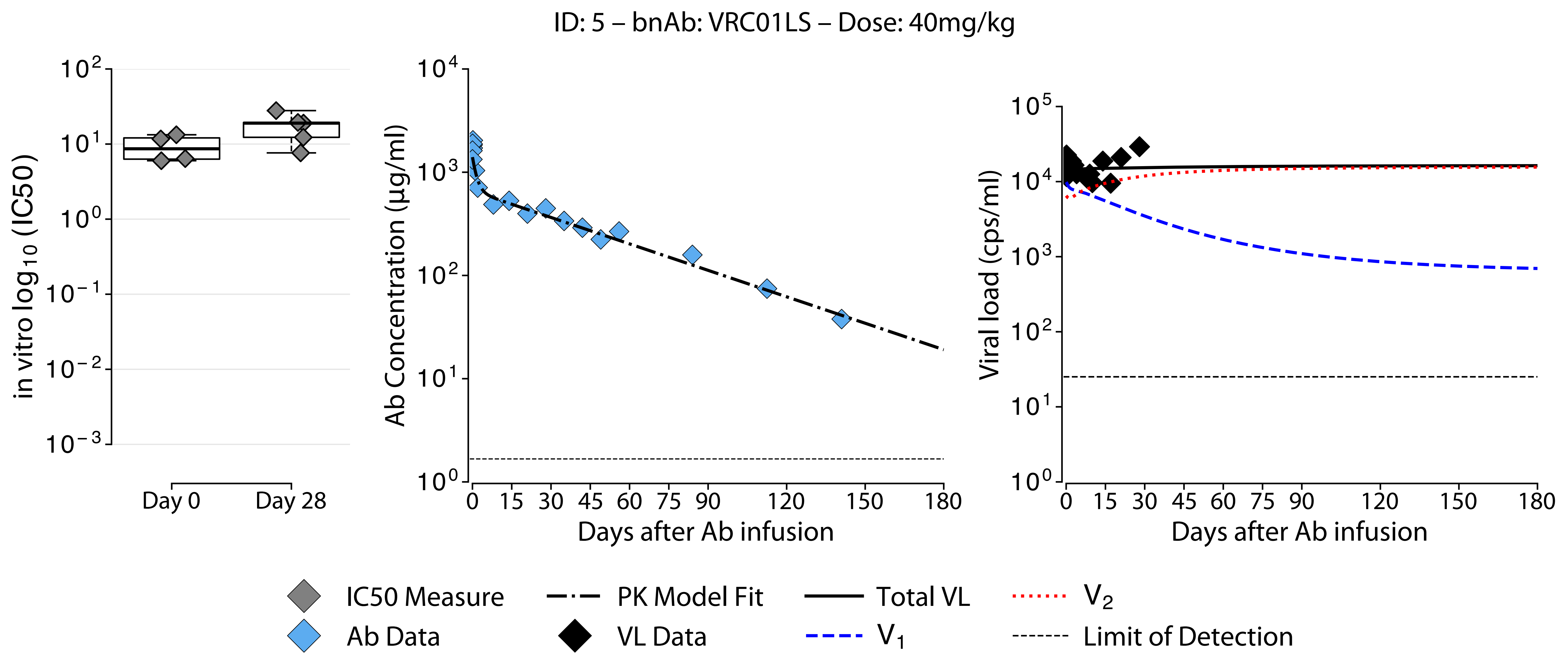

ID: 5 – bnAb: VRC01LS – Dose: 40mg/kg
in vitro log10 (IC50)
$10^2$
$10^1$
$10^0$
$10^{-1}$
$10^{-2}$
$10^{-3}$
Day 0
Day 28
Ab Concentration (µg/ml)
$10^4$
$10^3$
$10^2$
$10^1$
$10^0$
0 15 30 45 60 75 90 120 150 180
Days after Ab infusion
Viral load (cps/ml)
$10^5$
$10^4$
$10^3$
$10^2$
$10^1$
$10^0$
0 15 30 45 60 75 90 120 150 180
Days after Ab infusion
IC50 Measure
Ab Data
PK Model Fit
VL Data
Total VL
$V_1$
$V_2$
Limit of Detection

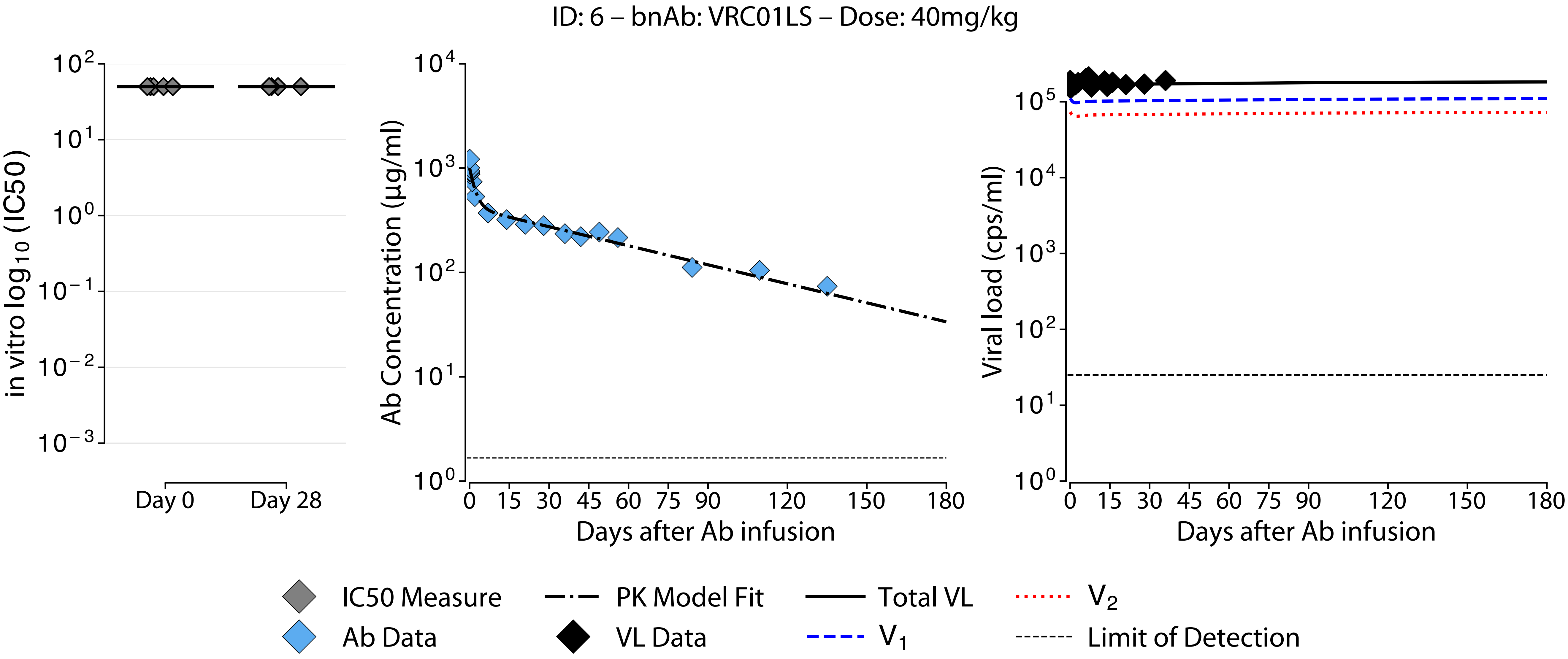
ID: 6 – bnAb: VRC01LS – Dose: 40mg/kg
in vitro log10 (IC50)
Day 0
Day 28
Ab Concentration (μg/ml)
Days after Ab infusion
Viral load (cps/ml)
Days after Ab infusion
IC50 Measure
Ab Data
PK Model Fit
VL Data
Total VL
V1
V2
Limit of Detection

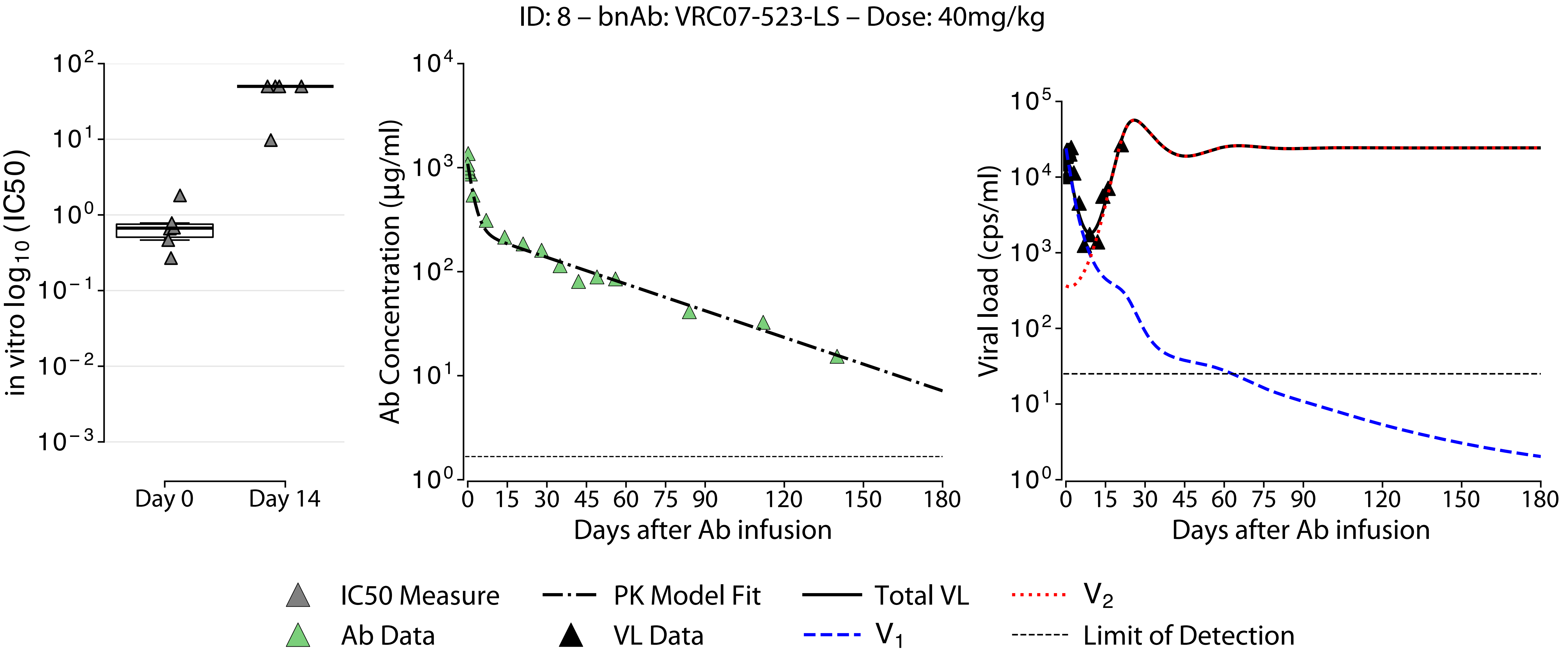

ID: 8 – bnAb: VRC07-523-LS – Dose: 40mg/kg
in vitro $\log_{10}$ (IC50)
$10^{2}$
$10^{1}$
$10^{0}$
$10^{-1}$
$10^{-2}$
$10^{-3}$
Day 0
Day 14
Ab Concentration (μg/ml)
$10^{4}$
$10^{3}$
$10^{2}$
$10^{1}$
$10^{0}$
0 15 30 45 60 75 90 120 150 180
Days after Ab infusion
Viral load (cps/ml)
$10^{5}$
$10^{4}$
$10^{3}$
$10^{2}$
$10^{1}$
$10^{0}$
0 15 30 45 60 75 90 120 150 180
Days after Ab infusion
IC50 Measure
Ab Data
PK Model Fit
VL Data
Total VL
$V_1$
$V_2$
Limit of Detection

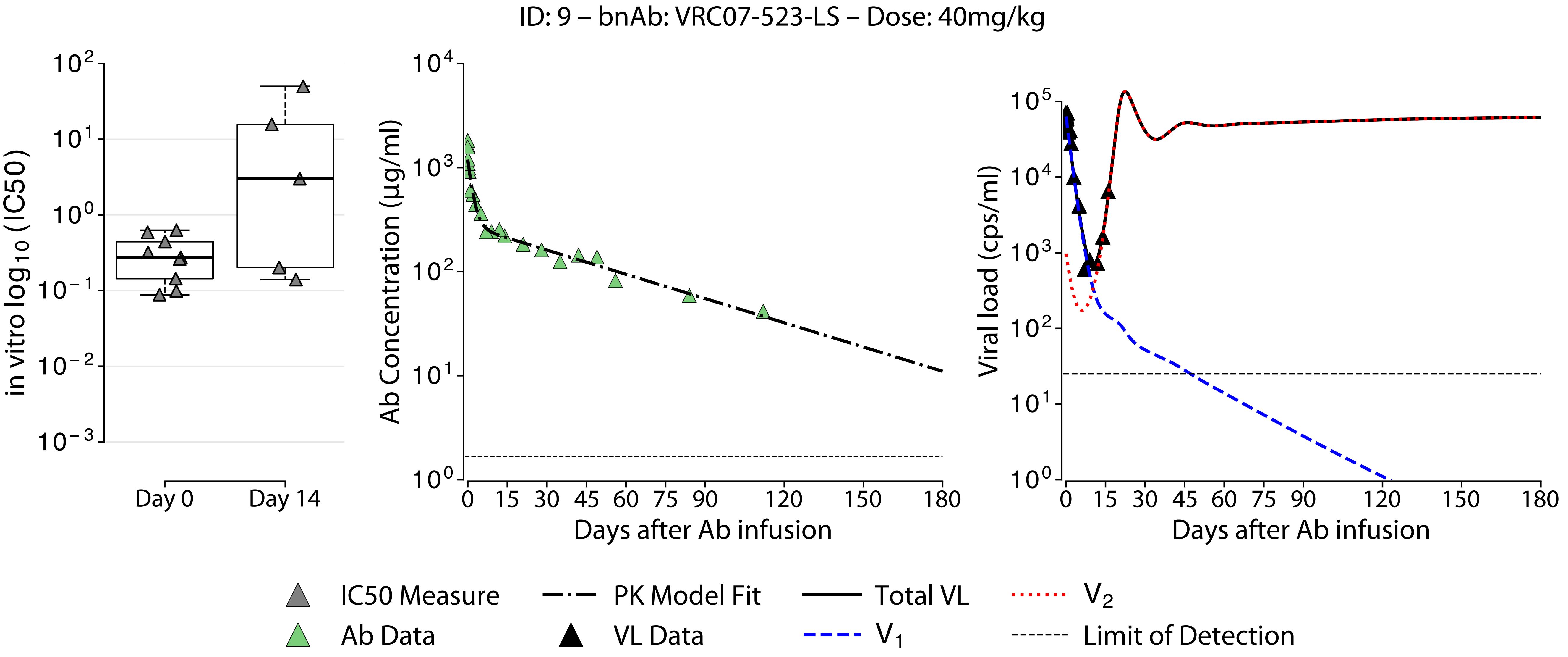

ID: 9 – bnAb: VRC07-523-LS – Dose: 40mg/kg
in vitro log10 (IC50)
Day 0
Day 14
Ab Concentration (μg/ml)
Days after Ab infusion
Viral load (cps/ml)
Days after Ab infusion
IC50 Measure
Ab Data
PK Model Fit
VL Data
Total VL
V1
V2
Limit of Detection

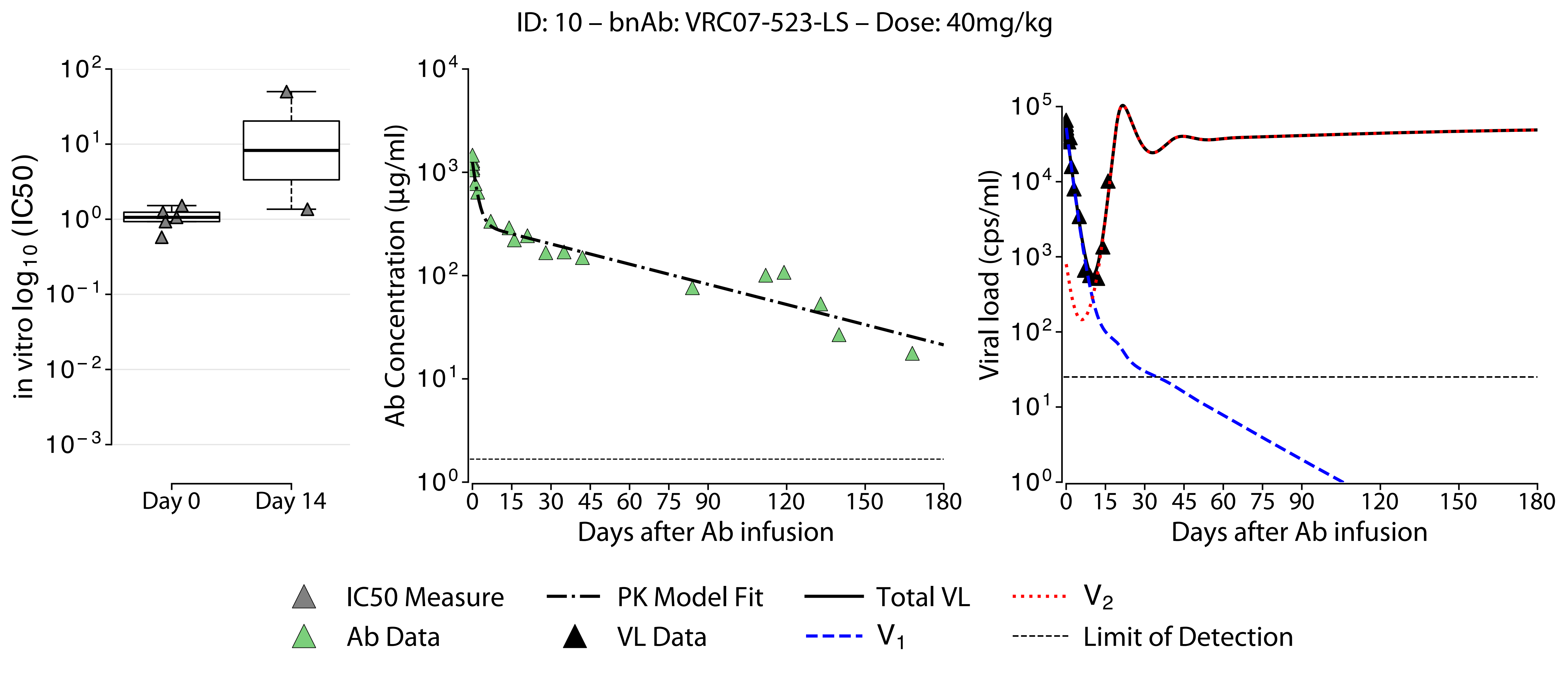

ID: 10 – bnAb: VRC07-523-LS – Dose: 40mg/kg
in vitro log10 (IC50)
Day 0
Day 14
Ab Concentration (µg/ml)
Days after Ab infusion
Viral load (cps/ml)
Days after Ab infusion
IC50 Measure
Ab Data
PK Model Fit
VL Data
Total VL
V1
V2
Limit of Detection

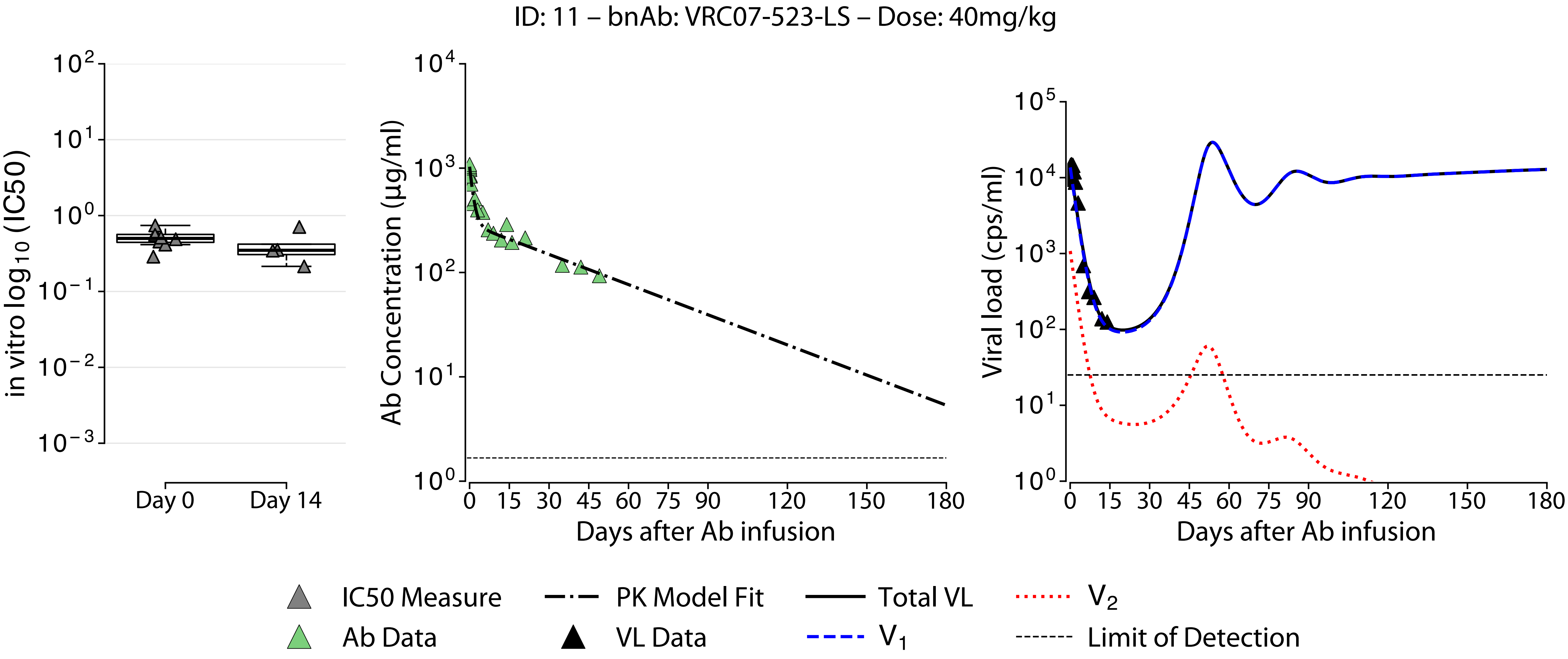
ID: 11 – bnAb: VRC07-523-LS – Dose: 40mg/kg
in vitro log10 (IC50)
Day 0
Day 14
Ab Concentration (μg/ml)
Days after Ab infusion
Viral load (cps/ml)
Days after Ab infusion
IC50 Measure
Ab Data
PK Model Fit
VL Data
Total VL
V1
V2
Limit of Detection

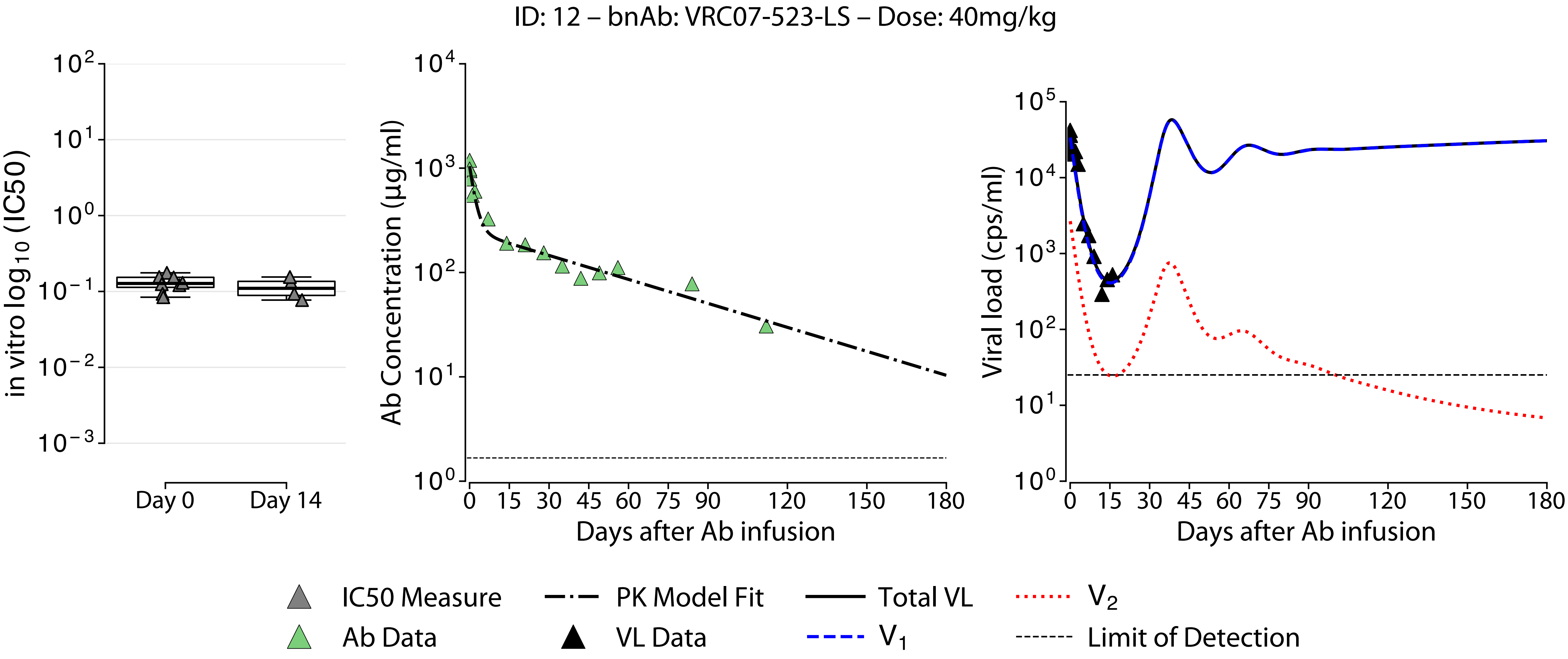

ID: 12 – bnAb: VRC07-523-LS – Dose: 40mg/kg
in vitro log10 (IC50)
Day 0
Day 14
Ab Concentration (µg/ml)
Days after Ab infusion
Viral load (cps/ml)
Days after Ab infusion
IC50 Measure
Ab Data
PK Model Fit
VL Data
Total VL
V1
V2
Limit of Detection

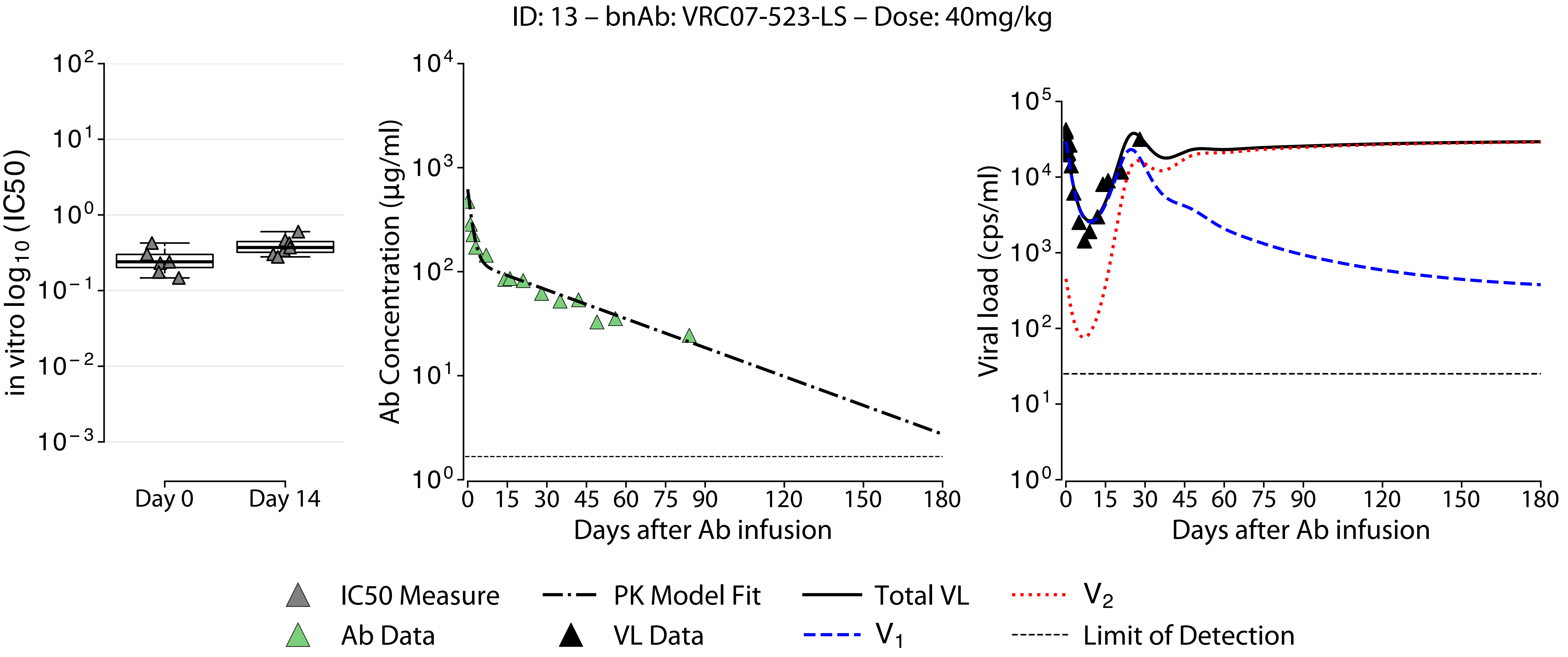
ID: 13 – bnAb: VRC07-523-LS – Dose: 40mg/kg
in vitro $\log_{10}$ (IC50)
Day 0
Day 14
Ab Concentration (μg/ml)
Days after Ab infusion
Viral load (cps/ml)
Days after Ab infusion
IC50 Measure
Ab Data
PK Model Fit
VL Data
Total VL
$V_1$
$V_2$
Limit of Detection

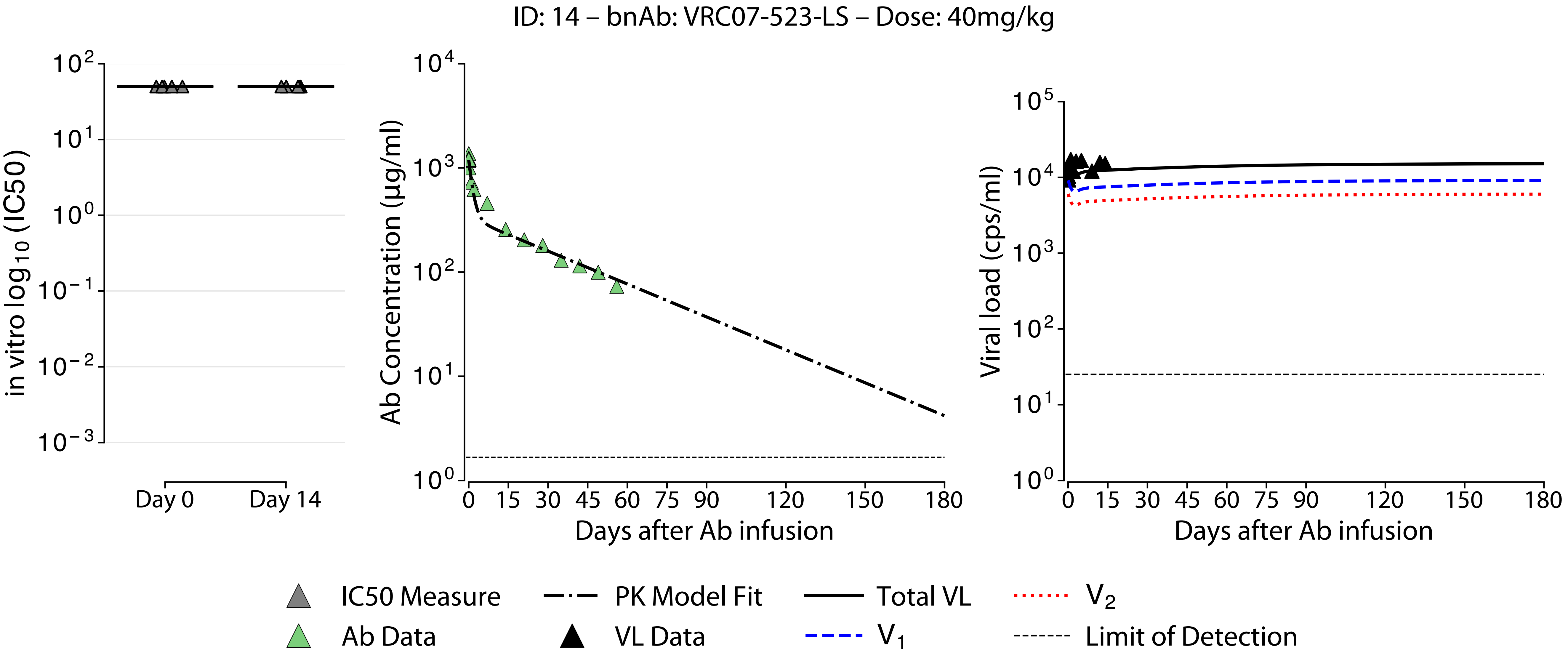

ID: 14 – bnAb: VRC07-523-LS – Dose: 40mg/kg
in vitro log10 (IC50)
Day 0
Day 14
Ab Concentration (μg/ml)
Days after Ab infusion
Viral load (cps/ml)
Days after Ab infusion
IC50 Measure
Ab Data
PK Model Fit
VL Data
Total VL
V1
V2
Limit of Detection

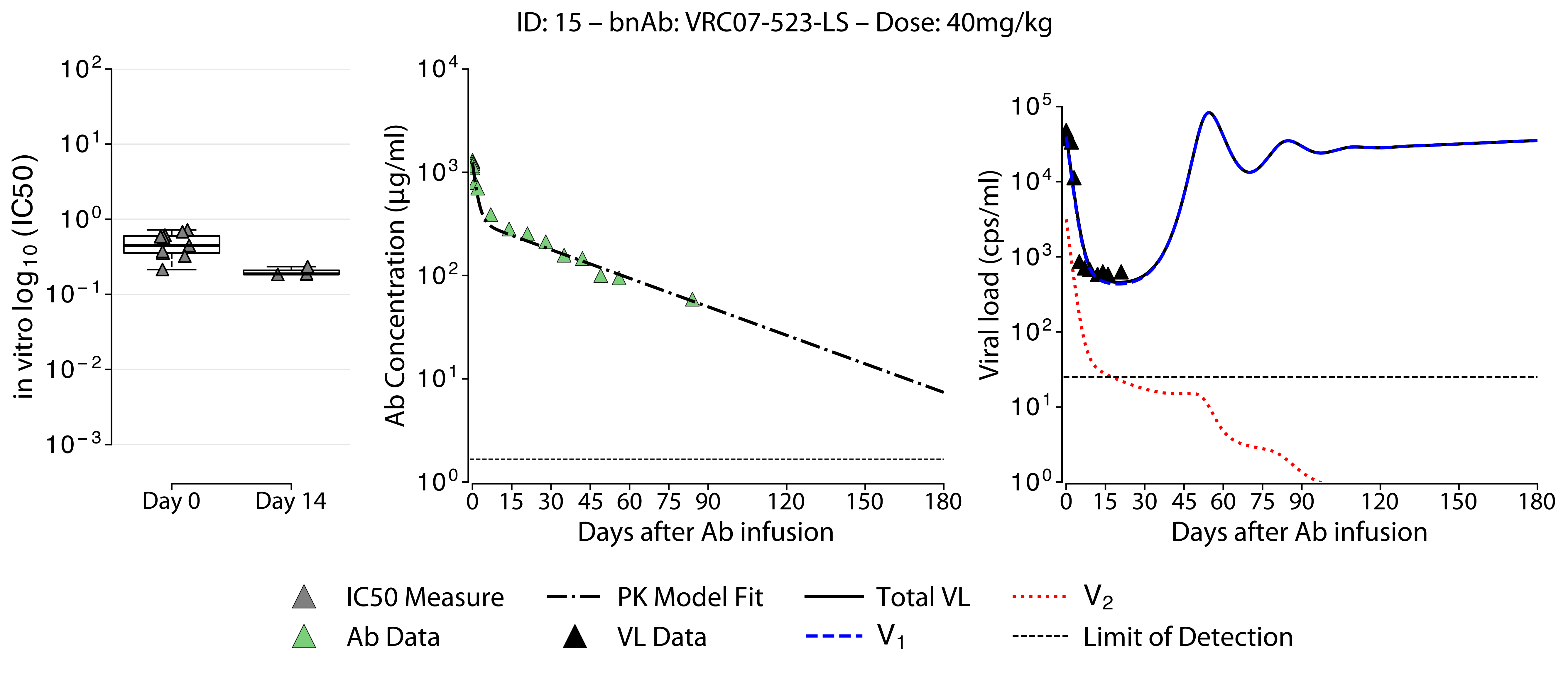

ID: 15 – bnAb: VRC07-523-LS – Dose: 40mg/kg
in vitro log10 (IC50)
Day 0
Day 14
Ab Concentration (µg/ml)
Days after Ab infusion
Viral load (cps/ml)
Days after Ab infusion
IC50 Measure
Ab Data
PK Model Fit
VL Data
Total VL
V1
V2
Limit of Detection

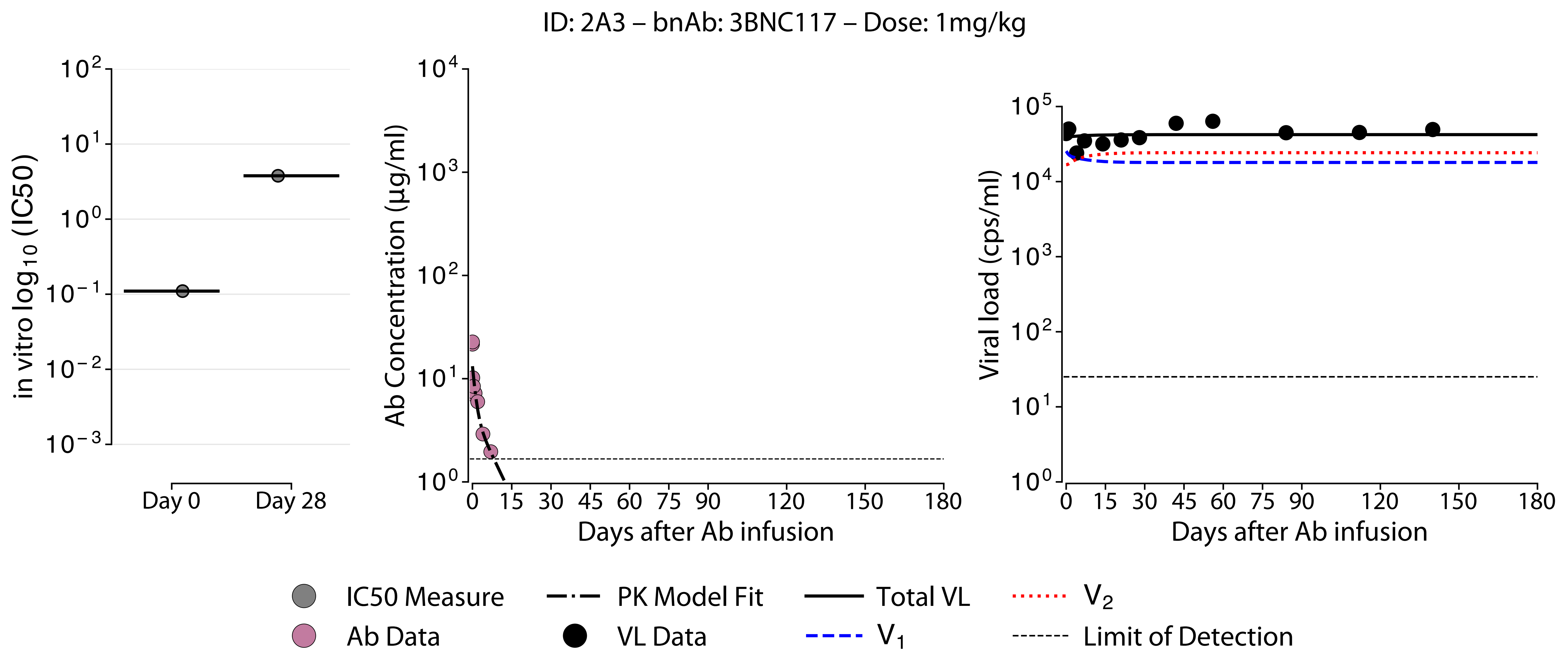
ID: 2A3 – bnAb: 3BNC117 – Dose: 1mg/kg
in vitro log10 (IC50)
Day 0
Day 28
Ab Concentration (μg/ml)
Days after Ab infusion
Viral load (cps/ml)
Days after Ab infusion
IC50 Measure
PK Model Fit
Total VL
V2
Ab Data
VL Data
V1
Limit of Detection

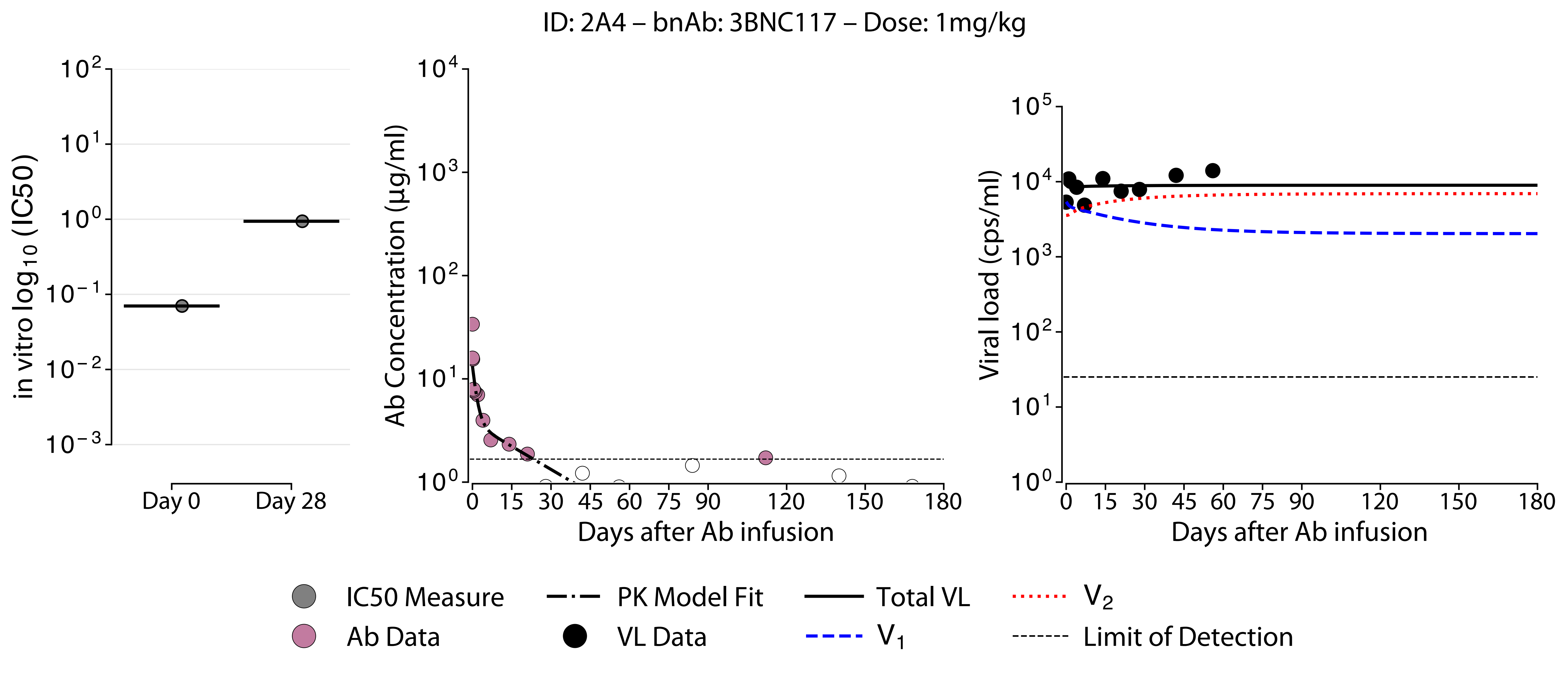
ID: 2A4 – bnAb: 3BNC117 – Dose: 1mg/kg
in vitro $\log_{10}$ (IC50)
$10^{2}$
$10^{1}$
$10^{0}$
$10^{-1}$
$10^{-2}$
$10^{-3}$
Day 0
Day 28
Ab Concentration (µg/ml)
$10^{4}$
$10^{3}$
$10^{2}$
$10^{1}$
$10^{0}$
0 15 30 45 60 75 90 120 150 180
Days after Ab infusion
Viral load (cps/ml)
$10^{5}$
$10^{4}$
$10^{3}$
$10^{2}$
$10^{1}$
$10^{0}$
0 15 30 45 60 75 90 120 150 180
Days after Ab infusion
IC50 Measure
Ab Data
PK Model Fit
VL Data
Total VL
$V_1$
$V_2$
Limit of Detection

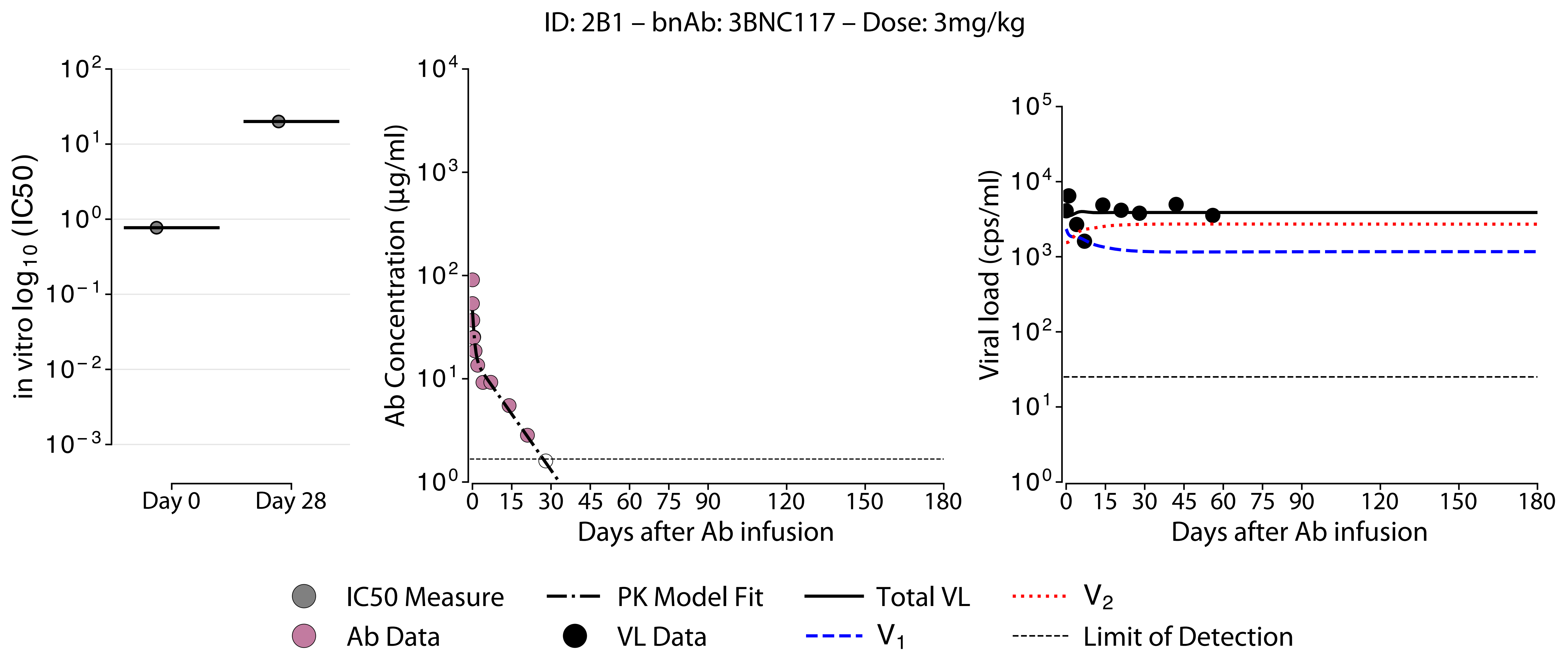

ID: 2B1 – bnAb: 3BNC117 – Dose: 3mg/kg
in vitro log$_{10}$ (IC50)
$10^2$
$10^1$
$10^0$
$10^{-1}$
$10^{-2}$
$10^{-3}$
Day 0
Day 28
Ab Concentration (μg/ml)
$10^4$
$10^3$
$10^2$
$10^1$
$10^0$
0 15 30 45 60 75 90 120 150 180
Days after Ab infusion
Viral load (cps/ml)
$10^5$
$10^4$
$10^3$
$10^2$
$10^1$
$10^0$
0 15 30 45 60 75 90 120 150 180
Days after Ab infusion
IC50 Measure
PK Model Fit
Total VL
$V_2$
Ab Data
VL Data
$V_1$
Limit of Detection

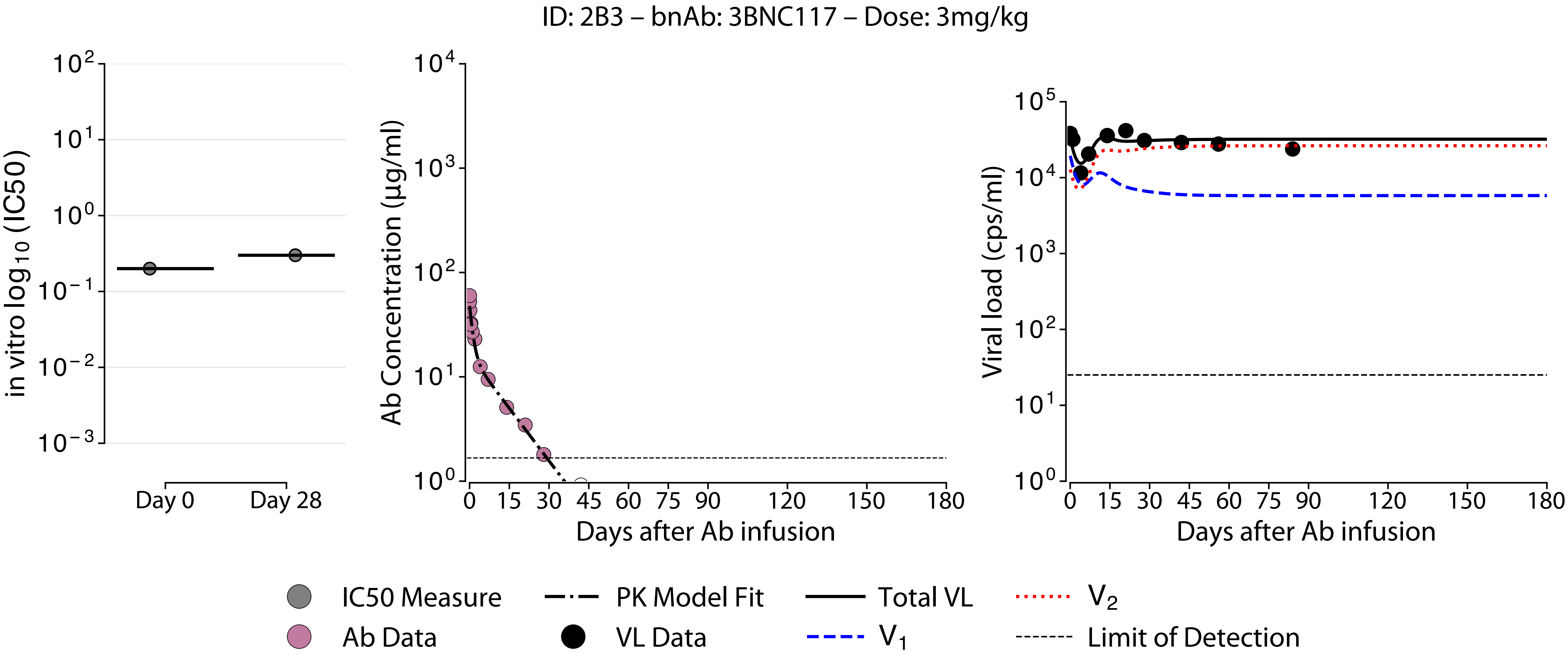
ID: 2B3 – bnAb: 3BNC117 – Dose: 3mg/kg
in vitro $\log_{10}$ (IC50)
$10^2$
$10^1$
$10^0$
$10^{-1}$
$10^{-2}$
$10^{-3}$
Day 0
Day 28
Ab Concentration (µg/ml)
$10^4$
$10^3$
$10^2$
$10^1$
$10^0$
0 15 30 45 60 75 90 120 150 180
Days after Ab infusion
Viral load (cps/ml)
$10^5$
$10^4$
$10^3$
$10^2$
$10^1$
$10^0$
0 15 30 45 60 75 90 120 150 180
Days after Ab infusion
IC50 Measure
PK Model Fit
Total VL
$V_2$
Ab Data
VL Data
$V_1$
Limit of Detection

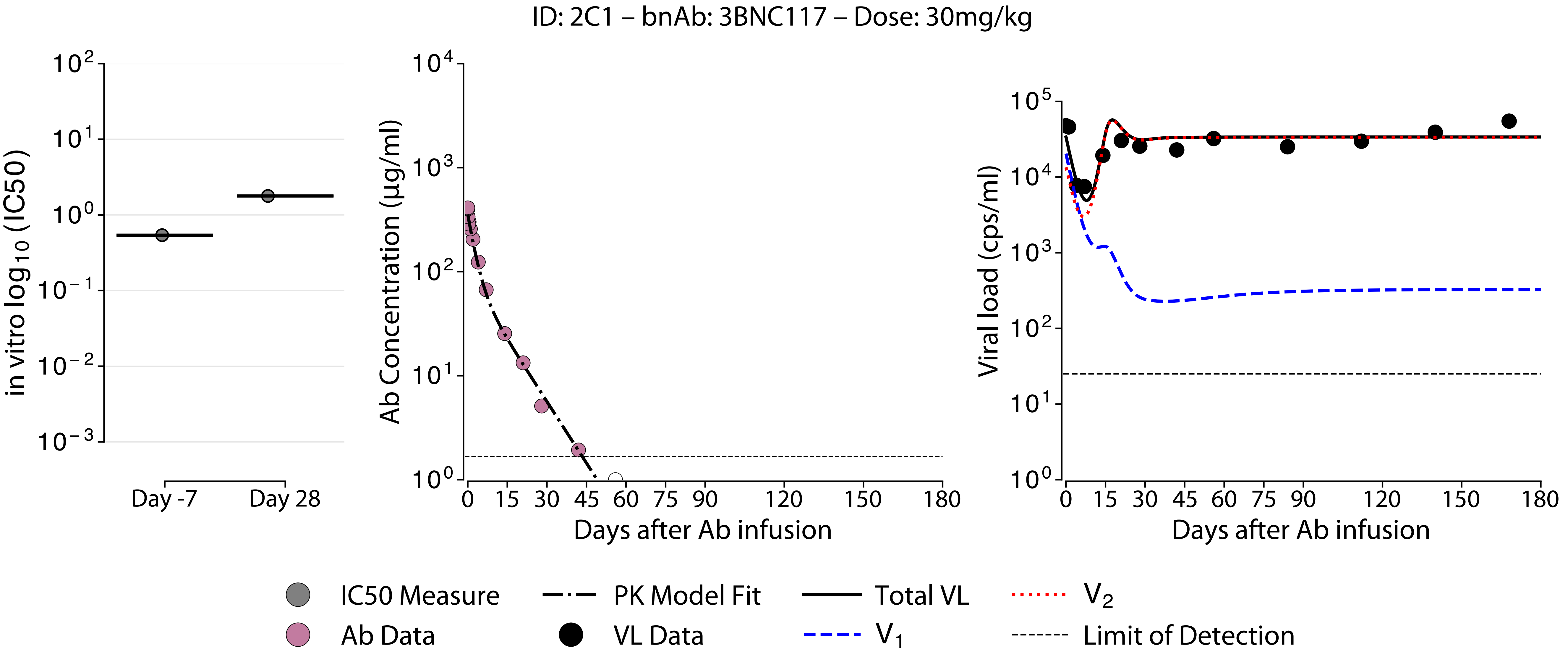
ID: 2C1 – bnAb: 3BNC117 – Dose: 30mg/kg
in vitro log10 (IC50)
Day -7
Day 28
Ab Concentration (μg/ml)
Days after Ab infusion
Viral load (cps/ml)
Days after Ab infusion
IC50 Measure
Ab Data
PK Model Fit
VL Data
Total VL
V1
V2
Limit of Detection

ID: 2C4 – bnAb: 3BNC117 – Dose: 10mg/kg

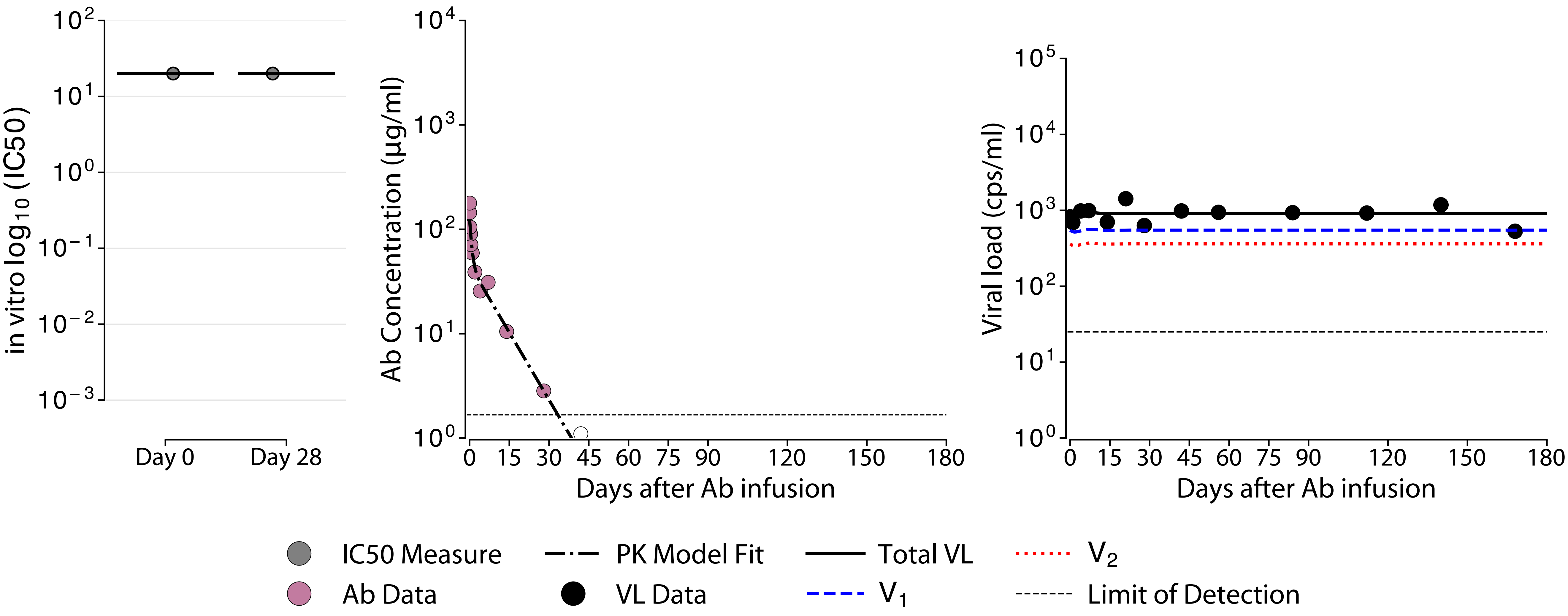

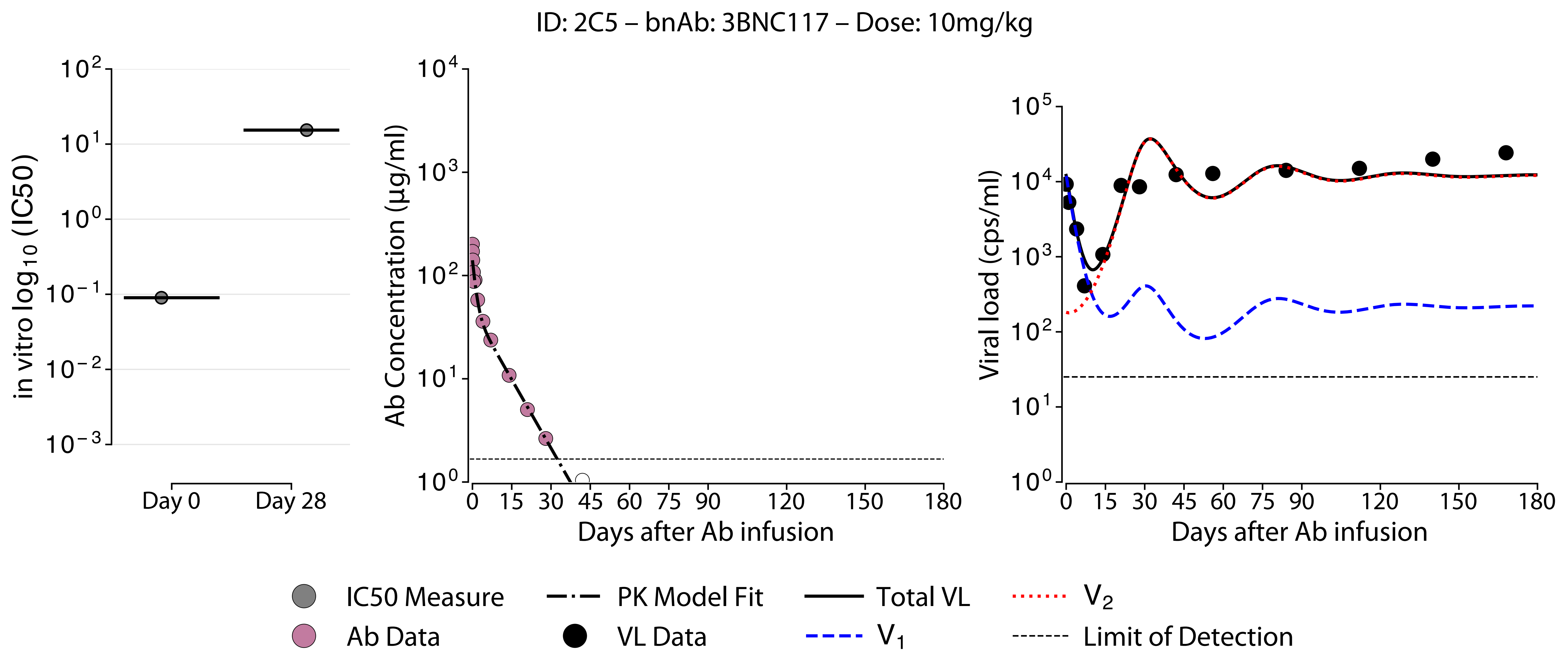

ID: 2C5 – bnAb: 3BNC117 – Dose: 10mg/kg
in vitro log10 (IC50)
Day 0
Day 28
Ab Concentration (µg/ml)
Days after Ab infusion
Viral load (cps/ml)
Days after Ab infusion
IC50 Measure
Ab Data
PK Model Fit
VL Data
Total VL
V1
V2
Limit of Detection

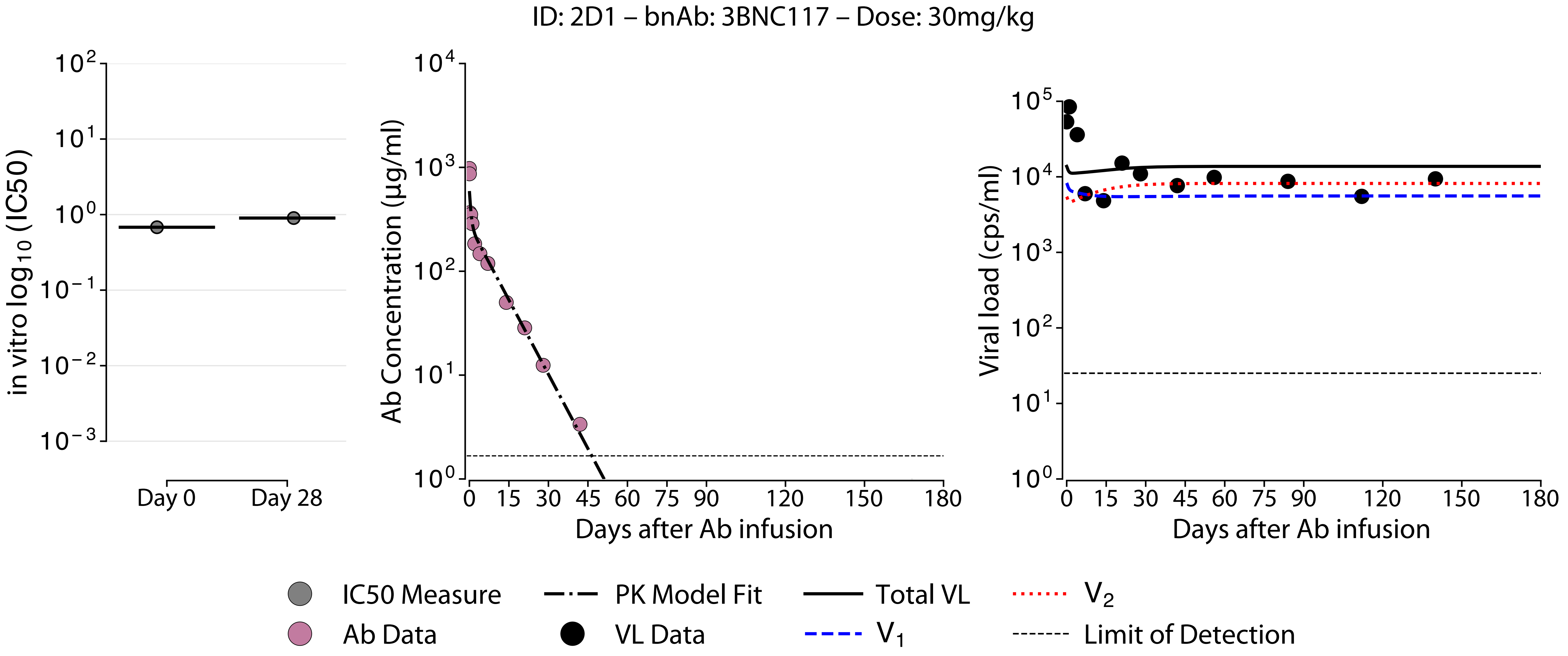

ID: 2D1 – bnAb: 3BNC117 – Dose: 30mg/kg
in vitro log10 (IC50)
Day 0
Day 28
Ab Concentration (μg/ml)
Days after Ab infusion
Viral load (cps/ml)
Days after Ab infusion
IC50 Measure
Ab Data
PK Model Fit
VL Data
Total VL
V1
V2
Limit of Detection

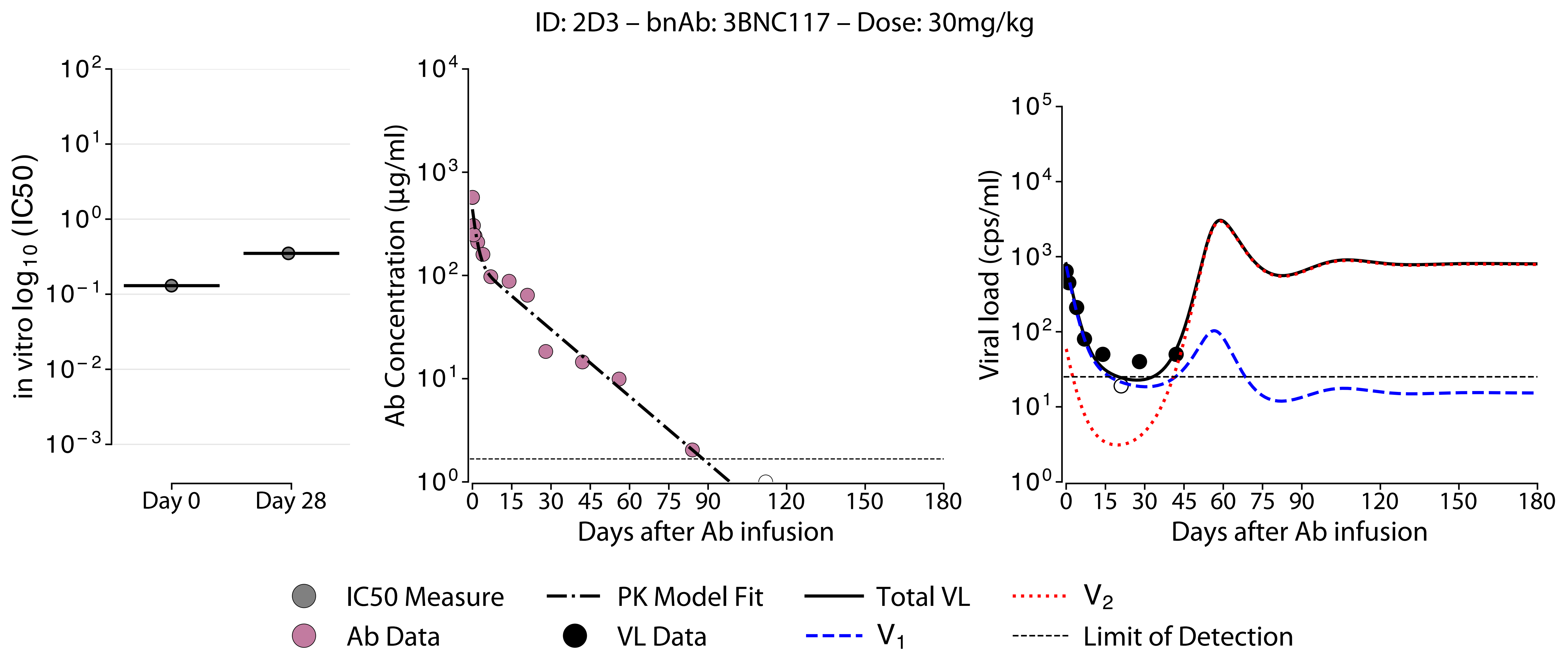
ID: 2D3 – bnAb: 3BNC117 – Dose: 30mg/kg
in vitro log10 (IC50)
Day 0
Day 28
Ab Concentration (µg/ml)
Days after Ab infusion
Viral load (cps/ml)
Days after Ab infusion
IC50 Measure
PK Model Fit
Total VL
V2
Ab Data
VL Data
V1
Limit of Detection

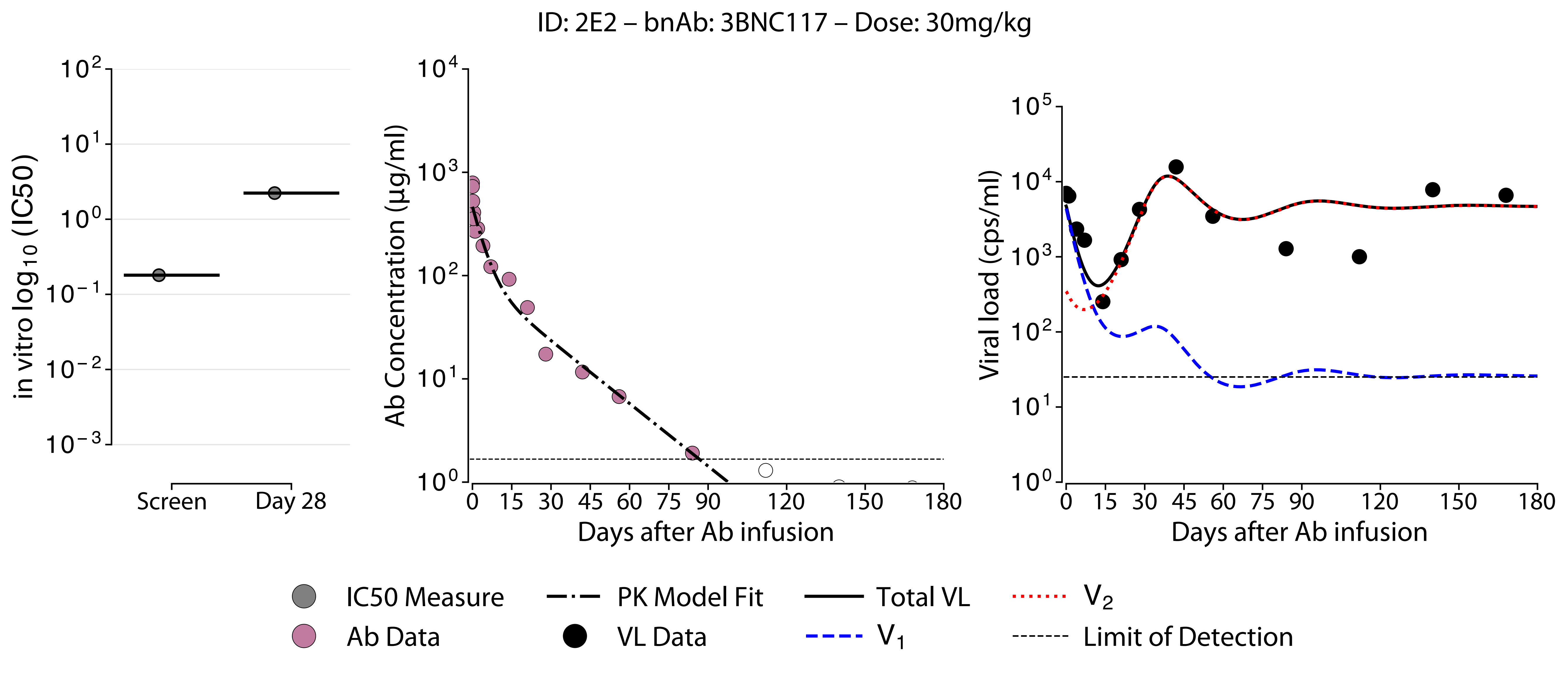

ID: 2E2 – bnAb: 3BNC117 – Dose: 30mg/kg
in vitro log10 (IC50)
Screen
Day 28
Ab Concentration (μg/ml)
Days after Ab infusion
Viral load (cps/ml)
Days after Ab infusion
IC50 Measure
Ab Data
PK Model Fit
VL Data
Total VL
V1
V2
Limit of Detection

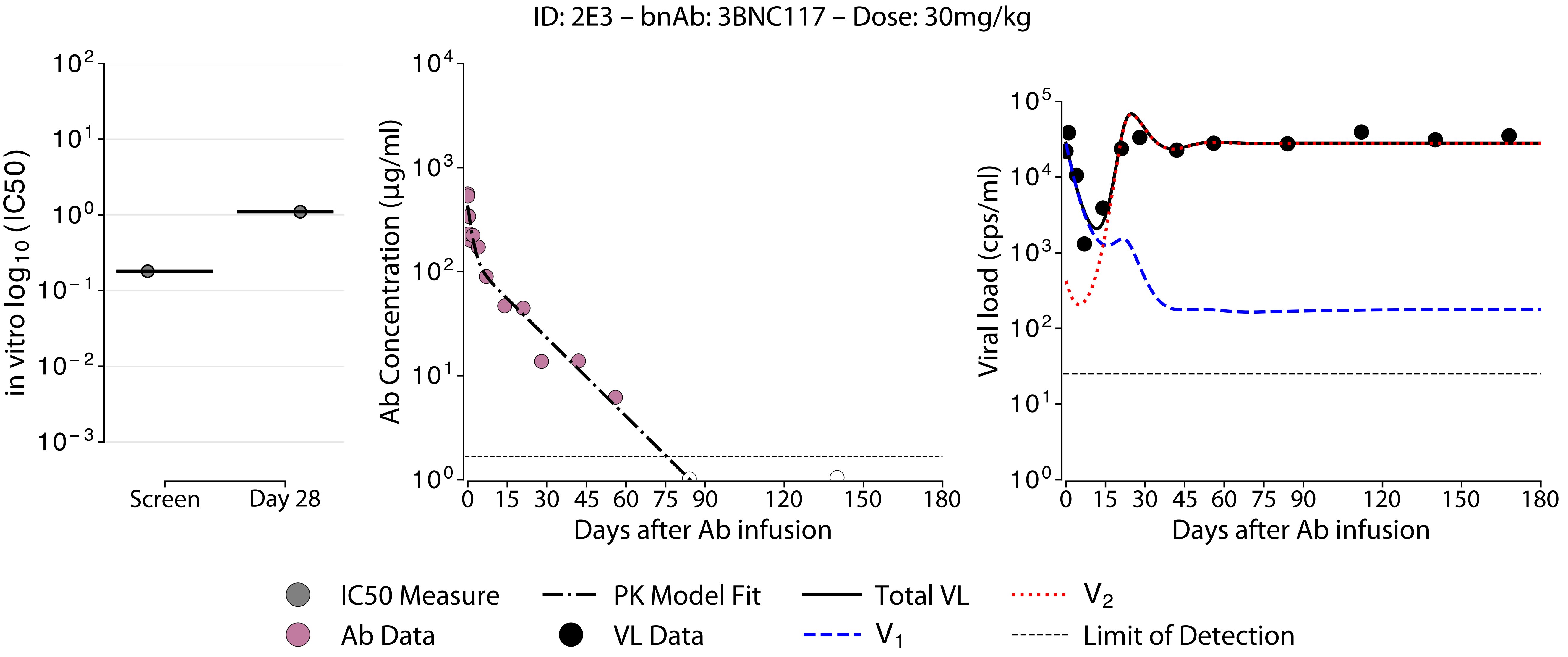
ID: 2E3 – bnAb: 3BNC117 – Dose: 30mg/kg
in vitro $\log_{10}$ (IC50)
$10^2$
$10^1$
$10^0$
$10^{-1}$
$10^{-2}$
$10^{-3}$
Screen
Day 28
Ab Concentration (μg/ml)
$10^4$
$10^3$
$10^2$
$10^1$
$10^0$
0 15 30 45 60 75 90 120 150 180
Days after Ab infusion
Viral load (cps/ml)
$10^5$
$10^4$
$10^3$
$10^2$
$10^1$
$10^0$
0 15 30 45 60 75 90 120 150 180
Days after Ab infusion
IC50 Measure
PK Model Fit
Total VL
$V_2$
Ab Data
VL Data
$V_1$
Limit of Detection

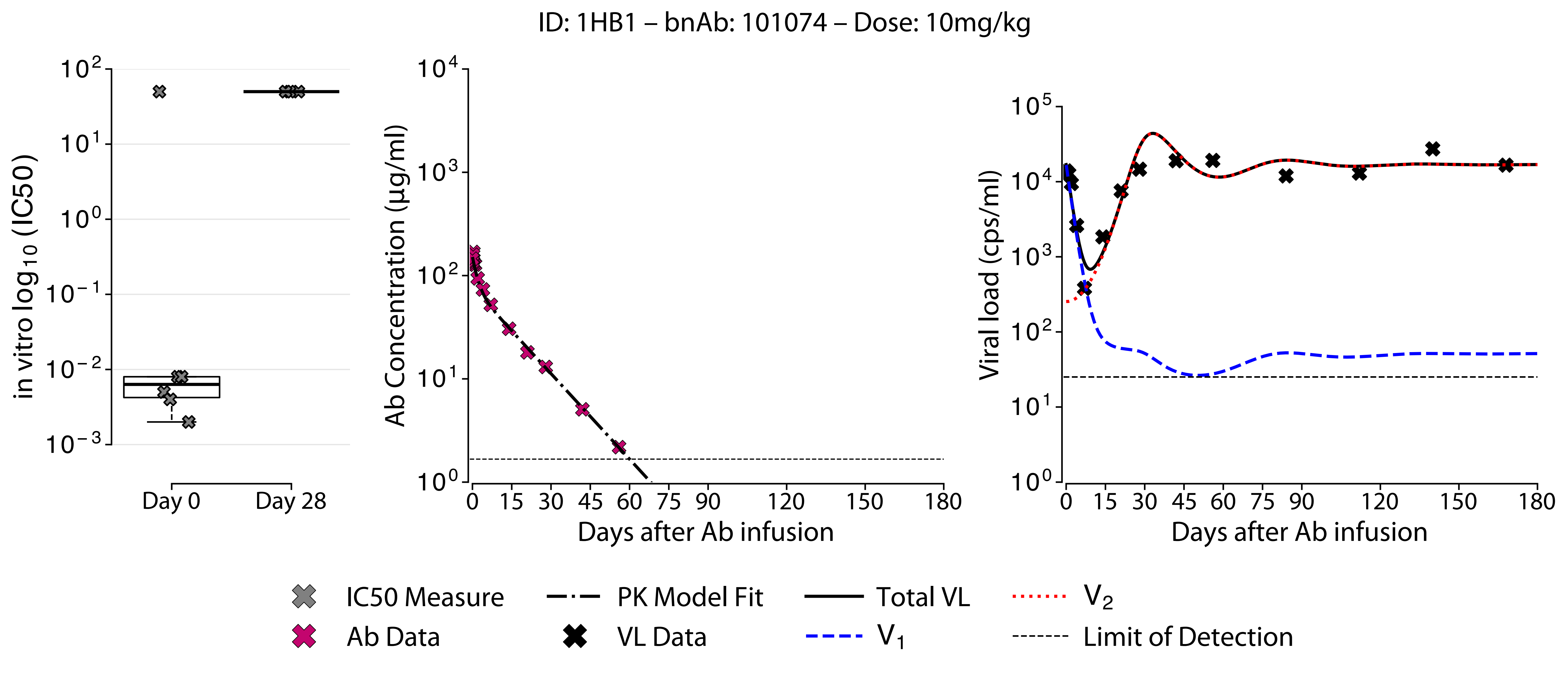

ID: 1HB1 – bnAb: 101074 – Dose: 10mg/kg
in vitro log10 (IC50)
Day 0
Day 28
Ab Concentration (µg/ml)
Days after Ab infusion
Viral load (cps/ml)
Days after Ab infusion
IC50 Measure
Ab Data
PK Model Fit
VL Data
Total VL
V1
V2
Limit of Detection

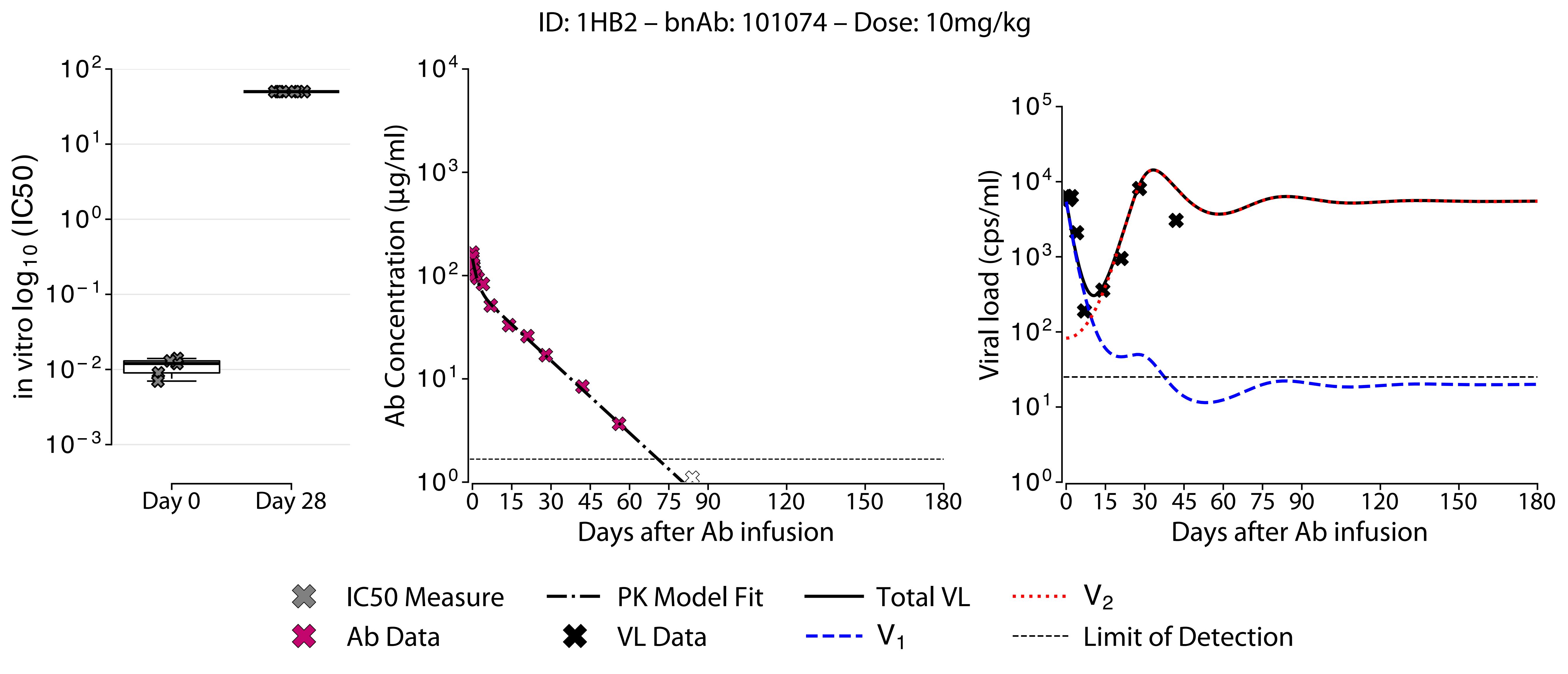

ID: 1HB2 – bnAb: 101074 – Dose: 10mg/kg
in vitro log10 (IC50)
Day 0
Day 28
Ab Concentration (µg/ml)
Days after Ab infusion
Viral load (cps/ml)
Days after Ab infusion
IC50 Measure
Ab Data
PK Model Fit
VL Data
Total VL
V1
V2
Limit of Detection

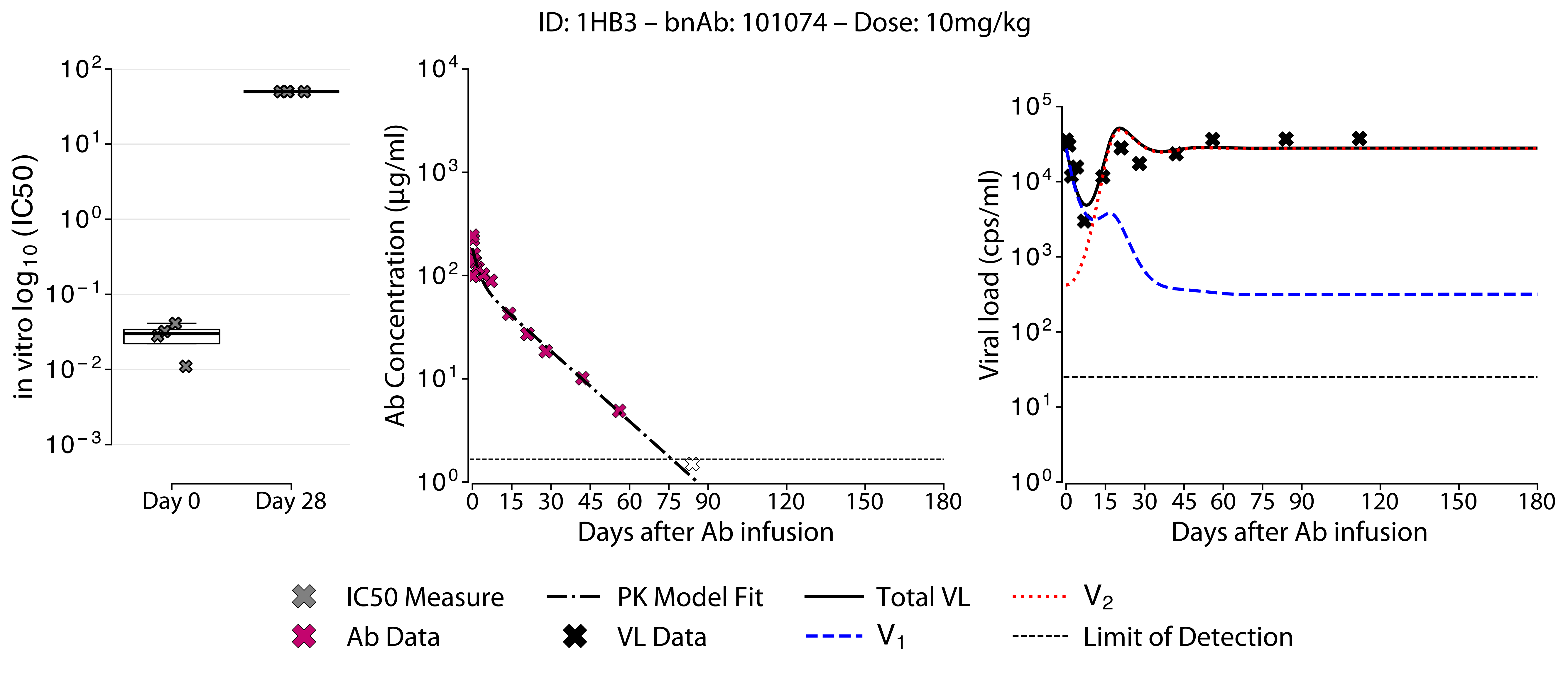

ID: 1HB3 – bnAb: 101074 – Dose: 10mg/kg
in vitro $\log_{10}$ (IC50)
$10^{2}$
$10^{1}$
$10^{0}$
$10^{-1}$
$10^{-2}$
$10^{-3}$
Day 0
Day 28
Ab Concentration (µg/ml)
$10^{4}$
$10^{3}$
$10^{2}$
$10^{1}$
$10^{0}$
0 15 30 45 60 75 90 120 150 180
Days after Ab infusion
Viral load (cps/ml)
$10^{5}$
$10^{4}$
$10^{3}$
$10^{2}$
$10^{1}$
$10^{0}$
0 15 30 45 60 75 90 120 150 180
Days after Ab infusion
IC50 Measure
Ab Data
PK Model Fit
VL Data
Total VL
$V_1$
$V_2$
Limit of Detection

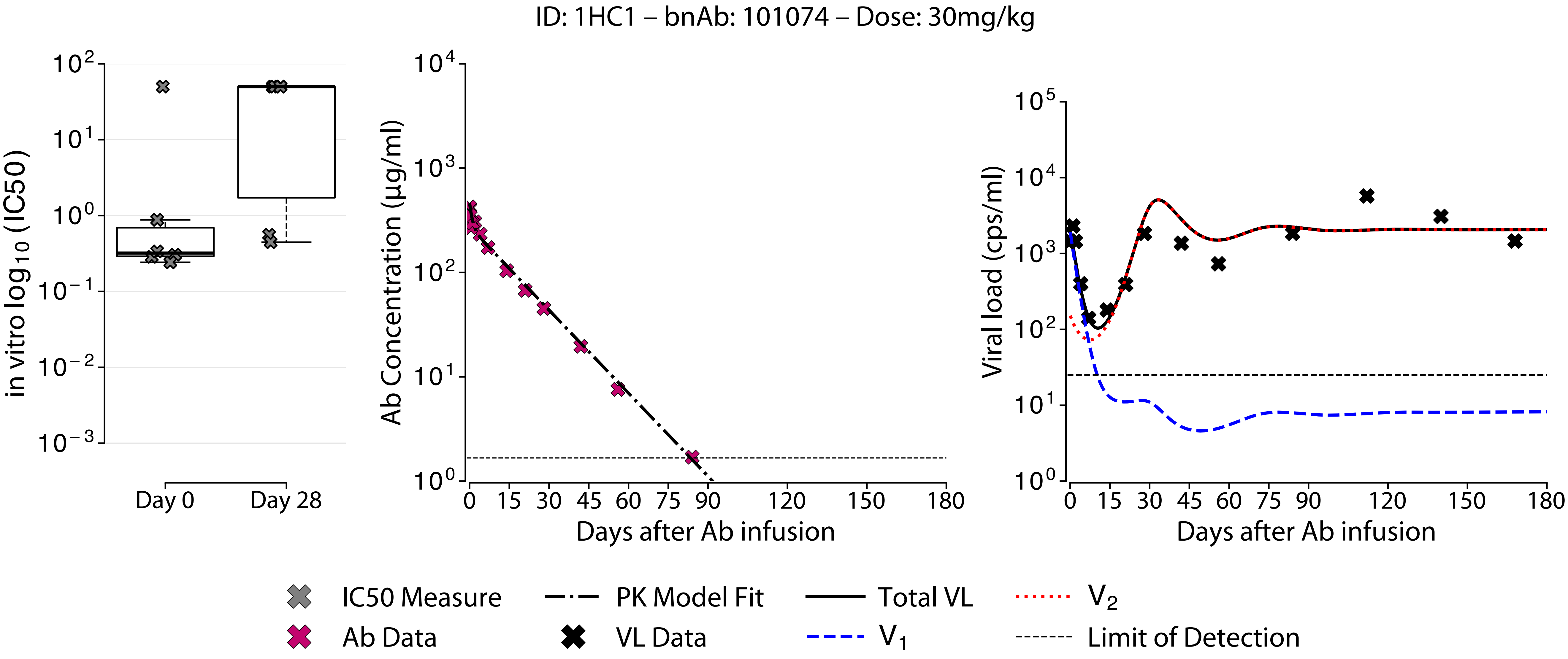

ID: 1HC1 – bnAb: 101074 – Dose: 30mg/kg
in vitro log10 (IC50)
Day 0
Day 28
Ab Concentration (µg/ml)
Days after Ab infusion
Viral load (cps/ml)
Days after Ab infusion
IC50 Measure
Ab Data
PK Model Fit
VL Data
Total VL
V1
V2
Limit of Detection

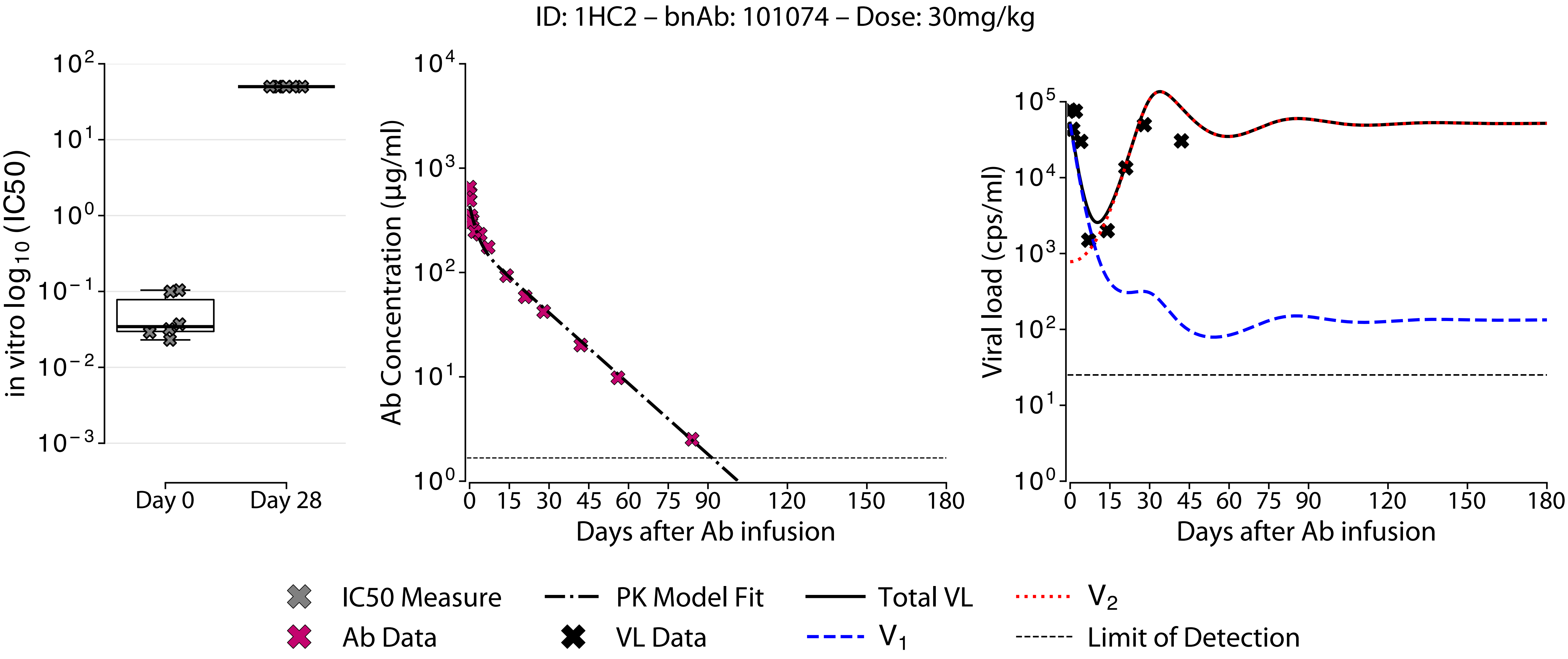
ID: 1HC2 – bnAb: 101074 – Dose: 30mg/kg
in vitro log10 (IC50)
Day 0
Day 28
Ab Concentration (µg/ml)
Days after Ab infusion
Viral load (cps/ml)
Days after Ab infusion
IC50 Measure
Ab Data
PK Model Fit
VL Data
Total VL
V1
V2
Limit of Detection

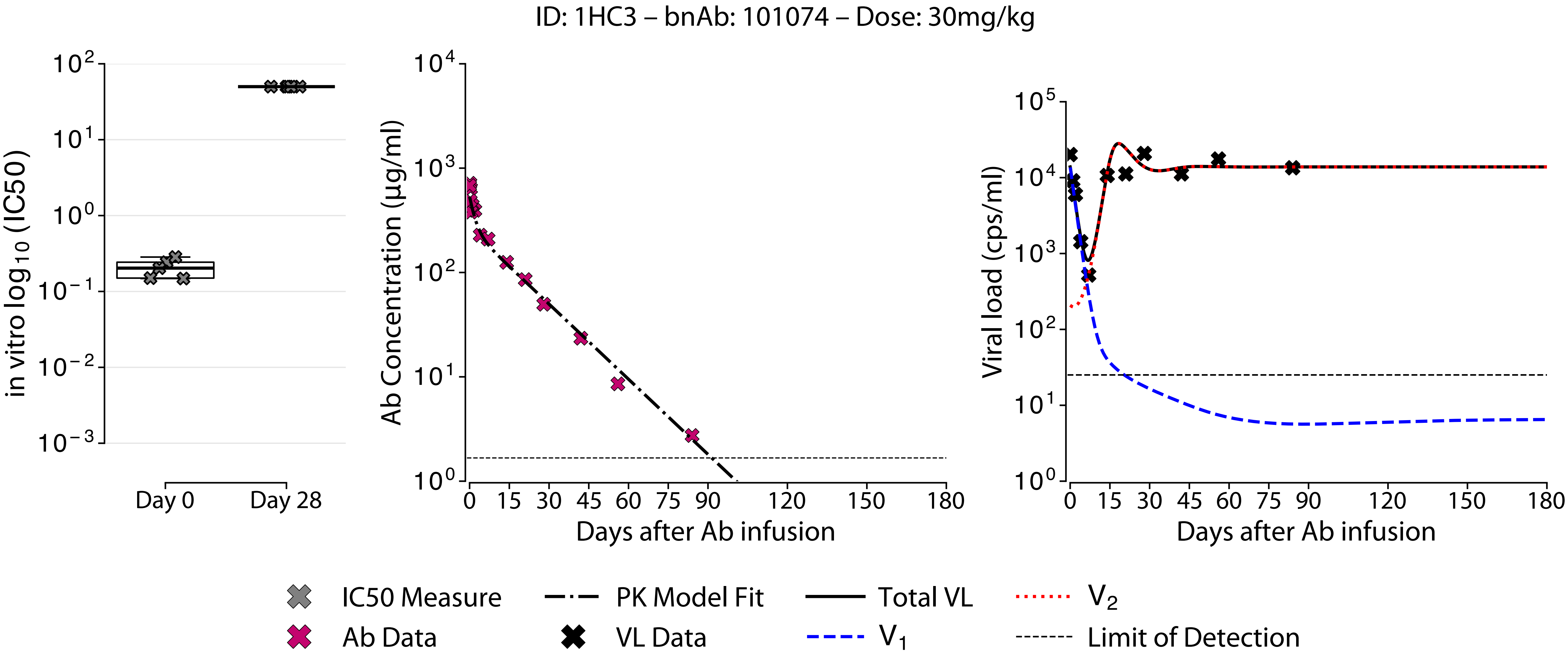
ID: 1HC3 – bnAb: 101074 – Dose: 30mg/kg
in vitro log10 (IC50)
Day 0
Day 28
Ab Concentration (µg/ml)
Days after Ab infusion
Viral load (cps/ml)
Days after Ab infusion
IC50 Measure
Ab Data
PK Model Fit
VL Data
Total VL
V1
V2
Limit of Detection

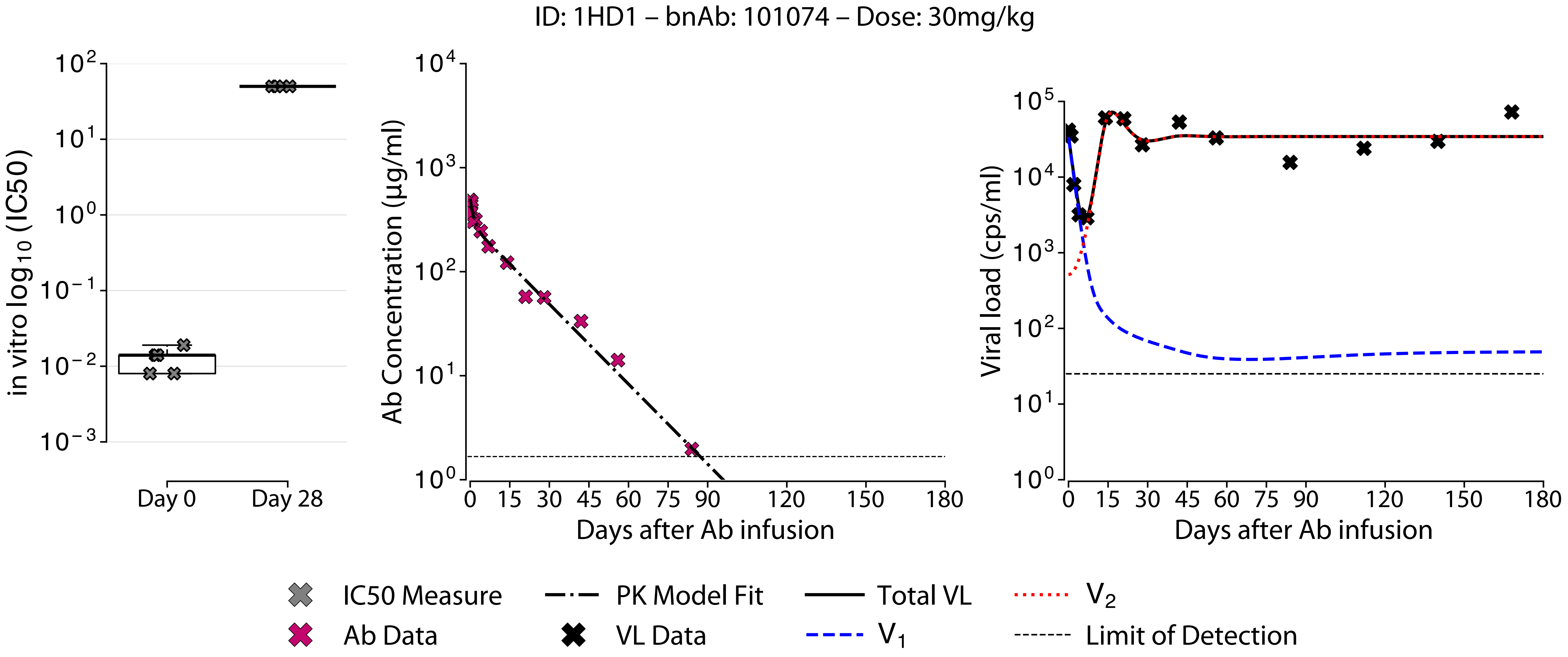

ID: 1HD1 – bnAb: 101074 – Dose: 30mg/kg
in vitro log10 (IC50)
Day 0
Day 28
Ab Concentration (µg/ml)
Days after Ab infusion
Viral load (cps/ml)
Days after Ab infusion
IC50 Measure
PK Model Fit
Total VL
V2
Ab Data
VL Data
V1
Limit of Detection

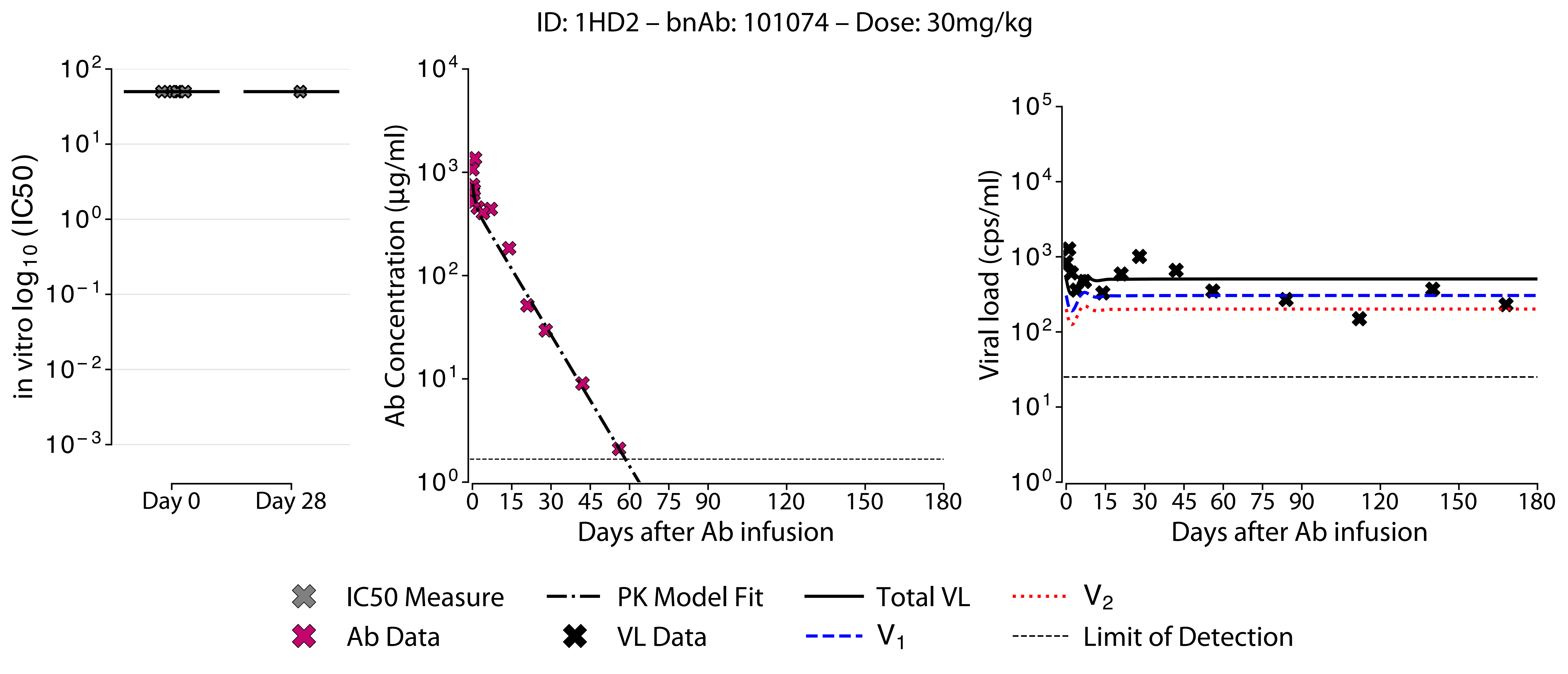
ID: 1HD2 – bnAb: 101074 – Dose: 30mg/kg
in vitro log10 (IC50)
Day 0
Day 28
Ab Concentration (μg/ml)
Days after Ab infusion
Viral load (cps/ml)
Days after Ab infusion
IC50 Measure
Ab Data
PK Model Fit
VL Data
Total VL
V1
V2
Limit of Detection

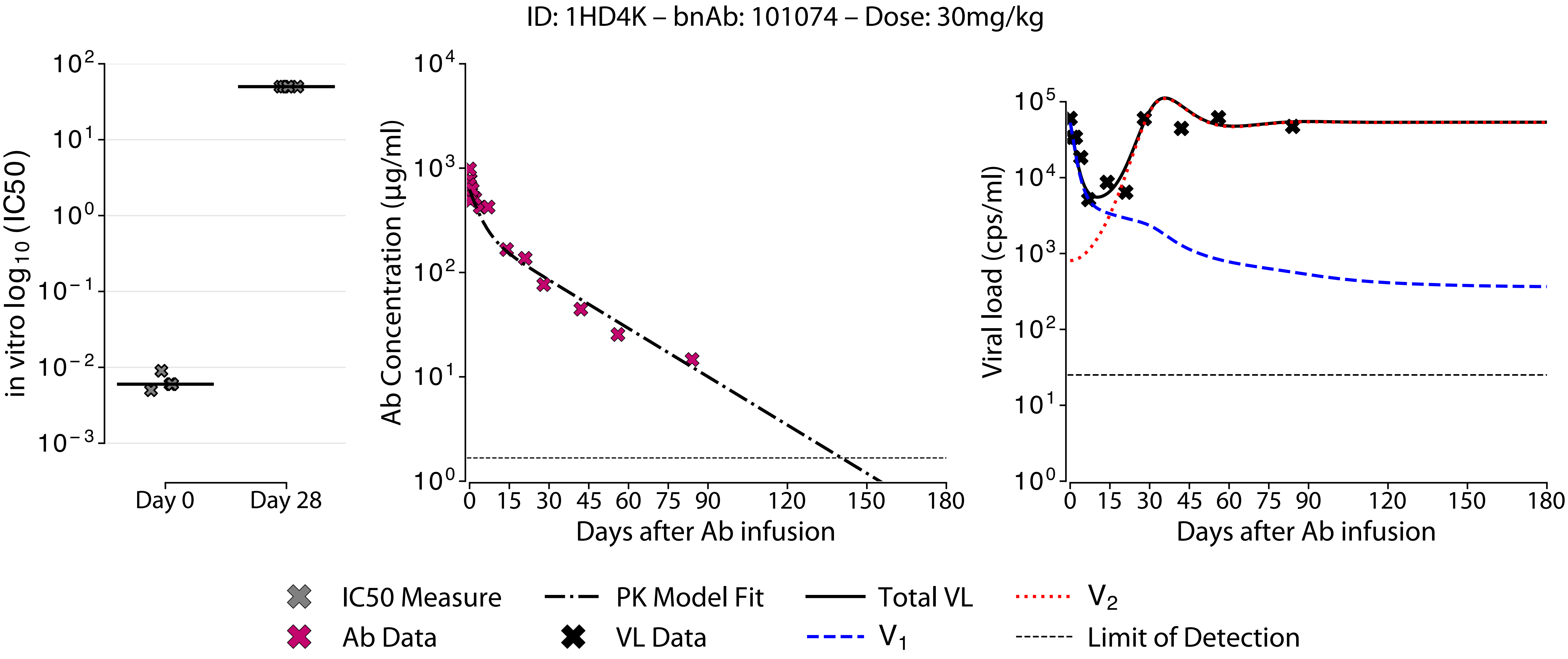
ID: 1HD4K – bnAb: 101074 – Dose: 30mg/kg
in vitro log10 (IC50)
$10^2$
$10^1$
$10^0$
$10^{-1}$
$10^{-2}$
$10^{-3}$
Day 0
Day 28
Ab Concentration (µg/ml)
$10^4$
$10^3$
$10^2$
$10^1$
$10^0$
0 15 30 45 60 75 90 120 150 180
Days after Ab infusion
Viral load (cps/ml)
$10^5$
$10^4$
$10^3$
$10^2$
$10^1$
$10^0$
0 15 30 45 60 75 90 120 150 180
Days after Ab infusion
IC50 Measure
Ab Data
PK Model Fit
VL Data
Total VL
$V_1$
$V_2$
Limit of Detection

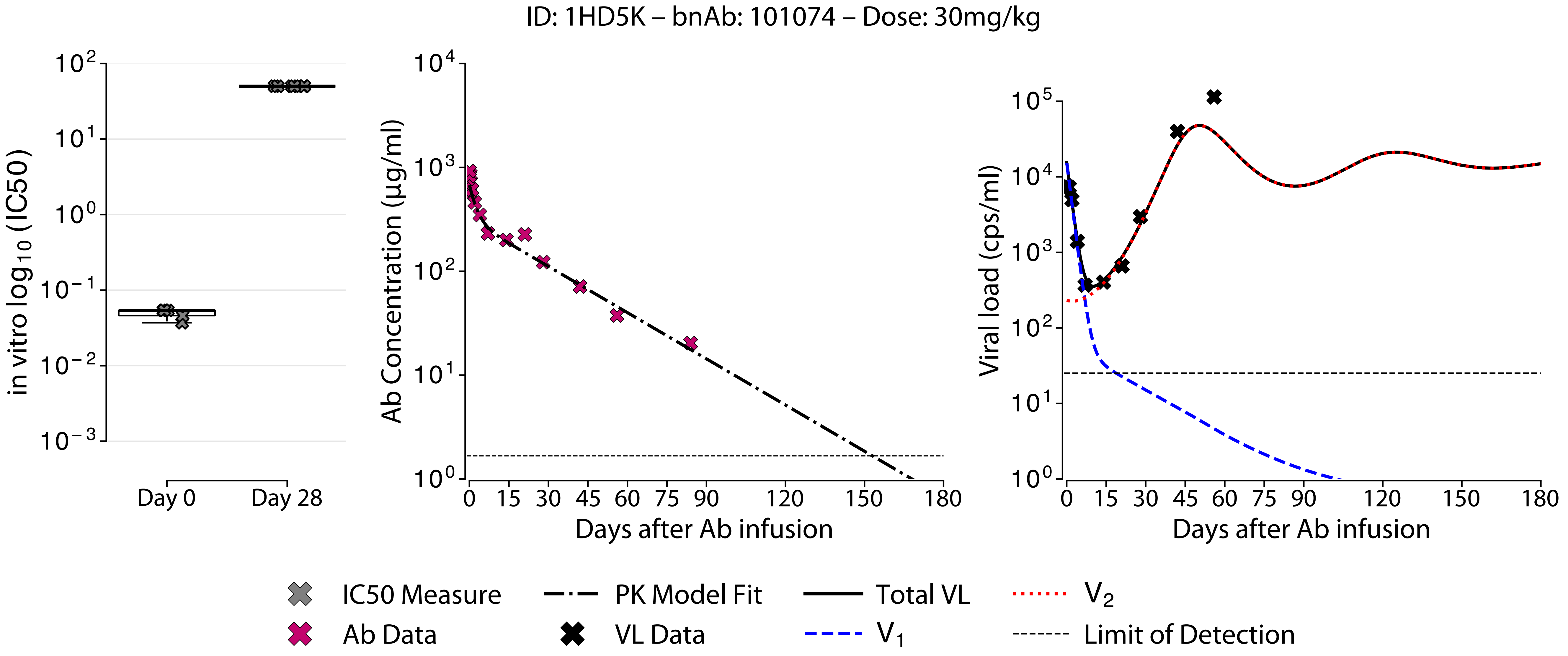

ID: 1HD5K – bnAb: 101074 – Dose: 30mg/kg
in vitro log10 (IC50)
Day 0
Day 28
Ab Concentration (μg/ml)
Days after Ab infusion
Viral load (cps/ml)
Days after Ab infusion
IC50 Measure
Ab Data
PK Model Fit
VL Data
Total VL
V1
V2
Limit of Detection

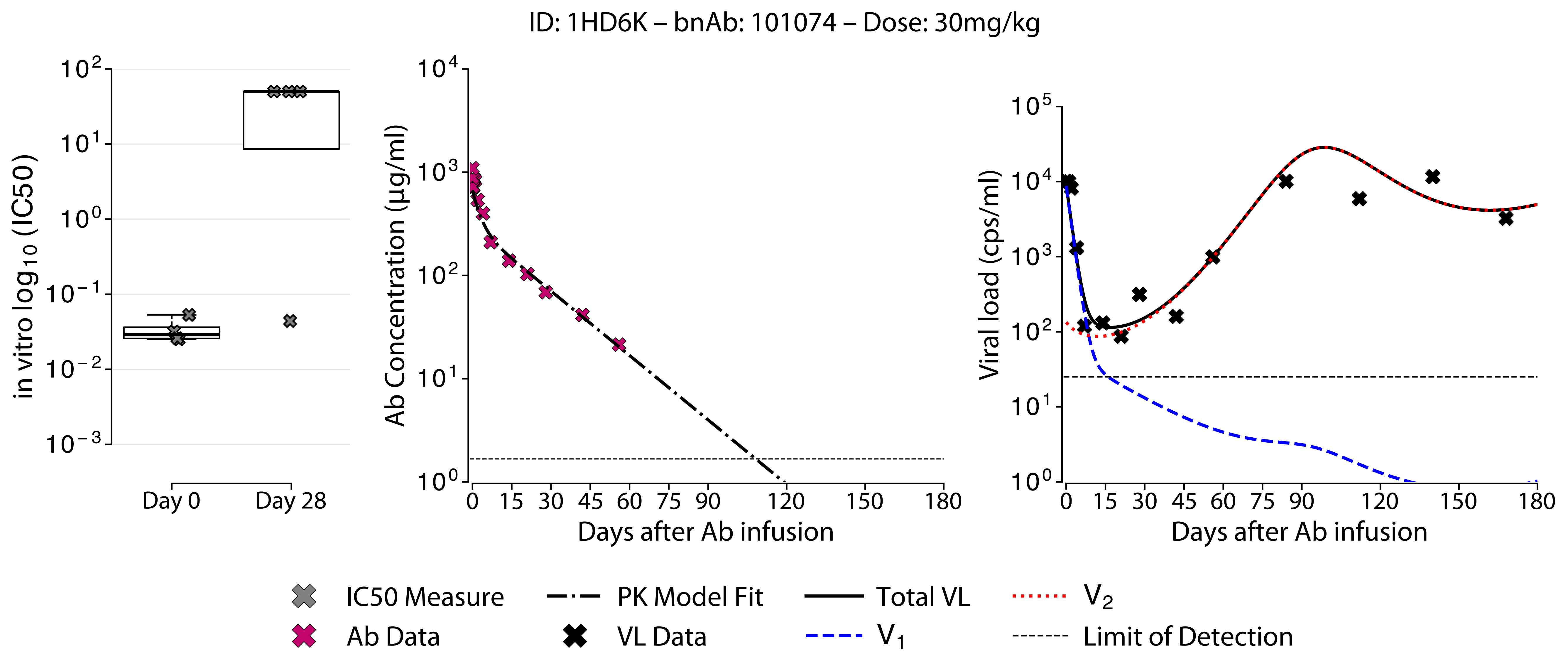

ID: 1HD6K – bnAb: 101074 – Dose: 30mg/kg
in vitro log10 (IC50)
Day 0
Day 28
Ab Concentration (μg/ml)
Days after Ab infusion
Viral load (cps/ml)
Days after Ab infusion
IC50 Measure
Ab Data
PK Model Fit
VL Data
Total VL
V1
V2
Limit of Detection

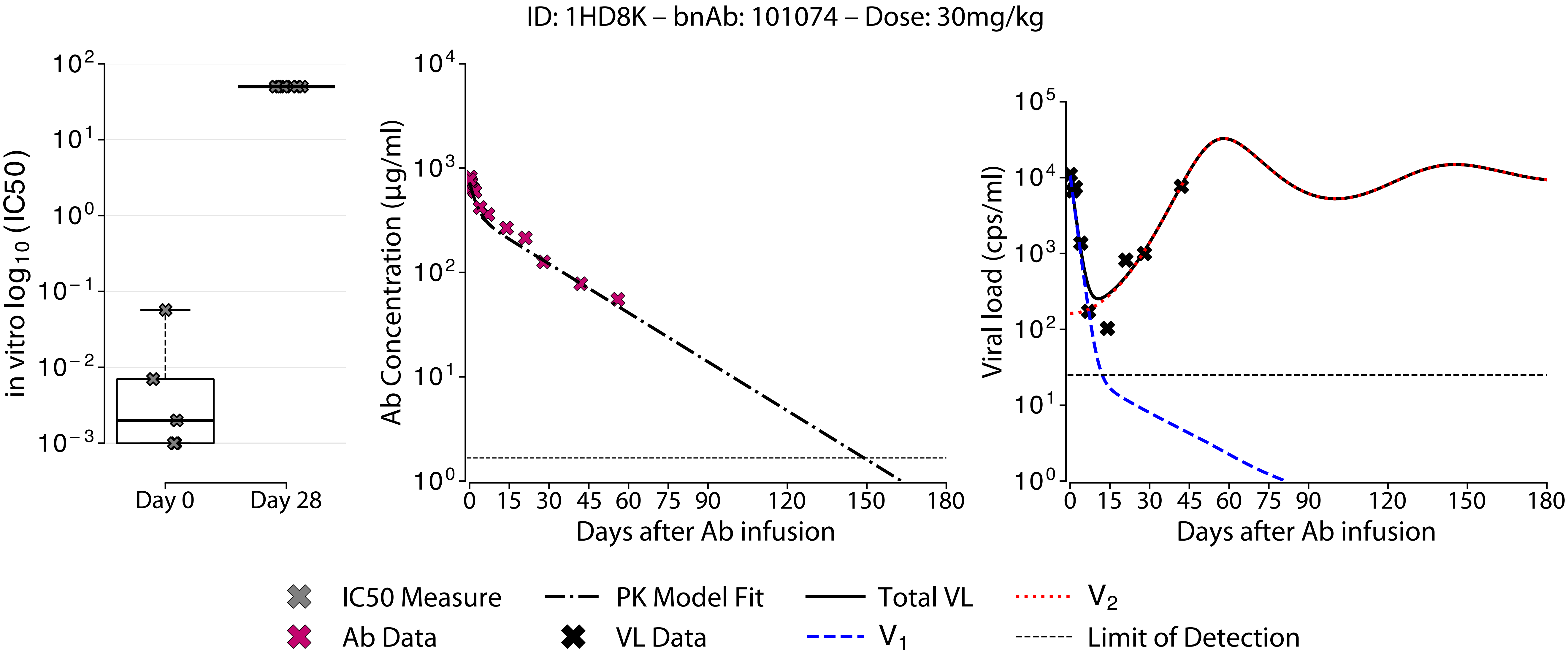
ID: 1HD8K – bnAb: 101074 – Dose: 30mg/kg
in vitro log10 (IC50)
Day 0
Day 28
Ab Concentration (µg/ml)
Days after Ab infusion
Viral load (cps/ml)
Days after Ab infusion
IC50 Measure
Ab Data
PK Model Fit
VL Data
Total VL
V1
V2
Limit of Detection

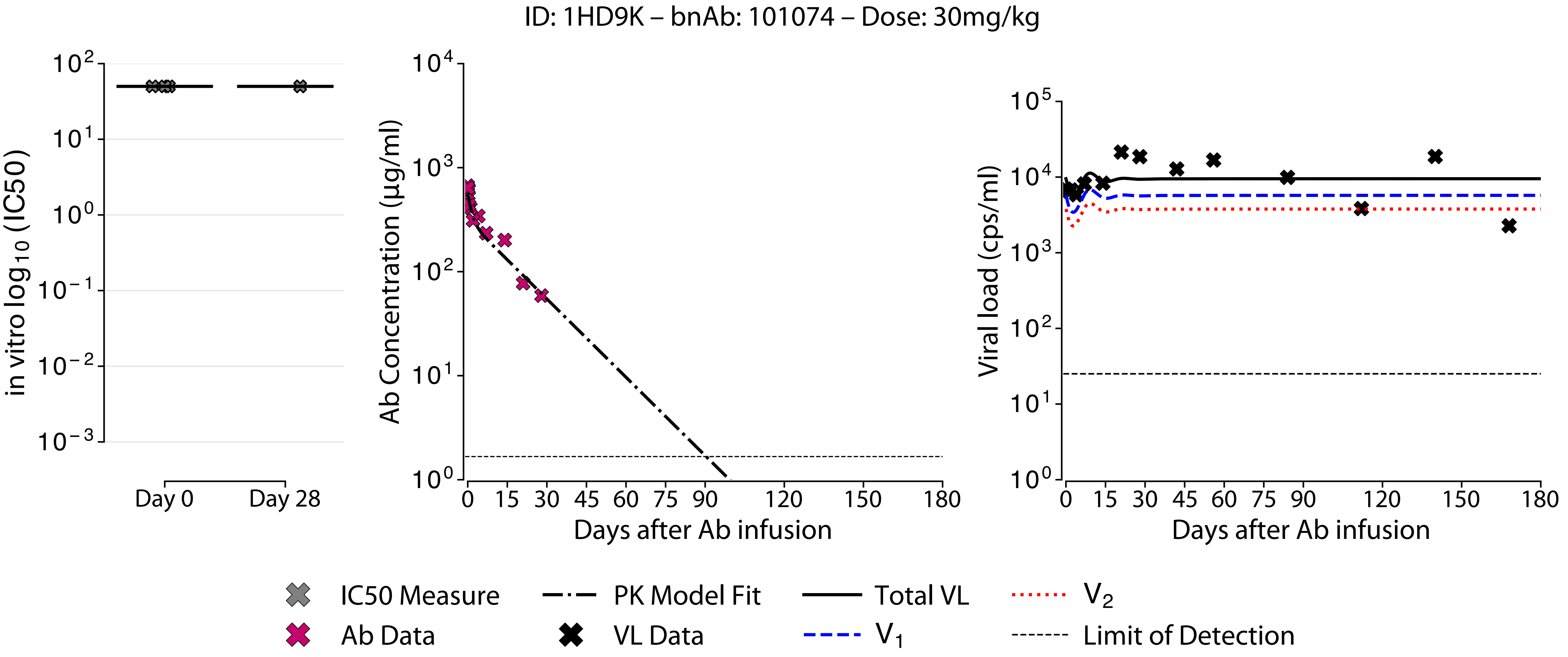

ID: 1HD9K – bnAb: 101074 – Dose: 30mg/kg
in vitro log10 (IC50)
Day 0
Day 28
Ab Concentration (µg/ml)
Days after Ab infusion
Viral load (cps/ml)
Days after Ab infusion
IC50 Measure
Ab Data
PK Model Fit
VL Data
Total VL
V1
V2
Limit of Detection
